# Functional and structural characterization of a combination of pan-sarbecovirus antibodies with potent antiviral activity

**DOI:** 10.1101/2025.03.30.646023

**Authors:** Grace C. Marden, Kimberly P. Schmitt, Maxwell I. Zimmerman, Alex H. Ramos, Monica Z. Menzenski, Geetika Sharma, Hongru Li, Eric Carlin, Anne Jecrois, Claudio L. Morales-Perez, Eugene Palovcak, Joseph D. Garlick, Brett Hannigan, Pamela Russell, Ross S. Federman, Alan Leung, Nicole Leon Vargas, Nadine M. Shaban, Vishvesha Vaidya, Hannah Towler, Nathan H. Joh, Dana M. Lord, Jenhan Tao, Zak Costello, Vincent Frappier, Kristen Hopson, Amarendra Pegu, Gavin C.K.W. Koh, Heather Van Epps, Adam Root, Gevorg Grigoryan, Daria Hazuda, Francesco Borriello

**Author notes:** These authors contributed equally (Grace C. Marden, Kimberly P. Schmitt, Maxwell I. Zimmerman, Alex H. Ramos).

## Abstract

Sarbecoviruses, a subgenus of coronaviruses, include strains with zoonotic spillover risk as exemplified by recent outbreaks (SARS-CoV-1, SARS-CoV-2). Monoclonal antibodies targeting conserved spike protein regions (RBD class 4, S2 fusion machinery) exhibit broad sarbecovirus neutralization, but their utility has been impacted by immune selection leading to escape and suboptimal neutralization. Using structure-conditioned machine-learning protein design we optimized two broadly neutralizing sarbecovirus antibodies to rescue the activity of a class 4 anti-RBD antibody against Omicron variants and improve the neutralization profile of an anti-S2 stem antibody, resulting in the identification of the lead candidates PRO-37587 and GB-0669. Both antibodies displayed remarkable resilience to SARS-CoV-2 evolution. Structural analyses elucidated the mechanistic basis of their broad neutralization profiles. In combination, these antibodies exhibited improved SARS-CoV-2 neutralization *in vitro* and *in vivo* and a higher barrier to resistance. These findings support further evaluation of such antibody combination as countermeasure against current and emerging sarbecoviruses.

## INTRODUCTION

4In recent decades, zoonotic spillover events involving the sarbecoviruses SARS-CoV-1 and SARS-CoV-2 have triggered major outbreaks in 2003 and 2019.^1–7^ The SARS-CoV-1 outbreak was effectively contained in 2003, with no additional cases reported since then.^8^ Conversely, SARS-CoV-2 caused a global pandemic that claimed millions of lives and is still circulating worldwide in humans as well as a variety of new animal reservoirs.^9^ The ongoing threat from SARS-CoV-2 infection is particularly significant for vulnerable populations, a risk further compounded by the evolution of immune escape variants harboring spike protein mutations that exhibit varying degrees of resistance to vaccine- and monoclonal antibody (mAb)-mediated immunity.^10–13^ Several sarbecoviruses also circulate in bats, posing a risk of future zoonotic spillovers, with evidence suggesting that such events may already be occurring in human communities that interact with wildlife.^14–16^ Therefore, it remains crucial to develop therapeutic measures that effectively control SARS-CoV-2 and prevent potential zoonotic spillover events of emerging sarbecoviruses, thereby averting future outbreaks and pandemics.

Several mAbs have been identified that exhibit broadly neutralizing activity against sarbecoviruses by targeting conserved regions of the spike protein, including the receptor-binding domain (RBD) and the S2 fusion machinery.^13,17,18^ Structural analysis of RBD in complex with human neutralizing antibodies has identified four major antigenic sites.^19^ Two of these sites (class 3 and 4) are targeted by most mAbs with broad neutralizing activity against sarbecoviruses.^13^ However, mutations in the RBD of several SARS-CoV-2 variants have led to escape and impaired neutralization by clinical and preclinical anti-RBD mAbs, limiting their therapeutic use.^10,13^ Neutralizing antibodies targeting the S2 fusion machinery typically bind the fusion peptide or the stem helix.^18^ These regions are under limited evolutionary pressure due to the lack of substantial population-level immunity directed against them, and therefore have not accrued escape mutations in SARS-CoV-2 variants. However, antibodies targeting the S2 fusion machinery generally have relatively low neutralization potency compared to anti-RBD antibodies, limiting their therapeutic utility.

Directed protein evolution is a powerful method for optimizing protein functions,^20^ and it has been used to optimize clinical-stage mAbs targeting SARS-CoV-2 RBD.^21^ However, application of this technique is challenging in settings where multiple molecular parameters need to be optimized simultaneously, including developability characteristics, especially when no straightforward selectable proxy trait exists. Therefore, to identify broadly neutralizing antibodies with appropriate characteristics for clinical development, we used a structure-conditioned machine learning (ML) protein design approach, leveraging integrated wet- and dry-lab workflows. Our approach uses a prior generative model to suggest modifications to a starting sequence in the first design round, with subsequent rounds using data from experimentally assessed variants to climb the landscape towards more potent sequences. In our experience, this approach generalizes well across problem settings, enabling robust optimization over any number of measurable parameters simultaneously, making it highly suited for therapeutic property and function tuning. In a single-parameter optimization setting, a related method recently showed success on improving antibody affinity.^22^

We selected the preclinical broadly neutralizing mAbs S2X259^23^ and CV3-25^24–26^, which respectively target the RBD class 4 region and the S2 stem helix, as starting points for our optimization campaigns. S2X259, isolated from a convalescent donor, inhibits the RBD-ACE2 interaction. This antibody exhibited neutralizing activity against SARS-CoV-2 pre-Omicron variants but lost this activity against most Omicron variants.^27,28^ CV3-25, also isolated from a convalescent donor, inhibits infection by impairing S2 refolding and fusion of SARS-CoV-2 with target cells. While CV3-25 retained neutralizing activity against SARS-CoV-2 Omicron variants, its neutralization potency and efficacy were suboptimal.^25^ By combining structure-conditioned ML protein design with high-throughput experimental assessment of spike binding and pseudovirus neutralization, we performed multi-parameter optimization to rescue the neutralization activity of S2X259 against SARS-CoV-2 Omicron variants without impairing its binding to other sarbecovirus RBDs. We also improved neutralization efficacy and potency of CV3-25. These efforts resulted in the identification of the lead candidates PRO-37587 and GB-0669. The structural basis for these antibodies’ unique breadth was elucidated, revealing a mechanism of action that may not have been achievable solely through rational engineering, thus highlighting the importance of multi-parameter optimization. Using Cryo-EM, we generated one of the few high-resolution structures of a SARS-CoV-2 Omicron lineage spike protein in an open conformation. This structure details a conformational rearrangement in the RBD epitope that explains both the well documented viral escape of SARS-CoV-2 Omicron lineages and the ability of PRO-37587 to rescue neutralization. Additionally, molecular dynamics simulations suggest that GB-0669 is optimized through allosteric stabilization of its apo ensemble. Notably, the combination of these optimized mAbs improved neutralization of SARS-CoV-2 variants *in vitro*, increased the barrier to resistance in an *in vitro* escape experiment with SARS-CoV-2 BA.1 live virus, and protected from SARS-CoV-2 infection in *in vivo* challenge models.

Overall, these findings demonstrate that enhanced antiviral activity is achieved by combining antibodies targeting distinct spike domains, each employing a different mechanism of action, thereby establishing the combination of PRO-37587 and GB-0669 as a promising strategy for sarbecovirus epidemic and pandemic response and preparedness.

## RESULTS

### ML-guided optimization of anti-S2 stem helix antibodies leads to improved neutralization potency

The S2 domain of SARS-CoV-2 spike contains highly conserved epitopes, namely the fusion peptide and the stem helix peptide. While neutralizing antibodies targeting these epitopes have been isolated from convalescent/vaccinated donors and immunized animals, they often are of low potency and not optimal for clinical development.^18^ Among published human S2-binding mAbs, the anti-S2 stem helix mAb CV3-25 emerged as a promising starting point for optimization due to its relatively higher neutralization potency against SARS-CoV-2 variants compared to other anti-S2 stem helix mAbs.^24–26^ CV3-25 binds to the spike protein in its pre-fusion state and blocks spike refolding and viral fusion with the target cells upon engagement with the ACE2 receptor.^24–26,29^

To explore the sequence space of the complementarity-determining regions (CDRs) and identify binders with improved S2 stem helix interaction, viral neutralization, and developability, we used a complex structure of CV3-25 Fab bound to the S2 stem helix (PDB: 7NAB). In the first round of design, we sampled 182 CDR variants from the sequence landscape compatible with the bound state using a structure-conditioned sequence generative model conditioned on the complex. The resulting sequences were expressed as human IgG1 with LS mutation in the Fc region to enhance half-life^30^ and assessed for developability (self-association, monomericity, polyspecific reactivity [data not shown]), binding to recombinant spike proteins (data not shown) and function (pseudovirus neutralization [SARS-CoV-2 Delta and BA.2]). With these data, we trained local sequence-to-function (seq2func) models that predicted cross-reactive potency and affinity from sequence. Finally, in the second design round, seq2func models were co-optimized with the main objective of the generative model, to produce 364 designs that were also assessed for developability, binding and pseudovirus neutralization (**Figures S1A and S1B**).

Experimental evaluation of round 1 showed that 26 designs (14%) exhibited improved SARS-CoV-2 Delta neutralization, and 13 designs (7%) demonstrated enhanced SARS-CoV-2 BA.2 neutralization compared to the reference CV3-25. In round 2, the fraction of designs with improved neutralization increased significantly, with 134 designs (36%) showing better SARS-CoV-2 Delta neutralization and 120 designs (33%) displaying improved SARS-CoV-2 BA.2 neutralization. Notably, the designs with enhanced SARS-CoV-2 BA.2 neutralization harbored between 1 to 13 mutations in the CDRs, with a median distance of 6.0 mutations from the reference CV3-25 (**Figure S1C**). After experimental evaluation of all 533 unique sequences generated from the two rounds of sequence design, GB-0669 was identified as lead candidate based on optimal functional, binding, and developability properties, and selected for further characterization (**Figure S1D**).

### GB-0669 binds to the S2 stem helix with higher affinity and displays improved neutralization potency against sarbecoviruses compared to CV3-25

A distinctive property of anti-S2 stem helix mAbs is their breadth of activity due to the conserved target region. To assess whether this was also true for GB-0669, we evaluated its binding and neutralization to a large panel of recombinant spike trimers and pseudoviruses representative of SARS-CoV-2 variants and non-SARS-CoV-2 sarbecoviruses. The reference mAb CV3-25 was used in the same set of experiments to quantify the extent of improvement. GB-0669 showed robust binding to all spike trimers as assessed by dissociation-enhanced lanthanide fluorescent immunoassay (DELFIA), and overall lower half maximal effective concentration (EC50) values compared to CV3-25, hence confirming improved binding (**Figure 1A**). These results were confirmed by surface plasmon resonance (SPR), with GB-0669 Fab showing ∼5-fold higher binding affinity to selected SARS-CoV-2 spike trimers compared to CV3-25 Fab (**Figure 1B**). No binding to MERS spike trimer was detected. Then, we evaluated GB-0669 neutralization with pseudoviruses representative of SARS-CoV-2 D614G (ancestral strain), Delta, Omicron BA.2, BA.4/5, BQ.1.1 and XBB.1.5 variants as well as the non-SARS-CoV-2 sarbecoviruses SARS-CoV-1 and WIV1. GB-0669 showed robust neutralization of all pseudoviruses and improved neutralization potency and efficacy compared to the reference mAb CV3-25 (**Figure 1C, Table S1**). Overall, GB-0669 functional characterization demonstrates its breadth of activity and improvement over the reference mAb CV3-25.

**Figure 1.**
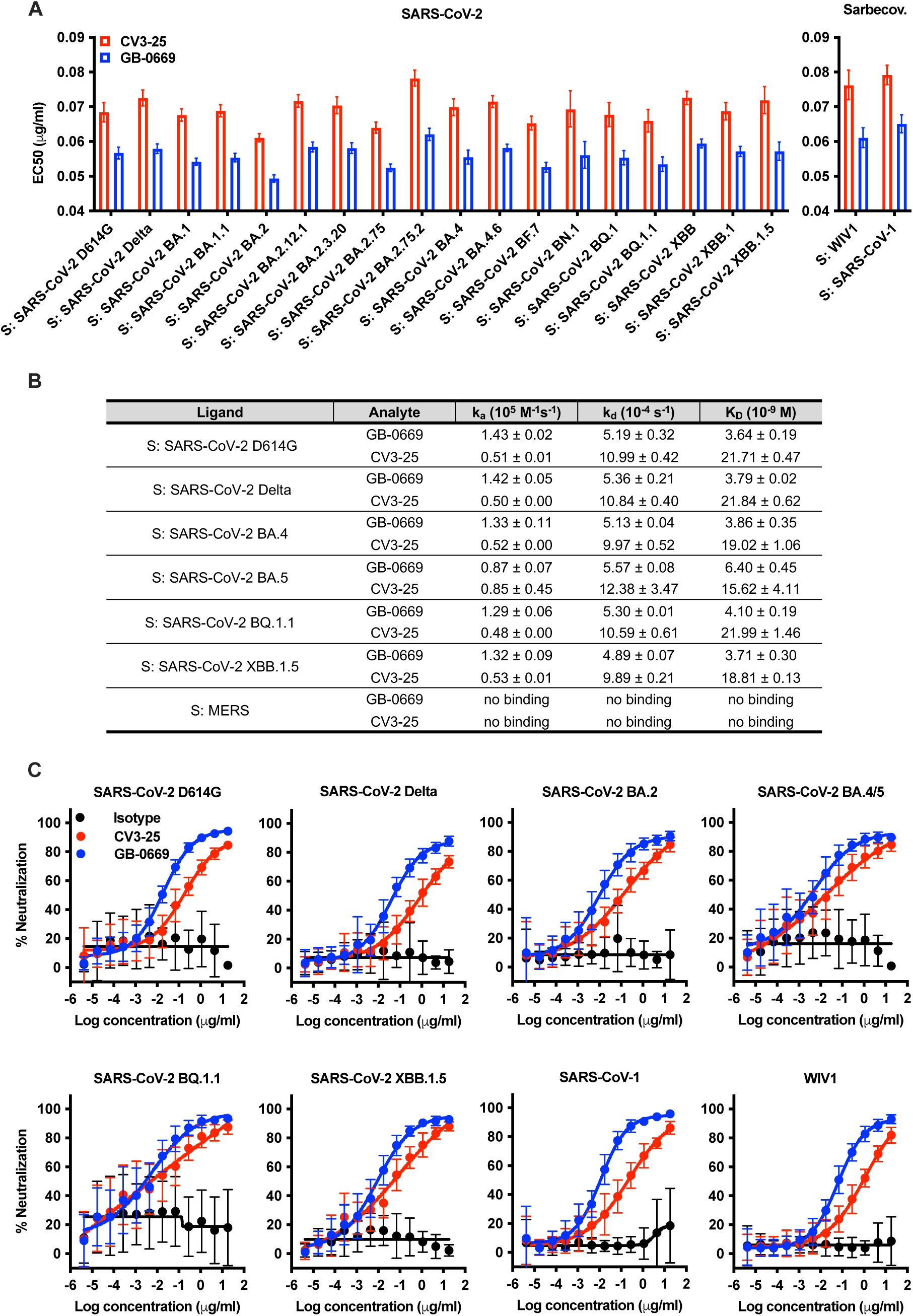
GB-0669 shows improved binding and neutralization profiles compared to the reference antibody CV3-25. (A) Binding of GB-0669 and CV3-25 to spike trimers (indicated as “S:” followed by the respective virus name) representative of major SARS-CoV-2 variants (SARS-CoV-2) and non-SARS-CoV-2 sarbecoviruses (Sarbecov.). Results are expressed as EC50 values (µg/ml), shown as mean ± SD and representative of two independent experiments, four technical replicates each. (B) Kinetics of GB-0669 and CV3-25 Fabs binding to a panel of SARS-CoV-2 spike trimers and negative control MERS spike trimer obtained by SPR at 25°C. The parameters displayed in the table represent the following: k_a_, association rate constant; k_d_, dissociation rate constant; K_D_, equilibrium dissociation constant. Results are expressed as mean ± SD of one experiment with two technical replicates. (C) Neutralization profiles of GB-0669, CV3-25 and isotype control mAb against pseudoviruses representative of SARS-CoV-2 variants (D614G, Delta, Omicron BA.2, BA.4/5, BQ.1.1 and XBB.1.5) and non-SARS-CoV-2 sarbecoviruses (SARS-CoV-1, WIV1). Results are reported as percentage (%) neutralization, shown as mean ± SD and representative of three to six independent experiments, four technical replicates each. See also Table S1.

To confirm binding mode of GB-0669, we initially performed a binding experiment with biotinylated peptides previously reported as part of CV3-25 characterization (partially overlapping peptides spanning the CV3-25 target region [amino acid 1133 - 1162], a peptide representing the C-terminal end of the stem helix [amino acid 1149 - 1167], a control 15mer peptide derived from HIV-1 Env protein).^25^ An additional biotinylated peptide was designed to encompass the SARS-CoV-2 spike S2 stem helix (amino acid 1143 - 1162). GB-0669 showed the same binding pattern as CV3-25 (**Figure 2A**). Of note, SPR analysis demonstrated that GB-0669 Fab binds to the S2 stem helix with higher affinity compared to CV3-25 Fab (**Figure 2B**).

**Figure 2.**
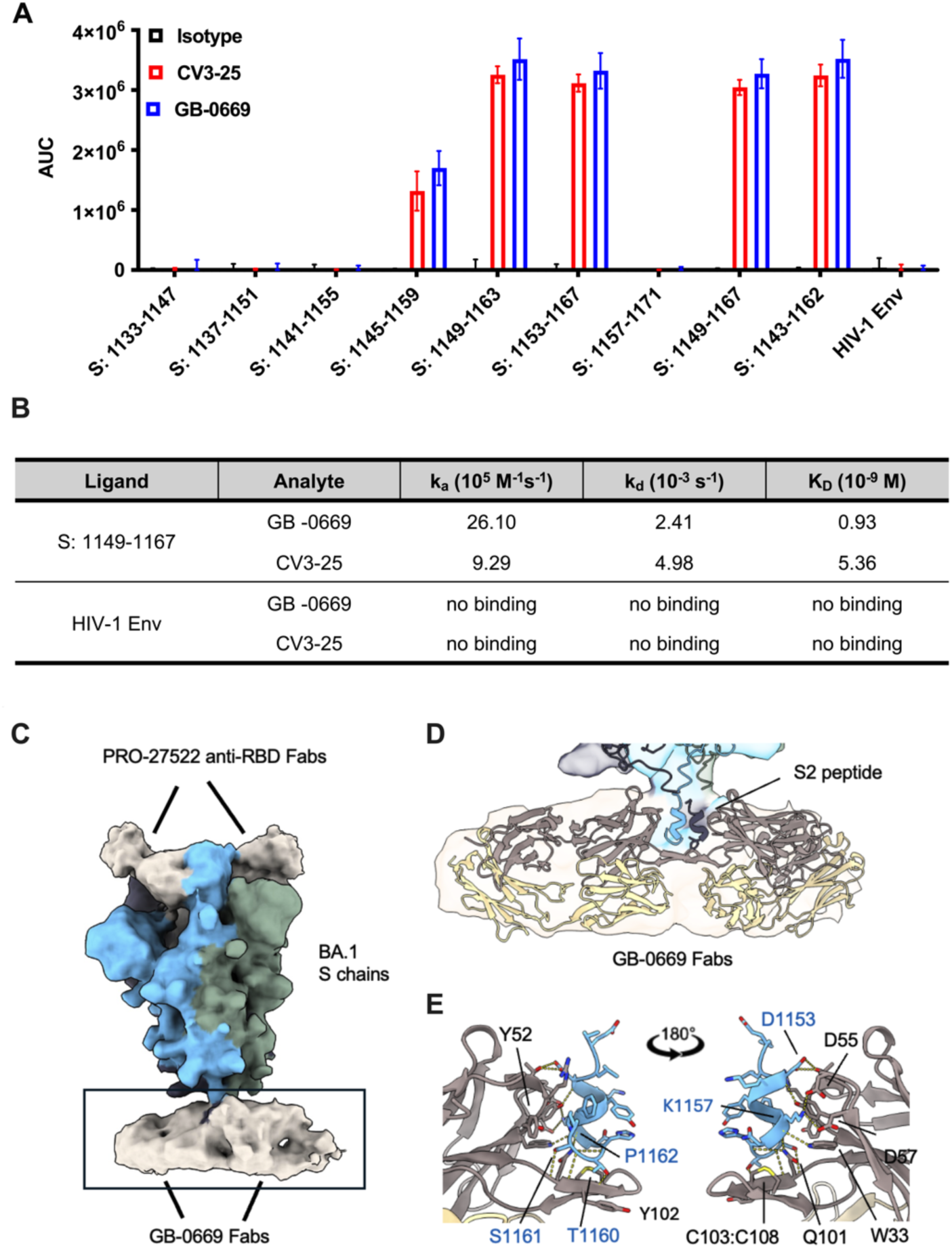
GB-0669 binds to the S2 stem helix with increased binding affinity compared to the reference antibody CV3-25. (A) Binding of GB-0669, CV3-25 and isotype control mAb to SARS-CoV-2 spike S2 peptides (indicated as “S:” followed by amino acid positions) and HIV-1 Env negative control peptide assessed by DELFIA. Results are expressed as area under the curve (AUC) values and shown as bars representing mean ± SD (representative of two independent experiments, three technical replicates each). (B) Kinetics of GB-0669 and CV3-25 Fabs binding to SARS-CoV-2 spike S2 stem helix peptide (indicated as “S:” followed by amino acid positions) and HIV-1 Env negative control peptide were obtained by SPR at 25°C. The parameters displayed in the table represent the following: k_a_, association rate constant; k_d_, dissociation rate constant; K_D_, equilibrium dissociation constant. Results are representative of one experiment without technical replicates. (C) Low-pass filtered Cryo-EM map (EMD-49461) of the SARS-CoV-2 Omicron BA.1 spike trimer bound to GB-0669 Fabs at the S2 stem helix and PRO-27522 Fabs at the RBDs. (D) Close-up on the density of the map in (C) with 2 copies of the GB-0669 S2 stem helix peptide fragment (PDB: 9NHP) docked into the map. (E) Atomic model of GB-0669 (PDB: 9NHP) heavy chain (grey structure and residue numbers) in complex with the S2 stem helix peptide (blue structure and residues numbers). Residues which make potentially stabilizing interactions on GB-0669 and the S2 stem helix peptide are labeled. Interface hydrogen bonds are drawn as dotted yellow lines.

### Multiple GB-0669 molecules engage the S2 stem helix in the SARS-CoV-2 Omicron spike

We performed structural analysis to investigate the interaction between GB-0669 Fab and the S2 stem helix epitope within the context of SARS-CoV-2 Omicron BA.1 spike. Single particle Cryo-EM experiments revealed that the interaction between GB-0669 Fab and S2 stem helix is inherently flexible, which initially made it challenging to identify bound molecules with high confidence. However, upon introducing an internally generated class 4 anti-RBD Fab (PRO-27522) we observed an improvement in map quality, allowing us to positively identify multiple bound Fab molecules of GB-0669 (**Figure 2C and 2D**). To gain further insight into the mechanistic details of the interaction, we solved the crystal structure at 1.93 Å resolution of GB-0669 Fab with the S2 stem helix peptide (PDB: 9NHP). GB-0669 Fab forms a network of hydrogen bonds with the S2 stem helix, consistent with what has been observed in CV3-25 (PDB: 7NAB). This extensive hydrogen bonding is primarily mediated by the heavy chain and involves residues from all three CDRs (**Figure 2E**).

Given the similarity of structures and contacting residues between GB-0669 and CV3-25 to the S2 stem helix, we hypothesized that differences in binding and neutralization may arise from differences in conformational ensembles. To assess this, we ran molecular dynamics simulations of each system, GB-0669 and CV3-25, in the apo form, and built Markov state models (MSMs) to characterize their ensembles.^31^ Projecting each MSM onto the root mean square deviation (RMSD) of CDR H3 compared to the native conformation reveals significant heterogeneity in both systems (**Figure S2**), where this loop has the ability to flip in an opposing direction. Though both systems have similar range of flexibility, CV3-25 populates these diverse states to a significantly higher degree compared with GB-0669. This result provides evidence that mutations help stabilize GB-0669 CDRs to favor the bound conformation, providing a mechanistic explanation to its improved binding and neutralization.

### ML-guided optimization of class 4 anti-RBD antibodies recovers neutralization of SARS-CoV-2 Omicron variants

Several antibodies targeting the RBD class 4 region have been reported since the beginning of the pandemic and some of them have demonstrated cross-reactivity within the sarbecovirus subgenus.^13^ However, mutations accrued in SARS-CoV-2 Omicron lineages have led to escape from this class of antibodies.^27,28^ We selected the class 4 anti-RBD mAb S2X259 as a starting point for optimization due to its breadth of activity.^23^ S2X259 binds to the RBD class 4 region and blocks RBD binding to ACE2 through steric hindrance. A complex structure of S2X259 Fab bound to RBD (PDB: 7M7W) was leveraged by our generative model. We conducted a five-round optimization campaign (with an average of ∼290 sequences tested per round) by using our structure-conditioned sequence generative model and seq2func models trained on experimental data, as indicated for the anti-S2 stem helix antibody campaign (**Figure S3A**). During the multiple optimization rounds, novel SARS-CoV-2 Omicron variants emerged, and the reference mAb S2X259 lost activity against such variants. Consequently, binding and neutralization experiments were updated from round to round to represent the most recent SARS-CoV-2 variants, ensuring that the multi-parameter optimization process would favor broadly neutralizing antibody sequences (**Figure S3B**). As optimization rounds progressed, top designs from each round exhibited progressively improved neutralization over S2X259 across five different SARS-CoV-2 variants, from Delta to Omicron BA.4/5 (**Figure S3C**). Experimental evaluation of round 5 showed a significant number of antibodies with robust neutralization of SARS-CoV-2 Omicron BA.4/5 (**Figure S3D**). The designs with enhanced SARS-CoV-2 Omicron BA.4/5 neutralization harbored between 9 to 17 mutations, with a median distance of 14 mutations relative to the reference S2X259 (**Figure S3E**). From this group, PRO-31391 was selected as candidate antibody (**Figure S3F**).

PRO-31391 showed potent neutralization of SARS-CoV-2 Omicron BA.4/5 pseudovirus even though the reference mAb S2X259 was completely inactive (**Figure S4A**). It did, however, retain a free cystine at light-chain position 51 from the reference mAb, which by virtue of being outside of the CDR region was not modified during optimization (**Figure S3F**). Since a free cysteine is a potential liability for developing a mAb due to the risk of self-aggregation or oxidation, we substituted it with an alanine, generating the molecule PRO-37587. Neutralization experiments demonstrated that this substitution had no impact on functional properties, as PRO-31391 and PRO-37587 had overlapping neutralization profiles (**Figure S4B**). Therefore, PRO-37587 was selected as lead candidate for further characterization based on optimal functional, binding, and developability properties.

### PRO-37587 has expanded breadth of binding and neutralization against sarbecoviruses compared to S2X259

Our class 4 anti-RBD campaign successfully demonstrated recovery of SARS-CoV-2 Omicron neutralization starting from a reference mAb which lost activity against these variants. Next, we asked whether this outcome came at the expense of cross-reactivity within the sarbecovirus subgenus. We leveraged yeast surface display to rapidly evaluate relative binding of PRO-37587 and the reference mAb S2X259 to RBDs representative of the sarbecovirus subgenus (SARS-CoV-2-related clade 1b, including SARS-CoV-2 variants, SARS-CoV-related clade 1a, Asia non-ACE2 bat sarbecoviruses clade 2, and Africa+Europe bat sarbecoviruses clade 3) as previously described.^32^ As expected, S2X259 showed poor binding to most of SARS-CoV-2 Omicron variants while PRO-37587 demonstrated binding to all RBDs representative of SARS-CoV-2-related clade 1b viruses (except for LC556375.1). Of note, S2X259 and PRO-37587 showed similar binding profiles to SARS-CoV-related clade 1 and Africa+Europe bat sarbecoviruses clade 3, and PRO-37587 also demonstrated expanded binding to RBDs representatives of Asia non-ACE2 bat sarbecoviruses clade 2 compared to S2X259 (**Figure 3A**). These results confirmed that our class 4 anti-RBD campaign led to recovery of SARS-CoV-2 Omicron reactivity while also preserving and in some cases expanding cross-reactivity within the sarbecovirus subgenus. To further substantiate these results, we assessed binding affinity of PRO-37587 and S2X259 mAbs to RBDs representative of SARS-CoV-2 D614G, Delta, Omicron BA.4/5, BQ.1.1 and XBB.1.5. Both PRO-37587 and S2X259 mAbs bound with high affinity to SARS-CoV-2 D614G and Delta RBDs. As expected, S2X259 showed lower binding affinity to SARS-CoV-2 Omicron BA.4/5, BQ.1.1 and XBB.1.5 RBDs, while PRO-37587 bound these RBDs with sub-nanomolar affinities (**Figure 3B**). These results mirrored pseudovirus neutralization data. S2X259 and PRO-37587 demonstrated comparable neutralization potencies against SARS-CoV-2 D614G and Delta, while only PRO-37587 neutralized SARS-CoV-2 Omicron variants (BA.2, BA.4/5, BQ.1.1, XBB.1.5). Both S2X259 and PRO-37587 showed neutralization of the non-SARS-CoV-2 sarbecoviruses SARS-CoV-1 and WIV1, with PRO-37587 showing a trend toward higher neutralization potencies compared to S2X259 (**Figure 3C, Table S2**). Overall, these results demonstrate that PRO-37587 exhibits a high degree of cross-reactivity within the sarbecovirus subgenus and significant improvement over the reference mAb S2X259.

**Figure 3.**
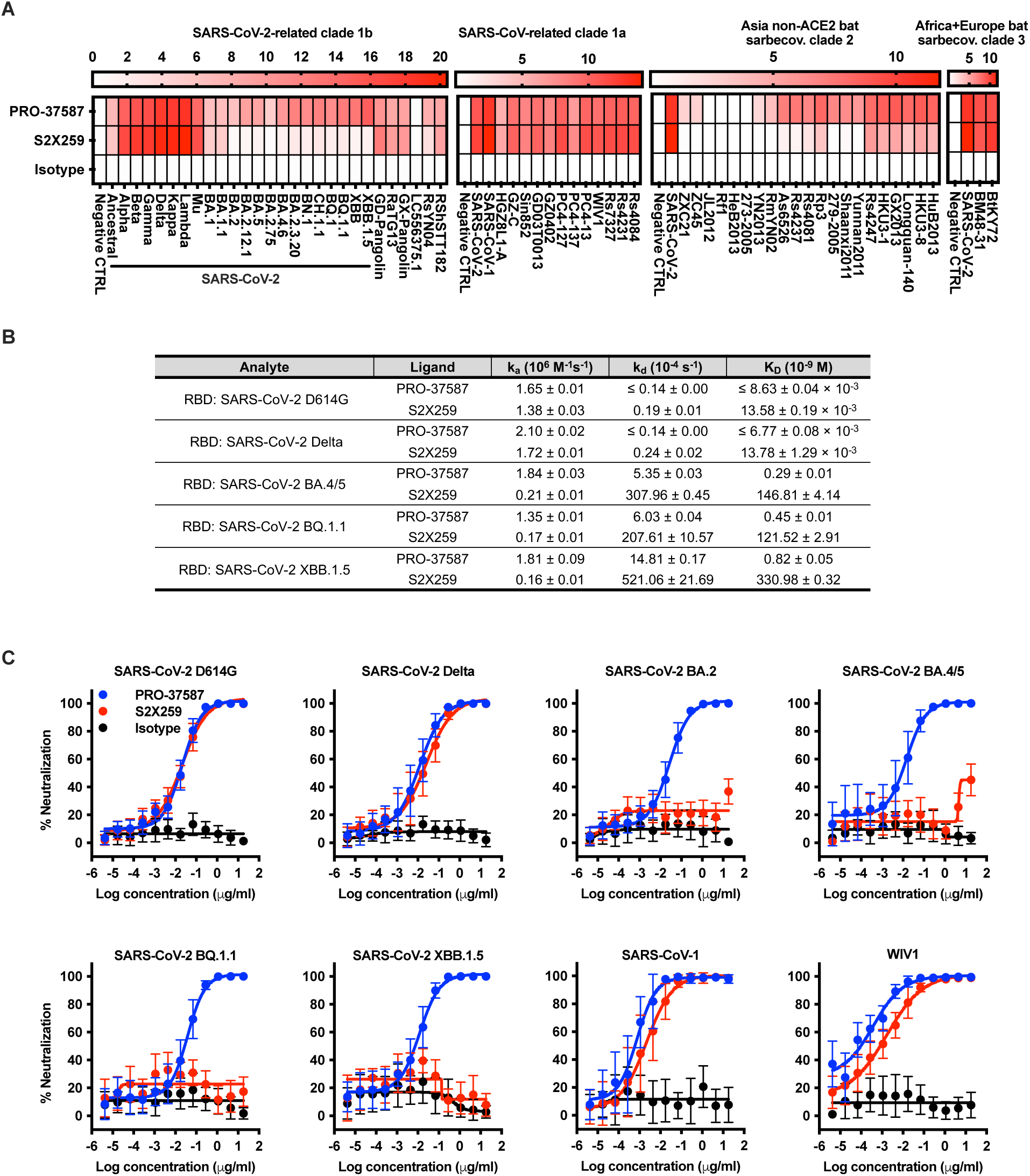
PRO-37587 shows improved binding and neutralization profiles compared to the reference antibody S2X259. (A) Flow cytometry binding of PRO-37587, S2X259 and isotype control mAb to RBDs representative of SARS-CoV-2-related clade 1b, SARS-CoV-related clade 1a, Asia non-ACE2 bat sarbecovirus (sarbecov.) clade 2, and Africa+Europe bat sarbecovirus clade 3 expressed on yeasts. Relative binding results are shown as median fluorescence intensity of the mAb signal divided by the median fluorescence intensity of the expression signal for each combination and reported as heatmaps relative to the mAb/RBD combination with the highest overall signal. (B) Kinetics of PRO-37587 and S2X259 mAbs binding to a panel of SARS-CoV-2 RBDs were obtained by SPR at 25°C. The parameters displayed in the table represent the following: k_a_, association rate constant; k_d_, dissociation rate constant; K_D_, equilibrium dissociation constant. Results are expressed as mean ± SD of one independent experiment with two technical replicates. (C) Neutralization profiles of PRO-37587, S2X259 and isotype control mAb against pseudoviruses representative of SARS-CoV-2 variants (D614G, Delta, Omicron BA.2, BA.4/5, BQ.1.1 and XBB.1.5) and non-SARS-CoV-2 sarbecoviruses (SARS-CoV-1, WIV1). Results are reported as percentage (%) neutralization and shown as mean ± SD (representative of three independent experiments, four technical replicates each). See also Table S2.

### PRO-37587 binding to the RBD class 4 region accommodates the mutations and RBD structure prevalent in SARS-CoV-2 Omicron lineages

Binding of class 4 anti-RBD Fabs requires a large-scale conformational change of the spike protein that reveals a cryptic site on the RBD.^33^ Broadly speaking, a cryptic site is a region that is not exposed in most conformational states but is still accessible to immune detection through conformational changes. Residues in this cryptic binding site are highly conserved across spike lineages, due to its partial overlap with the ACE2 binding site and the low probability of exposure. In total, this makes the site an appealing target for a broadly neutralizing antibody.

To gain mechanistic insights into the broad neutralization profile of PRO-37587, we generated a Cryo-EM map of PRO-37587 Fab bound to the full-length SARS-CoV-2 Omicron BA.1 spike protein (**Figure 4A**). As is well known, the spike protein open conformation poses challenges for obtaining a homogeneously high-resolution map. Indeed, 3D classification of the system revealed significant motion of the RBD upon PRO-37587 Fab binding (**Video S1**). To address this, we employed local refinement techniques focusing on PRO-37587 and the RBD, which enabled us to successfully achieve a homogeneously high-resolution map (2.97 Å) of the entire spike and anti-RBD Fab. To our knowledge, this represents the highest resolution map of a fully open SARS-CoV-2 Omicron BA.1 spike trimer with an anti-RBD Fab binding at the cryptic site.

**Figure 4.**
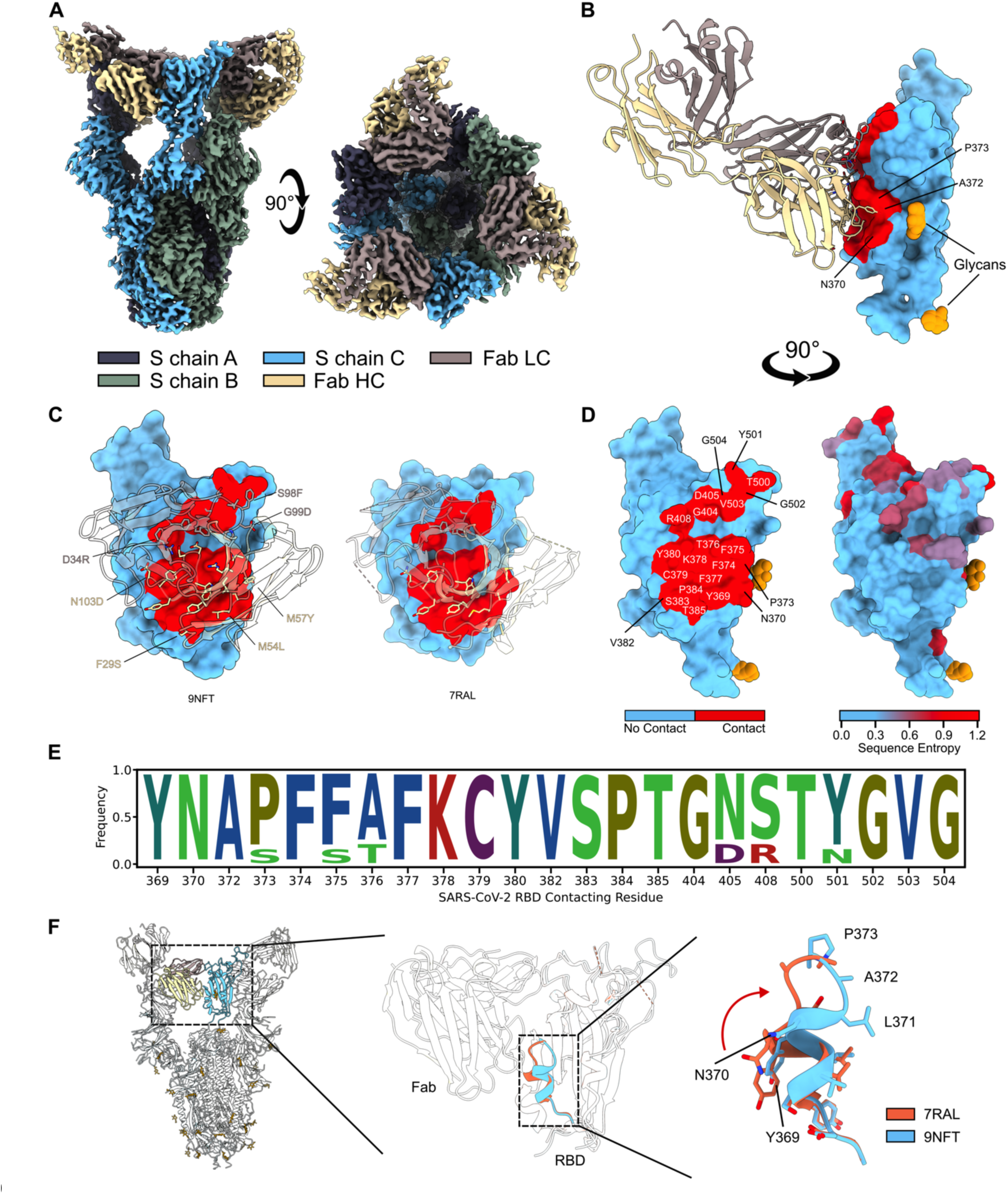
PRO-37587 binds to a conserved cryptic site on the SARS-CoV-2 RBD and can accommodate mutations prevalent in SARS-CoV-2 Omicron lineages. (A) Cryo-EM density map of PRO-37587 Fab bound to SARS-CoV-2 Omicron BA.1 spike trimer. Spike chains (S chain A, B or C) are colored either gray, cyan, or green. PRO-37587 Fab heavy (HC) and light (LC) chains are colored brown and beige, respectively. (B) Molecular model of PRO-37587 Fab bound to a single SARS-CoV-2 Omicron BA.1 RBD. PRO-37587 Fab backbone is depicted as cartoon with colors as shown in (A). SARS-CoV-2 RBD is depicted with a surface representation, with contacts colored red, non-contacts colored cyan, and glycans colored orange. Contacting residues on the RBD are highlighted. (C) Comparison of contacting residues for PRO-37587 Fab (PDB: 9NFT, left) and reference molecule S2X259 Fab (PDB: 7RAL, right). Backbone is depicted as a transparent cartoon and contacts on the Fabs are shown as sticks. Mutational differences between PRO-37587 and S2X259 Fabs that make contacts at the interface are highlighted. SARS-CoV-2 RBD is colored as in (B). (D) SARS-CoV-2 Omicron BA.1 RBD contacts (left) contrasted against sequence conservation (right). Contacts are highlighted with text and are colored as in (B). Sequence conservation is measured as the positional sequence entropy (in bits) of 18 different SARS-CoV-2 variants, including Omicron lineages (SARS-CoV-2 WIV04, D614G, Delta, Omicron BA.1, BA.2, BA.4, BA.5, JN.1, XBB.1.1, LB.1, XBB.1.5, EG.5.1, BQ.1.1, KP.2, XBB.1.16.1, BA.2.86, KP.3 and KP.3.1.1). A higher entropy (red) indicates a larger variation of mutations at that position, and a lower entropy (cyan) indicates less variation. (E) Sequence population of SARS-CoV-2 RBD contacting residues, as measured across the variants used in (D). (F) From left to right, model of SARS-CoV-2 Omicron BA.1 spike with PRO-37587 Fabs bound to all three RBDs, a close up of PRO-37587 Fab bound to SARS-CoV-2 Omicron BA.1 RBD with an overlay of SARS-CoV-2 pre-Omicron RBD (PDB: 7RAL), and a close-up of the altered helix conformation observed in our structure and previously reported. Molecules are colored as in (B), with SARS-CoV-2 pre-Omicron RBD reference colored orange.

A clear structural rationalization as to why PRO-37587 confers broad neutralization and the reference molecule S2X259 does not is difficult because: 1) PRO-37587 and S2X259 Fabs bind in a highly similar manner, and 2) the contacting epitope of PRO-37587 is largely conserved between SARS-CoV-2 variants (**Figures 4B-E**). Of the mutational differences between PRO-37587 and S2X259, only one mutation contacts a non-conserved amino acid in the RBD: the methionine-to-tyrosine substitution at position 57 (M57Y). This mutation introduces a tyrosine that forms a strong π-stacking interaction with proline 373 (P373), an Omicron lineage mutation in the RBD (**Figure 4C**).^34^ It is therefore tempting to speculate that this contact may be key to how broad-spectrum neutralization is achieved. However, the significance of this interaction is not immediately clear since the serine-to-proline mutation at position 373 (S373P) of the RBD is not identified as being an escape mutation to S2X259.^28^

Looking for escape mutations proximal to P373, we identified the serine-to-leucine mutation at position 371 (S371L), a prominent Omicron lineage escape mutation to class 4 anti-RBD mAbs including S2X259.^28,35,36^ Residue position 371 is located toward the end of an α-helix (α2), approximately spanning residues 366-371. Although P373 is a contacting residue, the sidechain at position 371 does not make prominent contacts to either S2X259 or PRO-37587 as seen in a previously reported model (PDB: 7RAL) or our model (PDB: 9NFT). The lack of contacts suggest that the mechanism of neutralization is allosteric in nature.

As ours is the only molecular model of a SARS-CoV-2 Omicron BA.1 spike with the α2 helix seen at high resolution, we could probe the conformational changes observed between SARS-CoV-2 variants to assess if changes impact neutralization. Our structure revealed a significant conformational change at the epitope between Wuhan and Omicron lineages. Specifically, the α2 helix is translocated, with the largest change centered near the RBD escape mutation S371L (**Figure 4F**). While S371L does not directly participate in the epitope/paratope interface, we posited that this mutation alters the conformation of the epitope at this region to confer neutralization resistance. Furthermore, we found that PRO-37587 Fab light chain mutations, methionine-to-leucine at position 55 (M55L) and M57Y, can accommodate the altered contacts to RBD positions 369 and 373, respectively, offering insight into the importance of M57Y in conferring the ability to broadly neutralize. Supporting this structural hypothesis, we observed that reversing mutation M57Y back to the reference mAb amino acid (PRO-37587 Y57M) attenuates broad-spectrum neutralization. However, reversal of serine-to-phenylalanine at position 98 (S98F) and to a lesser extent glycine-to-aspartic acid at position 99 (G99D) and asparagine-to-aspartic acid at position 103 (N103D) also negatively impacted the neutralization profile (**Figure S5A**). In addition, adding M57Y, S98F or M54L to the reference mAb S2X259 without additional mutations is not sufficient for achieving breadth of neutralization (**Figure S5B**). Overall, these data suggest nuanced synergistic effects between mutations.

Summarizing our structural findings, our high-resolution structure of PRO-37587 Fab bound to a SARS-CoV-2 Omicron BA.1 spike protein revealed a conformational change on the RBD near a well-known viral escape mutation. PRO-37587 can accommodate this conformational change, where S2X259 cannot. Mutational data, namely reversion of key antibody binding residues, support this hypothesis, but also point to more complicated effects such as allostery or cooperativity. This highlights a strength of our ML-based optimization approach: while the prior model is structurally driven by nature, experimental data collected in optimization rounds direct exploration towards functionally advantageous regions of the sequence landscape. This allowed us to harness complex binding effects, irrespective of whether the underlying mechanism is easily describable in structural terms.

### Combination of GB-0669 and PRO-37587 shows enhanced SARS-CoV-2 neutralization and resistance to escape *in vitro*

Anti-viral combination therapies offer the advantage of improving neutralization potency and/or increasing barrier to resistance compared to the single agents.^37–41^ GB-0669 and PRO-37587 target two separate binding sites, namely S2 stem helix and RBD class 4 region, which makes them amenable to combination. In addition, GB-0669 and PRO-37587 have two different mechanisms of action (MOAs), respectively inhibition of viral fusion and ACE2 binding, and evidence suggests that targeting distinct spike regions and MOAs may enhance SARS-CoV-2 neutralization.^42–46^ We hypothesized that combination of GB-0669 and PRO-37587 (GB-0669 + PRO-37587) would improve SARS-CoV-2 neutralization potency compared to GB-0669 or PRO-37587 single agents. Experiments with pseudoviruses representative of SARS-CoV-2 D614G, Omicron XBB.1.16.1, EG.5.1 and BA.2.86 confirmed enhanced neutralization potency of the GB-0669 + PRO-37587 combination as demonstrated by the lower EC50 and 90% maximal effective concentration (EC90) values compared to the single agents (**Figures 5A and 5B**). To further substantiate these results, we evaluated the GB-0669 + PRO-37587 combination in live virus neutralization experiments with SARS-CoV-2 Eng20 (ancestral strain), Delta, Omicron BQ.1.1 and XBB.1.1. The clinical-stage anti-RBD mAb bebtelovimab was used as experimental control. As expected, bebtelovimab potently neutralized SARS-CoV-2 Eng20 and Delta but failed to neutralize SARS-CoV-2 BQ.1.1 and XBB.1.1.^47^ Confirming the results of pseudovirus neutralization experiments, the GB-0669 + PRO-37587 combination showed increased neutralization potencies (i.e., lower EC50 and EC90) compared to the single agents (**Figures 5C and 5D**).

**Figure 5.**
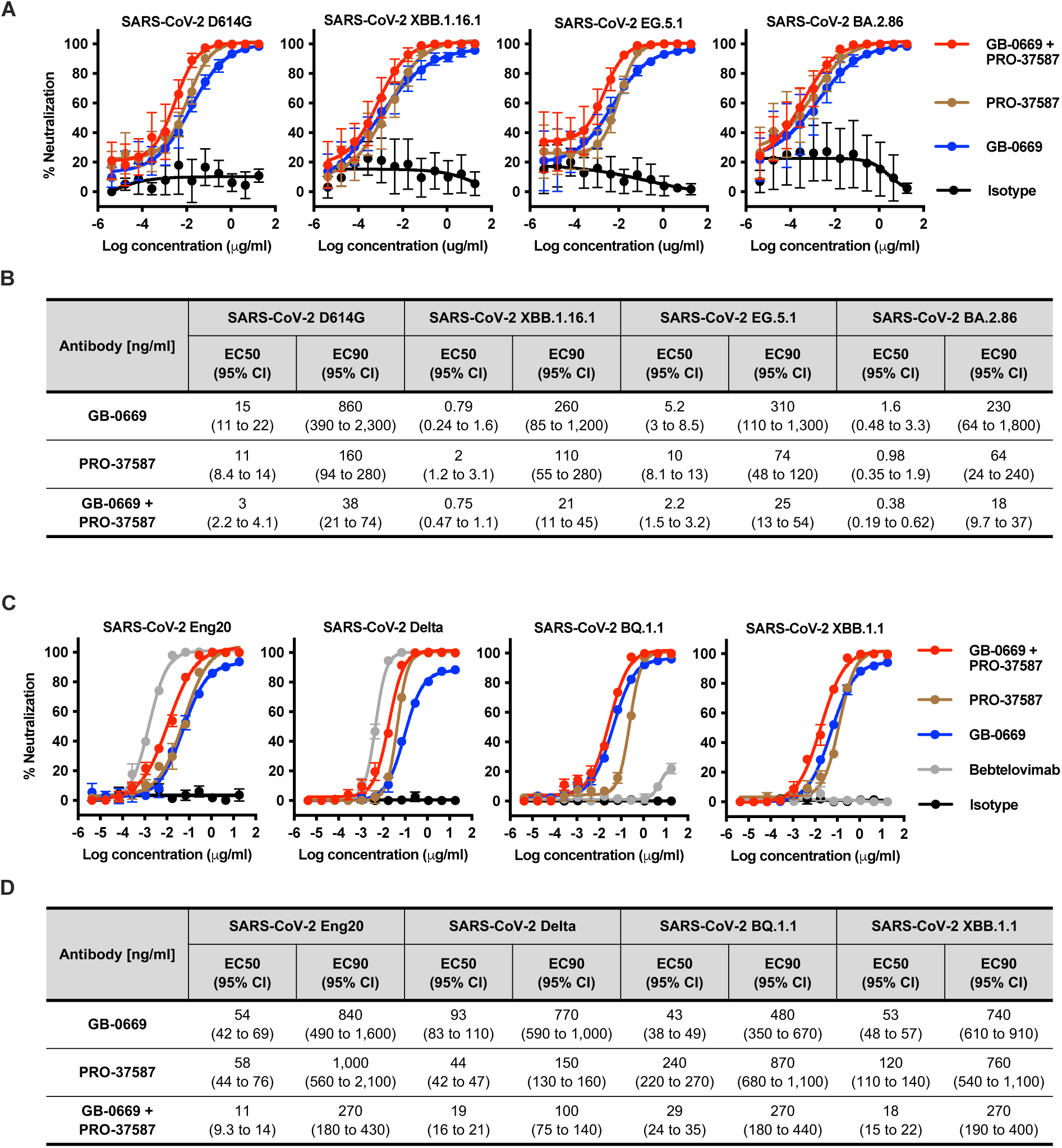
Combination of GB-0669 and PRO-37587 shows improved neutralization of SARS-CoV-2 pseudoviruses and live viruses. (A, B) Neutralization profiles of GB-0669 and PRO-37587 as single agents or in combination (GB-0669 + PRO-37587), and isotype control mAb against pseudoviruses representative of SARS-CoV-2 D614G (ancestral strain), Omicron XBB.1.16.1, EG.5.1 and BA.2.86 variants. (A) Results are reported as percentage (%) neutralization and shown as mean ± SD (representative of four independent experiments, four technical replicates each). (B) EC50 and EC90 values (with 95% confidence intervals in parentheses) relative to the curves shown in (A) are reported in the table. (C, D) Neutralization profiles of GB-0669 and PRO-37587 as single agents and combination (GB-0669 + PRO-37587), bebtelovimab and isotype control mAb against SARS-CoV-2 Eng20 (ancestral strain), Delta, Omicron BQ.1.1 and XBB.1.1 live viruses. (C) Results are reported as percentage (%) neutralization and shown as mean ± SD (representative of one experiment, three technical replicates). (D) EC50 and EC90 values (with 95% confidence intervals in parentheses) relative to the curves shown in (C) are reported in the table. GB-0669, PRO-37587 and bebtelovimab tested as single agents were combined with isotype control mAb at the same concentrations to control for overall mass of GB-0669 and PRO-37587 tested in combination. The x-axis indicates the concentration of each individual antibody in the combinations.

Next, we assessed whether the GB-0669 + PRO-37587 combination posed a higher barrier to resistance using an *in vitro* SARS-CoV-2 Omicron BA.1 live virus escape experiment. The virus was passaged in the presence of GB-0669 + PRO-37587 or the single agents, and the passage numbers (indicated as P*x*, with *x* being the passage number) where cytopathic effect was observed at an antibody concentration of 1-10 µg/ml or > 10 µg/ml were considered as partial escape or complete escape, respectively. GB-0669 was readily escaped by the virus, with partial escape observed on P2 and complete escape on P3. PRO-37587 showed partial escape on P6 and complete escape only on P10. Notably, the combination of GB-0669 and PRO-37587 did not show complete or even partial escape through any of the 10 passages. Sequencing analysis was performed on harvested viruses to characterize escape mutations within the spike protein by mapping the reads to the SARS-CoV-2 Omicron BA.1 input reference genome. Escape from GB-0669 at P3 was associated with histidine-to-tyrosine mutation at position 1156 (H1156Y, all coordinates relative to SARS-CoV-2 Omicron BA.1 spike), which falls within the GB-0669 epitope in the S2 stem helix. Partial escape from PRO-37587 was associated with the aspartic acid-to-glycine mutation at position 402 (D402G) in RBD already detected at P5, but it took a second mutation in RBD (proline-to-serine at position 381 [P381S]) emerging on P10 to lead to complete escape. Finally, only one mutation in the GB-0669 epitope (phenylalanine-to-leucine at position 1153 [F1153L]) was identified on P10 for the GB-669 + PRO-37587 combination (**Figure 6A**). To confirm the impact of such mutations on GB-0669 and PRO-37587 neutralization, we used resistance passaged viruses harvested after emergence of resistance for GB-0669 and PRO-37587 as single agents (respectively at P3 [GB-0669 P3 virus] and P10 [PRO-37587 P10 virus]) or at P10 for the GB-0669 + PRO-37587 combination (GB-0669 + PRO-37587 P10 virus). As controls, we used viruses passaged with no antibodies and harvested at P3 (untreated P3 virus) and P10 (untreated P10 virus). GB-0669 P3 virus was not neutralized by GB-0669 alone but could still be neutralized by PRO-37587 alone and the GB-0669 + PRO-37587 combination. PRO-37587 P10 virus was neutralized by GB-0669 and the GB-0669 + PRO-37587 combination while PRO-37587 showed a marked increase in EC50. Finally, GB-0669 + PRO-37587 P10 virus escaped GB-0669 alone but was neutralized by PRO-37587 alone and the combination, even though the latter showed increased EC50 compared to the respective control virus which again may be explained by loss of GB-0669 contribution to the combination (**Figure 6B**). The escape mutations described above emerged in the context of *in vitro* selective pressure. To assess their relevance within the context of evolutionary pressure exerted by population-level immunity, which can determine the likelihood of emergence of SARS-CoV-2 variants with such mutations, we evaluated the relative frequencies of PRO-37587-associated mutations (X381S and X402G alone and in combination, where X represents any possible amino acid at the indicated position) and GB-0669-associated mutations (X1153L and X1156Y) among publicly available SARS-CoV-2 genomes retrieved from the GISAID database in two time periods: from March 2023 to March 2024, and from January 2020 to March 2024. Interestingly, all mutations were highly infrequent, and the two PRO-37587-associated mutations have never been detected in combination (**Figure 6C**). To further explore the impact of PRO-37587-associated mutations, we performed an *in vitro* viral growth experiment with the PRO-37587 P10 virus and the respective control (untreated P10 virus). Interestingly, PRO-37587 P10 virus showed less efficient viral growth compared to the control virus, suggesting a potentially negative impact of the concomitant presence of the two PRO-37587-associated mutations (**Figure 6D**).

**Figure 6.**
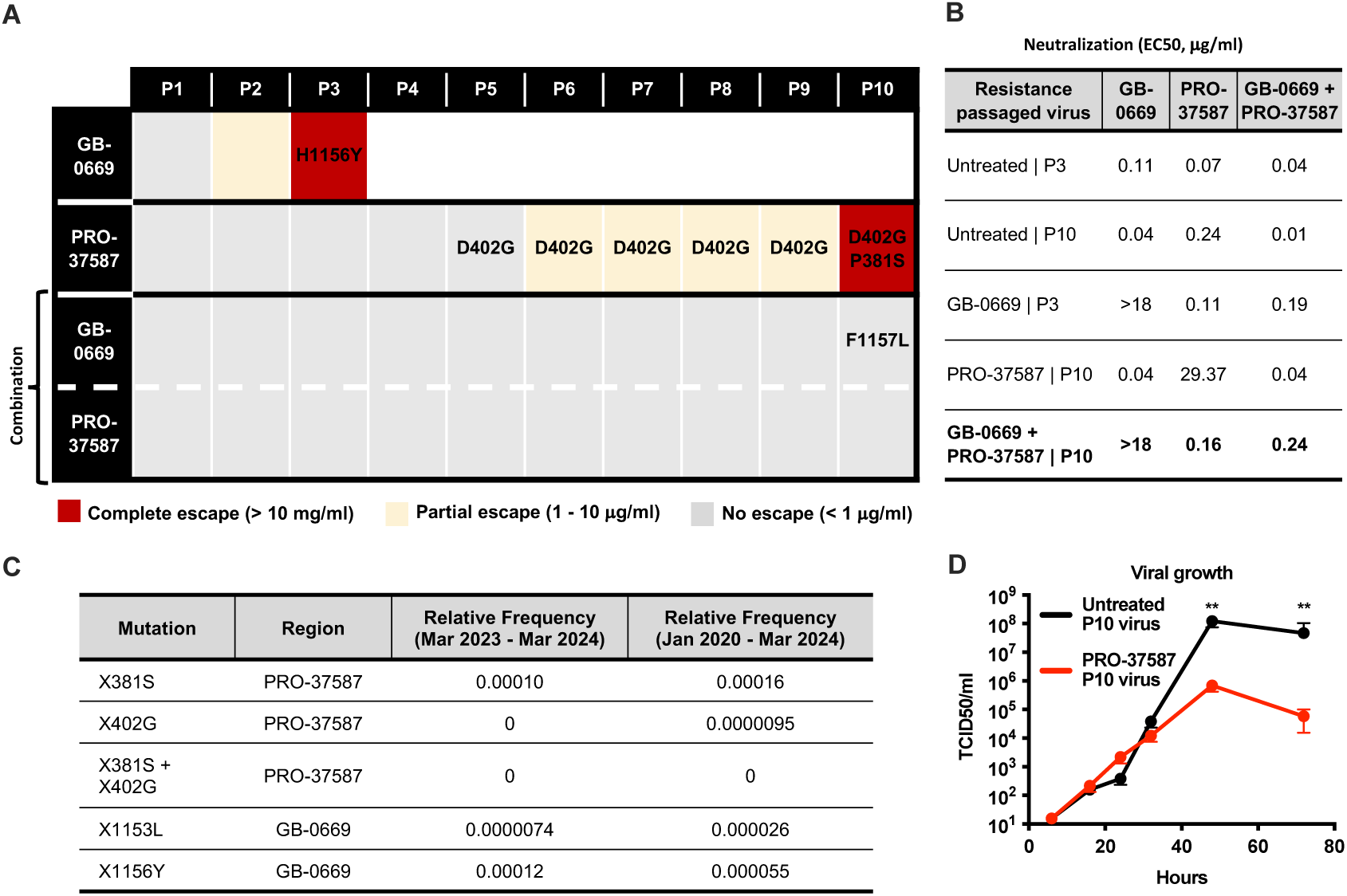
Combination of GB-0669 and PRO-37587 increases barrier of resistance to SARS-CoV-2 Omicron BA.1 live virus escape. GB-0669 and PRO-37587 were tested *in vitro* as single agents and combination with SARS-CoV-2 Omicron BA.1 live virus for 10 passages. (A) Table indicating passage number (reported as P*x*, with *x* being the passage number) and associated mutations that resulted in the virus partially or completely escaping neutralizing activity of the mAbs. Mutation coordinates are relative to the SARS-CoV-2 Omicron BA.1 spike sequence. Complete and partial escape were determined by cytopathic effect (CPE) at each passage. CPE observed at an antibody concentration of 10 µg/ml or above was considered complete escape (indicated with red), while CPE observed at antibody concentrations between 1-10 µg/ml was considered partial escape (indicated as yellow). CPE observed at antibody concentrations < 1 µg/ml, or no CPE, was considered no escape (indicated as gray). (B) Table reporting EC50 (µg/ml) values of GB-0669 and PRO-37587 as single agent and in combination (GB-0669 + PRO-37587) against resistance passaged viruses harvested after emergence of resistance for GB-0669 and PRO-37587 as single agents (respectively at P3 [GB-0669 P3 virus] and P10 [PRO-37587 P10 virus]) or at P10 for the GB-0669 + PRO-37587 combination (GB-0669 + PRO-37587 P10 virus). Viruses passaged with no antibodies and harvested at P3 (untreated P3 virus) and P10 (untreated P10 virus) were used as controls. (C) Table reporting relative frequencies of mutations identified in the escape experiment outlined in (A) among publicly available SARS-CoV-2 genomes representative of the indicated time periods (March [Mar] 2023 - Mar 2024, or January [Jan] 2020 - Mar 2024). Mutation coordinates are referenced relative to the SARS-CoV-2 Omicron BA.1 spike protein. (D) Replication kinetics (TCID50/ml) of the PRO-37587 P10 virus and the respective control (untreated P10 virus). Results are shown as mean ± SD (representative of one experiment, two technical replicates). ** indicates p ≤ 0.01 when comparing PRO-37587 P10 virus and untreated P10 virus across the same timepoints.

Altogether, our results demonstrate that the GB-0669 + PRO-37587 combination potently neutralizes SARS-CoV-2 variants and increases barrier to resistance *in vitro*. In addition, mutations associated with GB-0669 or PRO-37587 escape are highly infrequent among circulating variants likely due to limited population-level immune pressure and/or negative impact on viral growth and fitness.

### Combination of GB-0669 and PRO-37587 protects from *in vivo* SARS-CoV-2 challenge

Considering the promising *in vitro* anti-viral profile of the GB-0669 + PRO-37587 combination, we next tested its activity in a hamster model of SARS-CoV-2 Omicron BA.2 challenge.^48^ One day before SARS-CoV-2 Omicron BA.2 infection, male hamsters were treated with different dose levels of the GB-0669 + PRO-37587 combination as well as GB-0669 and PRO-37587 as single agents at top dose (12 mg/kg). The clinical-stage SARS-CoV-2 mAbs sotrovimab and bebtelovimab were used at top dose as positive controls, and the clinical-stage RSV mAb palivizumab was used as isotype control. Lungs were collected four days post-infection to quantify viral load (TCID50), genomic and subgenomic RNA (the latter being representative of replicating virus). The group treated with GB-0669 + PRO-37587 combination at top dose showed significant reduction in viral load, comparable to the bebtelovimab group and greater than the GB-0669 or PRO-37587 groups (**Figure 7A**). Similar results were obtained when evaluating genomic and subgenomic RNA (**Figures 7B and 7C**).

**Figure 7.**
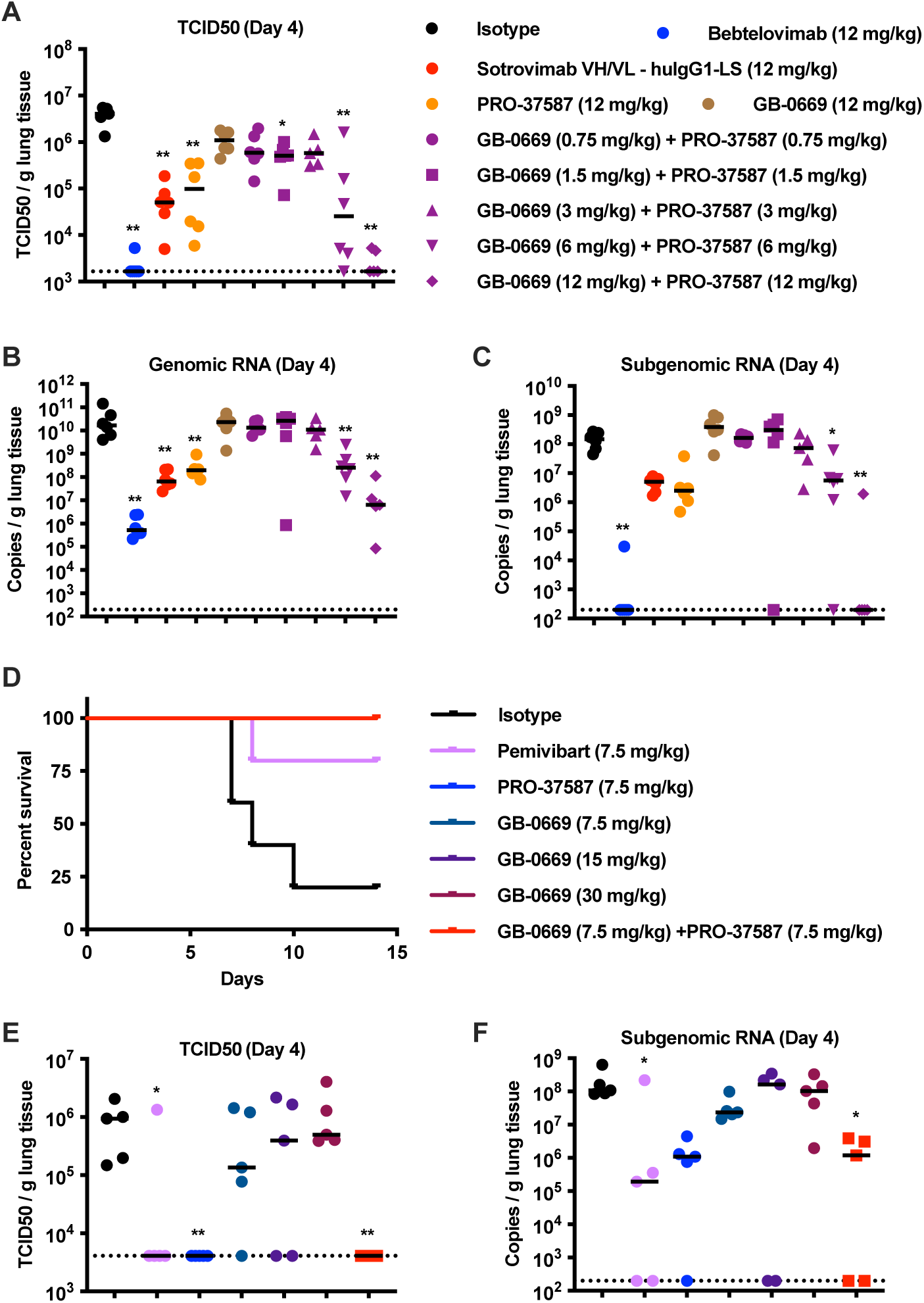
Combination of GB-0669 and PRO-37587 protects from *in vivo* SARS-CoV-2 challenge. (A-C) Golden Syrian hamsters (6 per group) were prophylactically treated with isotype control (palivizumab [anti-RSV F mAb] variable regions expressed as human IgG1 with LS mutation), bebtelovimab, sotrovimab VH/VL - huIgG1-LS (sotrovimab variable regions expressed as human IgG1 with LS mutation), PRO-37587 and GB-0669 as single agents and in combination, at the indicated doses one day prior to challenge with SARS-CoV-2 Omicron BA.2 (2.8x10^3^ TCID50/dose). Four days post-challenge the hamsters were euthanized, lungs were harvested, and the following parameters measured: (A) TCID50 per gram (g) of lung tissue (dotted line indicates lower limit of detection, 1,656 TCID50/g). (B, C) Copies of genomic RNA (B) and subgenomic RNA (C) per g of lung tissue (dotted line indicates lower limit of detection, 200 copies/g). (D-F) hACE2 AC70 mice (10 per group) were prophylactically treated with isotype control (palivizumab [anti-RSV F mAb] variable regions expressed as human IgG1 with LS mutation), pemivibart, PRO-37587 and GB-0669 as single agents and in combination, at the indicated doses one day prior to challenge with SARS-CoV-2 Omicron XBB.1.5. (D) Five mice per group were allowed to progress to study end date and percent survival for each group is reported up to day 14 post-challenge. (E, F) Five mice per group were euthanized on day 4 days post-challenge and lungs were harvested to quantify TCID50 and subgenomic RNA. (E) TCID50 per gram (g) of lung tissue (dotted line indicates lower limit of detection, 4,098 TCID50/g). (F) Copies of subgenomic RNA per g of lung tissue (dotted line indicates lower limit of detection, 200 copies/g). Results in A-C, E and F are reported as individual data points, with lines representing median values. * and ** respectively indicate p ≤ 0.05 and p ≤ 0.01 when comparing experimental groups to isotype group.

We also evaluated the protective efficacy of GB-0669 and PRO-37857 in a lethal human ACE2 (hACE2) AC70 mouse challenge model using SARS-CoV-2 Omicron XBB.1.5 virus. The hACE2 AC70 mouse, which expresses human ACE2, shows greater susceptibility to SARS-CoV-2 infection and is an effective model of COVID-19 disease.^49^ Similar to the hamster study, female hACE2 AC70 mice were treated one day before viral challenge with various dose levels of the GB-0669 + PRO-37857 combination, as well as GB-0669 and PRO-37857 as single agents, up to a top dose of 30 mg/kg. The clinical-stage SARS-CoV-2 mAb pemivibart was used as a positive control,^10^ and the clinical-stage RSV mAb palivizumab was used as an isotype control. Post viral challenge, animals were euthanized if they reached a weight loss of 20% compared to the pre-challenge body weight, or if observed to be moribund. In the isotype control group, all animals except one (20% survival) were euthanized due to significant weight loss observed between days 5 and 10 post-challenge (**Figure 7D**). In contrast, 80% survival was observed in animals treated with pemivibart, and groups treated with GB-0669, PRO-37857, or their combination exhibited 100% survival. Lung tissues were collected on day four post viral challenge to quantify TCID50 and subgenomic RNA levels (**Figures 7E and 7F**). Treatment with PRO-37857 alone, and to a slightly greater extent in combination with GB-0669, significantly reduced lung TCID50 and subgenomic RNA levels.

Overall, these results confirm that increased *in vitro* neutralization potency of the GB-0669 + PRO-37587 combination translates into greater *in vivo* protection from SARS-CoV-2 infection compared to GB-0699 and PRO-37587 as single agents.

## DISCUSSION

The ongoing SARS-CoV-2 pandemic still poses a threat to human health, especially for vulnerable populations in which vaccines have shown limited immunogenicity. Currently, only one mAb (pemivibart) is authorized by the FDA for emergency use in COVID-19 pre-exposure prophylaxis, while another candidate (SA55) is in development in China^50–53^ and many other mAbs have lost emergency use authorization due to their inability to neutralize emerging SARS-CoV-2 variants.^10^ AZD3152 (sipavibart, a half-life extended mAb derived from Omi-42)^54^ has recently demonstrated clinical efficacy in a Phase I/III study, however it is escaped by the mutation F456L that is present in currently circulating SARS-CoV-2 variants (e.g., KP.3.1.1, XEC, LP.8.1).^55^ Of note, recent reports show that both pemivibart and sipavibart are escaped by SARS-CoV-2 JN.1 sublineage variants.^56,57^ More generally, sarbecoviruses are at risk of zoonotic spillover which may lead to future epidemics and pandemics. Therefore, mAbs with broad neutralizing activity against sarbecoviruses could prove highly valuable in the context of the SARS-CoV-2 pandemic and future sarbecovirus zoonosis. Recently, the pan-sarbecovirus mAb VIR-7229 was identified by optimizing S2V29, a SARS-CoV-1 and SARS-CoV-2 cross-reactive mAb isolated from a convalescent donor. VIR-7229 demonstrates a high barrier to resistance *in vitro*, despite targeting one of the most variable regions of the spike protein (the receptor-binding motif) and being susceptible to escape mutations at the highly variable site 456.^58^ In our study, we intentionally chose orthogonal conserved regions on the spike protein, and used structure-conditioned ML-guided protein design with integrated wet- and dry-lab workflows to optimize neutralization activities of anti-S2 stem helix (CV3-25) and class 4 anti-RBD (S2X259) reference mAbs. The resulting mAbs, GB-0669 and PRO-37587, show improved neutralization of SARS-CoV-2 variants and non-SARS-CoV-2 sarbecoviruses as single agents over the respective reference mAbs, and when tested in combination they show enhanced neutralization potencies, higher barrier to *in vitro* resistance and increased *in vivo* efficacy compared to the single agents.

The broadly neutralizing activities of GB-0669 and PRO-37587, compared to their respective reference mAbs, may be explained by their binding profiles. Specifically, GB-0669 binds with higher affinity to the S2 stem helix compared to CV3-25, which our data suggest is accomplished through changes to its conformational ensemble. Additionally, PRO-37587 can accommodate binding to mutations within the RBD that are highly frequent in SARS-CoV-2 Omicron variants. Analysis of our SARS-CoV-2 Omicron BA.1 spike structure reveals a conformational change on the binding region that provides a structural mechanism to the loss of activity for other class 4 anti-RBD mAbs.^28,35,36^ Furthermore, enhancing antibody CDR preorganization and targeting a cryptic site showcases that our design strategy can optimize complex properties of binding that go beyond interfacial energetics, such as dynamic allostery and alternative conformational states.

The mechanism of enhanced neutralization by the GB-0669 + PRO-37587 combination is less clear. The anti-S2 mAb 76E1, which acts through binding of the fusion peptide and inhibition of S2’ cleavage, shows synergistic activity in neutralizing SARS-CoV-2 when combined with recombinant hACE2 or the anti-RBD mAb CB6.^43^ The proposed mechanism is that hACE2- or CB6-induced S1 shedding exposes the fusion peptide, therefore facilitating 76E1 binding. The GB-0669 reference mAb CV3-25 has been reported to increase the frequencies of RBDs in the open conformation upon binding to the S2 stem helix.^26^ Since PRO-37587 targets the inner surface of RBD, which is only exposed in the open conformation, it is tempting to speculate that the GB-0669 binding to the S2 stem helix favors the RBD open conformation and facilitates PRO-37587 binding, resulting in enhanced neutralization. Additional experiments will be required to test these hypotheses and define the molecular basis of enhanced neutralization by the GB-0669 + PRO-37587 combination.

The GB-0669 + PRO-37587 combination potently neutralizes SARS-CoV-2 Omicron variants with distinct mutational profiles, such as EG.5.1 and BA.2.86.^59,60^ Currently, circulating variants are mainly derived from BA.2.86 and have accrued additional mutations to increase escape from antibody-mediated immunity and/or infectivity. For example, mutations L455S, F456L, and Q493E identified in KP.3, but not in BA.2.86, may contribute to the immune-evasive profile of KP.3 and its dominance among other SARS-CoV-2 variants.^61^ Of note, such mutations are not part of the PRO-37587 binding region and, therefore, are not expected to significantly impact its neutralization potency. Additionally, F456L is present in EG.5.1, which is neutralized by PRO-37587. Our *in vitro* escape experiment revealed that mutations leading to full PRO-37587 escape are not represented in currently circulating strains, possibly because such mutations impact viral growth. Taken together, these results suggest a low likelihood that current or future SARS-CoV-2 variants will fully escape PRO-37587. The same considerations apply to GB-0669, especially due to the limited evolutionary pressure exerted on the GB-0669 epitope by population-level immunity, hence a lower likelihood of accruing escape mutations.

In conclusion, we have demonstrated the successful application of structure-conditioned ML-guided protein design and integrated wet- and dry-lab workflows for the optimization of a combination of broadly neutralizing mAbs against sarbecoviruses, with potential implications for pandemic response and preparedness. This approach may be generalizable to the optimization of other anti-viral mAbs with either limited neutralization breadth or potency, thus unlocking opportunities for the development of broadly neutralizing antibodies for the treatment and prophylaxis of viral diseases.

## MATERIALS AND METHODS

### Cell lines

For pseudovirus neutralization, Vero E6 cells (ATCC, CRL-1586) were cultured in MEM/EBSS (Cytiva, SH30024.02), 10% fetal bovine serum (FBS) (Gibco, A38403-01), sodium pyruvate (Gibco, 11360-070), 1X MEM NEAA (Gibco, 11140-050), 1X penicillin/streptomycin (Gibco, 15140-122). TMPRSS2-Vero E6 cells (BPS Bioscience, Inc., 60690-3) were cultured in Vero E6 media with the addition of 3 µg/ml puromycin (InvivoGen, ant-pr). Vero E6 and TMPRSS2-Vero E6 infection media consisted of MEM/EBSS (Cytiva, SH30024.02), 2.5% FBS (Gibco, A38403-01), sodium pyruvate (Gibco, 11360-070), 1X MEM NEAA (Gibco, 11140-050), 1X penicillin/streptomycin (Gibco, 15140-122). For live virus neutralization, Vero E6 cells were cultured in M199 medium GlutaMax (Gibco, 41150087), supplemented with 5% FBS (Gibco, 10500064) and 1X penicillin/streptomycin (Gibco, 15070063). Vero E6 cell infection media consisted of M199 medium GlutaMax, supplemented with 0.4% BSA (Gibco, 15260037) and 1X penicillin/streptomycin. Expi293™ cells (ThermoFisher Scientific, A14527) used for the expression of SARS-CoV-2 spike Fab and BA.1 trimer and were cultured in Gibco Expi293™ Expression Medium (ThermoFisher Scientific, A14351-01).

### Viruses

The following pseudotyped Vesicular Stomatitis Viruses (VSV) and lentiviruses expressing a luciferase reporter that were used for neutralization assays were acquired from BPS Bioscience Inc: spike (WIV1) pseudotyped deltaG-VSV (custom production), spike (SARS-CoV-1) pseudotyped deltaG-VSV (custom production), spike (SARS-CoV-2 D614G) pseudotyped deltaG-VSV (78642), spike (SARS-CoV-2 Delta [B.1.617.2]) pseudotyped deltaG-VSV (78640), spike (SARS-CoV-2 Omicron BA.2) pseudotyped deltaG VSV (78635), spike (SARS-CoV-2 Omicron BA.2.12.1) pseudotyped deltaG VSV (78643), spike (SARS-CoV-2 Omicron BA.4/5) pseudotyped deltaG-VSV (78644), spike (SARS-CoV-2 Omicron BQ.1.1) pseudotyped deltaG-VSV (custom production), spike (SARS-CoV-2 Omicron XBB.1.5) pseudotyped deltaG-VSV (custom production), spike (SARS-CoV-2 Omicron XBB.1.5.10) pseudotyped deltaG-VSV (custom production), spike (WIV1) pseudotyped lentivirus (custom production), spike (SARS-CoV-1) pseudotyped lentivirus (78614), spike (SARS-CoV-2 Delta [B.1.617.2]) pseudotyped lentivirus (78215-2), spike (SARS-CoV-2 Omicron BA.1) pseudotyped lentivirus (78348), spike (SARS-CoV-2 Omicron BA.2) pseudotyped lentivirus (78625). SARS-CoV-2 isolate England/02/2020 (Eng20, ancestral strain) VRS stock P1 (BEI Resources NR-52359), SARS-CoV-2 Delta (B.1.617.2) VOC21APR-02 VRS stock P1 (NIBSC 101030), SARS-CoV-2 Omicron XBB.1.1 VRS stock P1 (NIBSC 101086) and SARS-CoV-2 Omicron BQ.1.1 VRS stock P1 (NIBSC 101082) were used for live virus neutralization. SARS-CoV2 Omicron BA.1 hCoV19/USA/MD-HP20874/2021 stock P1 in V/T/A (BEI Resources NR-56461) was used for resistance passaging. SARS-CoV-2 Omicron BA.2 (BEI Resources, NR-56522) and XBB.1.5 (BEI Resources, NR-59105) were used for *in vivo* studies.

### Antibodies

GB-0669 was produced at Lonza Biologics. CV3-25,^25,26^ S2X259,^23^ the isotype control (palivizumab [anti-RSV F mAb] variable regions [https://www.imgt.org/3Dstructure-DB/cgi/details.cgi?pdbcode=7753] expressed as human IgG1 with LS mutation), bebtelovimab (https://www.imgt.org/3Dstructure-DB/cgi/details.cgi?pdbcode=12145), and sotrovimab VH/VL -huIgG1-LS (sotrovimab variable regions [https://www.imgt.org/3Dstructure-DB/cgi/details.cgi?pdbcode=11766] expressed as human IgG1 with LS mutation) were produced at GenScript. PRO-37587 as well as all early screening designs were produced at Generate:Biomedicines. Briefly, full-length human IgG1 molecules were generated by subcloning the variable heavy (VH) chain and variable light (VL) chains (synthesized externally; Integrated DNA Technologies) into mammalian expression vectors containing a CMV promoter sequence, signal peptide, and corresponding constant regions using the Golden Gate cloning method (New England Biolabs, Catalog # E1601L). Purified DNA was transfected into ExpiCHO™ cells following the manufacturer’s recommended methods (Thermo-Fisher Scientific, Catalog # A29133). Expressed IgG molecules were affinity purified by MabSelect Sure Protein A affinity resin (Cytiva, Catalog #17543802) and in some cases, further purified by size exclusion chromatography using Superdex200 resin (Cytiva, Catalog # 17104301) following the manufacturer’s recommended methods.

### Peptides and Recombinant Proteins

Custom biotinylated SARS-CoV-2 spike S2 peptides (1133-1147, 1137-1151, 1141-1155, 1145-1159, 1149-1163, 1153-1167, 1157-1171, 1149-1167, 1143-1162) and an HIV-1 Env peptide with a biotin molecule conjugated to the amino-terminus were synthesized at Biosynth as previously described.^25^ Recombinant His-tagged spike trimers and spike RBDs were purchased from Acro Biosystems: SARS-CoV-1 spike trimer (SPN-S52H6), SARS-CoV-2 D614G spike trimer (SPN-C52H3), SARS-CoV-2 Delta (B.1.617.2) spike trimer (SPN-C52He), SARS-CoV-2 Omicron BA.1 spike trimer (SPN-C52Hz), SARS-CoV-2 Omicron BA.1.1 spike trimer (SPN-C5224), SARS-CoV-2 Omicron BA.2 spike trimer (SPN-C5223), SARS-CoV-2 Omicron BA.2.12.1 spike trimer (SPN-C522d), SARS-CoV-2 Omicron BA.2.3.20 spike trimer (SPN-C522n), SARS-CoV-2 Omicron BA.2.75 spike trimer (SPN-C522f), SARS-CoV-2 Omicron BA.2.75.2 spike trimer (SPN-C522r), SARS-CoV-2 Omicron BA.4 spike trimer (SPN-C5229), SARS-CoV-2 Omicron BA.4.6 spike trimer (SPN-C522m), SARS-CoV-2 Omicron BF.7 spike trimer (SPN-C522q), SARS-CoV-2 Omicron BN.1 spike trimer (SPN-C524b), SARS-CoV-2 Omicron BQ.1 spike trimer (SPN-C524a), SARS-CoV-2 Omicron BQ.1.1 spike trimer (SPN-C522s), SARS-CoV-2 Omicron XBB spike trimer (SPN-C5248), SARS-CoV-2 Omicron XBB.1 spike trimer (SPN-C522t), SARS-CoV-2 Omicron XBB.1.5 spike trimer (SPN-C524i), SARS-CoV-1 RBD protein (SPD-S52H6), SARS-CoV-2 (ancestral strain) RBD protein (SPD-C52H3), SARS-CoV-2 Delta (B.1.617.2) RBD protein (SPD-C52Hh), SARS-CoV-2 Omicron BA.1 RBD protein (SPD-C522e), SARS-CoV-2 Omicron BA.2 RBD protein (SDP-C522g), SARS-CoV-2 Omicron BA.4/5 RBD protein (SPD-C522r), SARS-CoV-2 Omicron BQ.1.1 RBD protein (SPD-C5240), SARS-CoV-2 Omicron XBB.1.5 RBD protein (SPD-C5242). Recombinant WIV1 spike trimer was produced at Generate:Biomedicines using a method similar to that used for the recombinant SARS-CoV-2 Omicron BA.1 spike trimer described below.

### ML-Guided Protein Optimization

To generate antibodies with improved neutralization, we employed a general integrated wet- and dry-lab approach that can be used to optimize effectively arbitrary combinations of measurable protein functions. The main challenge in optimizing proteins is that the combinatorially large sequence space is almost entirely consistent of inactive variants (i.e., those that fail to fold and/or have no measurable levels of function). That means that even starting from known functional sequences, one expects random perturbations to generally be inactive. While predictive sequence-to-function models can guide sampling toward higher success rates, these models are rarely available *a priori*, especially in cases requiring activity against new targets (e.g., neutralization against a novel virus).

To address this limitation and enable efficient optimization that starts with a known variant, we use a general protein structure-conditioned sequence generative model to serve as a strong prior for sequence exploration. Even without prior knowledge of target functions or properties, this model can propose diverse sequences likely to adopt a structure similar to the starting functional protein, thus increasing the probability of observing function. From experimental measurements of these initial variants, we then built supervised sequence-to-function (seq2func) models to further guide optimization based on both structural and functional insights.

As our prior we used an in-house structure-conditioned sequence-generative model similar to the sequence design module within Chroma (see https://github.com/generatebio/chroma).^62^ The first round focused on exploration, using only the structure of the reference molecule to sample an initial batch of novel sequences. Specifically, we enforced diversity and constrained the distance from the reference sequence (between 5 and 25 mutations, in different sub-batches). For the SARS-CoV-2 S2 stem helix campaign, we employed PDB structure 7NAB (CV3-25 Fab bound to S2 stem helix peptide), and for the RBD campaign, PDB structure 7M7W (S2X259 Fab bound to RBD class 4 region).

Subsequent rounds were more exploitative, guided by both the structure-based design model and surrogate seq2func models trained on the experimental data from previous rounds. These models predicted “fitness” based on the input sequence (**Figures S1A and S3A**) and were trained via ridge-regression with cross-validation. Given the scale of characterized sequences (hundreds of sequences per round), we used only linear, position-wise sequence features (i.e., no higher-order dependencies). Where multiple experimental readouts were available (e.g., binding, neutralization across viral strains), these were combined into composite scores to reflect a joint optimization objective (e.g., neutralization against multiple strains). **Figures S1 and S3** outline binding and pseudovirus neutralization measurements for each round, which were used for model training in subsequent rounds. Developability measurements (e.g., self-association, monomericity, polyspecific reactivity) were also collected in most rounds and used to exclude sequences when suboptimal behaviors were observed.

### DELFIA binding analysis

Three hundred and eighty-four well plates were either coated with recombinant spike trimers (5 μg/ml) and incubated overnight at 4°C or coated with neutravidin (2 μg/ml) overnight at room temperature. Neutravidin (Life Technologies, 31000) coated plates were then washed and coated with biotinylated peptides (SARS-CoV-2 spike S2 peptides, HIV-1 Env negative control peptide; 50 nM). All plates were washed, incubated with blocking solution (ThermoFisher, 28360) for 1 hour at room temperature, washed again and incubated with antibodies for 1 hour at room temperature. Antibody binding was assessed using a 12-point titration curve in technical quadruplicate (1:4 serial dilutions prepared in blocking solution and starting at 18 μg/ml). Plates were washed and incubated with Eu-labeled anti-human IgG (PerkinElmer, 1244-330) secondary antibody (0.1 μg/ml) for 30 minutes at room temperature. Finally, plates were washed and incubated with DELFIA Enhancement solution (PerkinElmer, 40010010) for 15 minutes. Time-resolved fluorescence was recorded at 615 nm with the EnVision plate reader (PerkinElmer, serial #411177671).

### SPR binding analysis

All SPR experiments were performed on a Biacore 8K+ (Cytiva, serial #2873569) instrument at 25°C using 1X HBS-EP+, pH 7.6 (Teknova, H8022) as the sample and running buffer. To evaluate the kinetics of the binding interactions described, several capture-based approaches were exploited, including a biotin CAPture kit (Cytiva, 28920233) and different capture antibodies immobilized onto a CM4 sensor chip (Cytiva, 29104989) using standard EDC/NHS chemistry. For experiments involving a capture antibody, a CM4 sensor surface was prepared across flow cells 1 and 2 and all 8 channels in series by activating with a 1:1 (v/v) mixture of 400 mM EDC and 100 mM NHS (Cytiva, BR100633) for 420 seconds, injecting a capture antibody diluted in sodium acetate, pH 4.5 (Cytiva, BR100350) for 420 seconds and deactivating the surface using 1 M ethanolamine, pH 8.5 (Cytiva, BR100633) for 420 seconds.

To assess the kinetics of GB-0669 and CV3-25 Fabs binding to peptides spanning the SARS-CoV-2 spike S2 stem helix region, the biotinylated peptides were captured using the CAP chip and biotin CAPture reagent to achieve a capture level of approximately 10 to 20 response units (RU). GB-0669 and CV3-25 Fabs were sequentially injected over the surface from low to high concentrations (1.2 nM - 300 nM, 3-fold serial dilution) at a flow rate of 30 μl/minute for 180 seconds followed by a dissociation time of 900 seconds. Biotin CAPture reagent injection and surface regeneration conditions were performed according to manufacturer instructions.

To obtain the kinetics of GB-0669 and CV3-25 Fabs binding to a panel of SARS-CoV-2 spike trimers, approximately 250 RU of each trimer was captured on a CM4 sensor chip immobilized with a mouse anti-histidine monoclonal antibody (Genscript, A00186). Each concentration of GB-0669 and CV3-25 Fabs (0.8 nM - 600 nM, 3-fold serial dilution) was injected over the surface in separate cycles at a flow rate of 30 μl/minute for 120 seconds followed by a dissociation time of 600 seconds. The surface was regenerated by injecting 10 mM glycine-HCl, pH 2.1 (Cytiva, 29234601) for 20 seconds at a flow rate of 10 μl/minute in between each cycle.

To determine the kinetics of PRO-37587 and S2X259 mAbs binding to a panel of SARS-CoV-2 RBD proteins, approximately 150 RU of each antibody was captured on a CM4 sensor chip immobilized with a goat anti-human polyclonal antibody (Southern Biotech, 2014-01). Each concentration of SARS-CoV-2 RBD proteins (0.04 nM - 30 nM, 0.08 nM - 60 nM or 1.4 nM - 1000 nM, 3-fold serial dilution) was injected over the surface in separate cycles at a flow rate of 30 μl/minute for 180 seconds followed by a dissociation time of 3600 seconds, 1200 seconds or 600 seconds, depending on the concentration series, respectively. The surface was regenerated by injecting two pulses of 1.5% phosphoric acid (ThermoFisher Scientific, 295700010) for 30 seconds at a flow rate of 30 μl/minute in between each cycle.

Kinetic parameters for each concentration series were obtained by double referencing and globally fitting the data to a 1:1 binding model using the Biacore Insight Evaluation software (Cytiva, version 4.0.8.19879). Results were expressed as average ± standard deviation of one independent experiment with technical replicates when appropriate. It is recommended to attain at least a 5% decrease in the binding response for the highest antigen concentration tested during the dissociation phase to accurately measure slow dissociation rate constants. For binding interactions reaching this recommendation for the 3600 seconds dissociation phase, the kinetic and affinity values were reported; however, for binding interactions that did not satisfy this condition, the dissociation rate constant k_d_ was fixed at ≤1.42 x 10^-5^ seconds^-1^ and affinity values were reported with this consideration.

### Yeast display of RBDs and binding analysis

Plates of clonal yeast strains were prepared with cells in each well expressing different RBDs. These yeast strains were then stained with purified mAbs, and flow cytometry was used to determine the relative binding across the different RBDs. Briefly, competent EBY100 yeast cells (ATCC, MYA4941) were prepared using the Frozen-EZ Yeast Transformation Kit (Zymo Research, T2001). Yeast cells were combined with linearized backbone and insert DNA at a ratio of 5 μl cells/50 ng backbone/50 ng insert DNA per reaction. RBD constructs (**Table S3**) fused to a c-myc epitope tag were synthesized as DNA fragments (Twist Biosciences) with overhangs for cloning into a yeast display vector. DNA inserts and digested vectors were transformed into yeast, and the full plasmid was generated *in vivo* by homologous recombination.^63^ Cultures were induced for RBD expression at an OD600 of 0.5 in selective growth media (SGCAA, 10% SDCAA, 1% Penicillin-Streptomycin) for 22 hours shaking at 20°C. Cells were washed and resuspended in PBS with 0.1% BSA at an OD600 of 3. Induced cells were stained for binding to purified IgG and anti-c-myc (Exalpha, ACMYC) for RBD expression. Cells were incubated overnight with IgG diluted to a concentration of 1 nM with 1:1000 chicken c-myc epitope tag added. Cells were spun down and washed three times in PBS with 0.1% BSA before incubation with R-Phycoerythrin AffiniPure™ F(ab’)₂ Fragment Donkey Anti-Chicken IgY (IgG) (H+L) PE (Jackson, 703-116-155) and AlexaFluor 647 anti-human IgG (Jackson, 109-605-098) at 4°C in the dark for 15 minutes. Cells were spun down and washed three times in PBS with 0.1% BSA. Fluorescence was measured on iQue3 (Sartorius, serial #3609) and analyzed using the FlowJo v10.8.1 software package (FlowJo LLC). Relative binding results are shown as median fluorescence intensity of the mAb signal divided by the median fluorescence intensity of the expression signal for each combination and reported as heatmaps relative to the mAb/RBD combination with the highest overall signal.

### Cryo-EM and X-ray crystallography sample preparation, data collection and data processing

GB-0669 Fab and SARS-CoV-2 spike S2 [aa 1149-1167] peptide were added in a 1:1 molar ratio, incubated on ice for 20 minutes, concentrated and injected onto a 16/900 Superdex 200 pg column (buffer: 25 mM HEPES, pH 7.5, 150 mM NaCl). Peak fractions were analyzed by SDS-PAGE and combined and concentrated to 10 mg/ml. An ARI Crystal Gryphon drop-setting robot used 96-3 Intelliplates to set crystallization drops for the complex at 10 mg/ml at three ratios (1:1, 2:1, and 3:1) of complex to the crystallization condition. The 96-well MCSG-3 commercial screen was set and incubated at 4°C and 20°C. Crystals from the MCSG-2 A2 (1:1 ratio) at 4°C were harvested and cryoprotected in 20% glycerol. Crystals were sent to the NSLS2 synchrotron and X-ray data sets were collected on AMX beamline with an Eiger X 9M detector. The data was processed using DIALS and XDS to 1.93 Å. Once processed, the data set was phased by molecular replacement using Phaser MR with an AlphaFold model of GB-0669 Fab and PDB ID 7RAQ for the peptide. Several rounds of refinement were performed using Phenix and Coot.

### Expression and purification of SARS-CoV-2 Omicron BA.1 spike trimer

SARS-CoV-2 Omicron BA.1 spike trimer was expressed in Expi293™ (ThermoFisher Scientific, catalog # A14527) cells following the Gibco™ Expi293™ Expression System protocol. In brief, three million cells were transfected with approximately 1 mg of plasmid DNA. The cells were incubated at 37°C, with 80% relative humidity and 8% CO_2_ on an orbital shaker at 150 RPM. Four days post-transfection, the cells were harvested and pelleted at 3,900 x g for 30 minutes at 4°C. The supernatant was decanted into 0.22 µm filter units and stored at 4°C until purification. The SARS-CoV-2 BA.1 spike trimer was purified using Nickel Sepharose Excel (Cytiva, catalog # 17371201). The supernatant was incubated with Nickel Sepharose Excel resin overnight at 4°C and purified by gravity flow.

### Expression and purification of Fabs targeting SARS-CoV-2 Omicron BA.1 spike trimer

GB-0669, PRO-27522 (class 4 anti-RBD), and PRO-37587 (class 4 anti-RBD) Fabs were expressed in Expi293™ cells following the same protocol used for SARS-CoV-2 Omicron BA.1 spike trimer. Supernatants containing GB-0669 or class 4 anti-RBD Fabs were incubated with either Capto™ L resin (Cytiva, catalog # 17547802), or LambdaFabSelect resin (Cytiva, catalog # 17548201) overnight at 4°C. The mixtures of supernatant and resin were loaded onto a 10 ml disposable column equilibrated in 1X PBS (pH 7.4). The column was then washed with 10 CV of 1X PBS (pH 7.4). The Fabs were eluted with 50 mM Glycine (pH 2.5). The proteins were immediately neutralized with 1 M Tris-HCl (pH 8.0), buffer exchanged using a PD-10 desalting column (Cytiva, 17085101) and eluted with 1X PBS (pH 7.4). The purified proteins were concentrated and kept at 4°C until complexation with the SARS-CoV-2 Omicron BA.1 spike trimer.

### Complexation of SARS-CoV-2 Omicron BA.1 spike trimer with GB-0669 and class 4 anti-RBD Fabs

Multiplexing of the SARS-CoV-2 Omicron BA.1 spike trimer was achieved by complexing the spike trimer with GB-0669 and PRO-27522 Fabs at a 1:2:2 molar ratio, whereas for complex formation between SARS-CoV-2 Omicron BA.1 spike trimer and PRO-37587 Fab the proteins were mixed at a molar ratio of 1:2. All samples were incubated at 4°C overnight with gentle mixing. The complexes were purified by size exclusion chromatography (SEC) on a Superose 6 Increase 10/300GL column (Cytiva, catalog # 29091596) equilibrated in 50 mM HEPES (pH 8.0), 150 mM NaCl. Peak fractions were analyzed by SDS-PAGE and analytical SEC (aSEC) on an SRT-C SEC-500 (Sepax, PN: 235500-4630) column equilibrated in 50 mM HEPES (pH 8.0), 150 mM NaCl.

### Cryo-EM sample preparation of SARS-CoV-2 Omicron BA.1 spike trimer and PRO-37587 Fab complexes

Four microliters of each spike trimer:Fab complexes were applied to Quantifoil gold grids (R1.2/1.3, [Electron Microscopy Sciences, catalog # 261655]) which were glow-discharged for 30 seconds at a plasma current of 0.15 mA with negative polarity. The grids were vitrified on the Vitrobot Mark IV (Thermo Fisher Scientific).

### Cryo-EM image and data processing of SARS-CoV-2 Omicron BA.1 spike trimer:GB-0669 Fab:PRO-27587 complexes

Cryo-EM images were acquired on a Glacios cryo-TEM using EPU software (v3.2). The Glacios was operated at 200 kV with a Falcon4i direct electron detector and a Selectris energy filter with a zero-loss slit width of 10 eV. 5160 movies were collected at 130,000x magnification at a pixel size of 0.876 Å. The total dose per movie was 51.3 electrons per square angstrom. The targeted defocus range was between 0.5-2.4 μm. All computational steps were performed using the cryoSPARC (v4.1.1) software suite (Structura Biotechnology, v4.1.1) and ChimeraX (UCSF RVBI, v1.5) molecular visualization software.^64,65^ Movies in EER format were imported and fractionated into 40 frames and sampled at the physical pixel size. Beam-induced motion correction, per-frame dose-weighting, and CTF estimation were performed using the patch motion correction and patch CTF jobs in cryoSPARC (v4.1.1). Exposures were selected semi-automatically using the interactive exposure curation tool. After setting stringent cut-offs on CTF fit resolution, defocus, and relative ice thickness, 2,921 images were selected for subsequent processing. Template-based particle picking was performed with projections from a 3D map of the SARS-CoV-2 Omicron BA.1 spike trimer low-passed filtered to 20 Å. 466,074 particles were extracted from 2921 micrographs. 2D classification was performed with 200 classes to identify incorrectly picked or broken particles, retaining 87,564 particles for subsequent processing. *Ab initio* reconstruction with three classes was performed, yielding an initial 3D map with the expected size and shape of the SARS-CoV-2 BA.1 spike trimer from a subset of 51,996 particles. Subsequent non-uniform 3D refinement in cryoSPARC of these particles yielded a ‘consensus map’ at 3.6 Å resolution by gold-standard FSC criteria.

### Focused classification and model fitting of SARS-CoV-2 Omicron BA.1 spike trimer:GB-0669 Fab complex

An atomic model of the SARS-CoV-2 Omicron BA.1 spike trimer without bound Fabs was computationally docked into the consensus map with the ‘fit in map’ tool in ChimeraX (UCSF RVBI, v1.5). The map fits snugly into the density, except for several unaccounted densities at the RBD and S2 stem helix regions of the map. Gaussian low-pass filtering of the map revealed these densities to be dumbbell shaped with a hole in the center, as expected for Fab molecules. The ‘Segment Map’ tool in ChimeraX and cryoSPARC’s volume tools were used to produce a focused mask around the S2 stem helix that enclosed the propeller-shaped putative Fab densities. Focused 3D classification without alignment in cryoSPARC was performed with 10 classes to identify a subset of 9,822 particles with stronger density of the putative S2 stem-helix bound Fabs. This subset was refined without applying symmetry using non-uniform refinement to 5.7 Å resolution by gold-standard FSC criteria. Local resolution estimation in cryoSPARC showed that the S2 helix binding Fabs have lower local resolution than the core of the SARS-CoV-2 Omicron BA.1 spike trimer. The CDR-containing domains of the Fabs were approximately 8 Å local resolution, while the flexibly-associated framework domains of the Fabs were approximately 14-20 Å local resolution. For this reason, a Gaussian low-pass filter in ChimeraX (UCSF RVBI, v1.5) with a width of 3 Å was applied. One Fab is distinctly weaker than the other two. Nevertheless, these densities were sufficient to unambiguously dock three copies of GB-0669 Fab-S2 complex crystal structure (PDB: 9NHP) into each of the three S2 helix Fab densities using ChimeraX’s ‘fit in map’ tool. The structure of the class 4 anti-RBD PRO-27522 Fab was also docked into dumbbell-shaped density at the RBD.

### Cryo-EM data collection and processing of SARS-CoV-2 Omicron BA.1 spike trimer:PRO-37587 Fab complex

Cryo-EM images were acquired on a Krios G4 cryo-TEM using EPU software (v3.2). The Krios was operated at 300 kV with a Falcon4i direct electron detector and a Selectris energy filter with a zero-loss slit width of 10 eV. Two sets of movies were collected at 165,000x magnification at a pixel size of 0.730 Å, one with 3,393 movies and the other with 7272 movies. The total dose per movie was 46.30 electrons per square angstrom. The targeted defocus range was between 0.5-2.4 μm.

Initial computational processing for CTF estimation and beam induced motion correction followed similar steps to that of GB-0669. Two processing streams were run in parallel before being merged for further refinement (**Figures S6 and S7**). After processing, 3,393 and 5,758 images were used with the CryoSPARC blob-picker to extract 143,663 particles and 221,815 particles for the first and second streams, respectively. 2D classification was performed with 200 and 100 classes to identify incorrectly picked or broken particles, retaining 79,652 particles and 128,742 particles for subsequent processing.

For the first stream, *Ab initio* reconstruction with three classes was performed, yielding an initial 3D map with the expected size and shape of the SARS-CoV-2 BA.1 spike trimer from a subset of 33,024 particles. A non-uniform refinement of these particles with C1 symmetry yielded a map of 3.03 Å resolution, however, the overall look of the map was poor. For the second stream, Ab initio reconstruction was performed with four classes, where two of the four classes were selected, yielding 79,171 particles for further processing. The particles from both streams were then merged to yield 112,195 particles, for which subsequent non-uniform 3D refinement in CryoSPARC with C3 symmetry yielded a ‘consensus map’ at 2.97 Å resolution by gold-standard FSC criteria (**Figure S8A**). Particles were then symmetry expanded around the C3 axis to align individual asymmetric subunits, creating 336,585 particles.

A focus mask was generated around the region comprising PRO-37587 bound to the RBD. This mask was generated by docking a postulated model of our designed PRO-37587 bound to the SARS-CoV-2 RBD into one of the asymmetric subunits and using ChimeraX’s volume tools to generate a density map around the structure to a 15 Å resolution. The focus mask was then binarized in CryoSPARC and dilated by 2 pixels with a soft padding width of 20 pixels.

A local refinement was performed using the focus mask. Following this refinement, a 3D classification without alignment was used to generate 10 unique classes. These classes detail the range of motion of the PRO-37587:RBD complex as aligned to S2 of the Spike protein. Interpolations between these classes were used to generate supplementary video S1.

Three of the classes from the 3D classification that were inspected and deemed most similar to one another were combined to form a set of 102,986 particles. A local refinement of these particles was then performed using the focus mask, yielding a resolution of 3.46 Å. We then performed another 3D classification without alignment to generate 3 unique classes. We then selected the highest resolution class, containing 100,545 particles to perform a final local refinement, which yielded a resolution of 3.00 Å by gold-standard FSC criteria (**Figure S8B**). The consensus map was then combined with the final local refinement map and symmetrized around the C3 axis to yield a final composite map of the full spike trimer.

### Molecular model of SARS-CoV-2 Omicron BA.1 spike trimer:PRO-37587 Fab complex

A molecular model was built into our final composite map of PRO-37587 bound to the SARS-CoV-2 Omicron BA.1 spike trimer. PDB 6VXX was used as a starting template, where individual domains of were extracted and docked into a single asymmetric unit of the map as rigid bodies. After docking, the domains were stitched together to form a single chain of the S1/S2 Spike protein. We then utilized this template with our foundation model Chroma, to build a homology model of the BA.1 spike asymmetric unit.

The molecular model of the asymmetric unit was then further refined using density guided MD simulations that leverage the Cryo-EM density map as an additional potential energy.^66^ Simulations were set up by placing the initial structure in a dodecahedron box of explicit TIP3P solvent, with a minimum distance of 1.0 nm from the protein to the nearest edge. Energy minimization proceeded using a steepest descent algorithm until the maximum force fell below 100 kJ/mol/nm, using a step size of 0.01 nm and a distance cutoff of 1.2 nm for the neighbor list, Coulomb interactions, and van der Waals interactions. Following equilibration, production runs comprised four 50 ns simulations using either cross-correlation or relative-entropy as the similarity measure for the MDFF potential. The conformation with the best fit to the experimental density map was then symmetrized around the C3 axis to form a trimer and used for further refinement. A template of PRO-37587 was then docked into the density to generate a full-length initial model.

The full-length model underwent iterations of real-space refinement using PHENIX followed by manual adjustments using COOT (0.9.7 version).

Sequence entropy analysis (**Figures 4D-E**) was performed by first aligning the following SARS-CoV-2 sequences: SARS-CoV-2 WIV04, D614G, Delta, Omicron BA.1, BA.2, BA.4, BA.5, JN.1, XBB.1.1, LB.1, XBB.1.5, EG.5.1, BQ.1.1, KP.2, XBB.1.16.1, BA.2.86, KP.3 and KP.3.1.1. At each position we compute a sequence entropy,

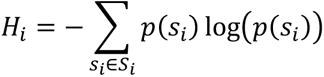

where *p*(*s_i_*) is the probability of observing mutation *s_i_* across all the mutations observed at that position, *S_i_*. Sequence entropies are colored on the surface of the RBD (**Figure 4D**) or shown as a logo-plot for residues contacting PRO-37587 (**Figure 4E**).

### Molecular dynamics simulations of GB-0669 and CV3-25

Simulations were run with Gromacs 2023.4 using the AMBER99sb-ildn forcefield with explicit TIP3P solvent.^67–69^ Initial structures of GB-0669 (PDB: 9NHP) and CV3-25 (PDB: 7NAB) were solvated in a dodecahedron box that extended 1.0 nm beyond the protein in any dimension. These systems were energy minimized using a steepest descent algorithm until the maximum force fell below 100 kJ/mol/nm using a step size of 0.01 nm and a cutoff distance of 1.2 nm for the neighbor list, Coulomb interactions, and van der Waals interactions. For production runs, all bonds were constrained with the LINCS algorithm and virtual sites were used to allow for a 4 fs time step.^70,71^ Cutoffs of 1.1 nm were used for the neighbor list and 0.9 nm for Coulomb interactions and van der Waals interactions. The stochastic velocity rescaling (v-rescale) thermostat^72^ was used to hold the temperature at 300 K and conformations were stored every 20 ps. To enhance sampling for each system, we ran FAST simulations.^73^ Each set of FAST-RMSD simulations consisted of 5 μs of aggregate simulation time: 5 rounds, of 10 simulations per round, with a simulation length of 100 ns. For each system, the directed component to the FAST-ranking was the RMSD of CDRs to the initial starting structure. Markov state models (MSMs) were built for each simulation set using enspara.^74^ The construction of each MSM followed the same basic protocol: 1) cluster conformations into discrete conformational states, 2) count transitions between these states, 3) generate conditional transition probabilities. For clustering, backbone heavy atoms were used with a k-centers clustering algorithm until every cluster center had a radius of less than 1.4 Å. A lag-time of 4 ns was used for counting transitions between states and transition probabilities were computed using row-normalization with a prior counts.

### Pseudovirus neutralization

Vero E6 or TMPRSS2-Vero E6 recombinant cells were seeded in 384 well tissue culture plates at a density of 3,500 cells per well in 20 µl of stimulation medium and incubated at 37°C, 5% CO_2_, for 2-4 hours. In parallel, antibodies were serially diluted and incubated with diluted pseudovirus at a desired multiplicity of infection ([MOI] 0.25 for SARS-CoV-2 pre-Omicron variants and non-SARS-CoV-2 sarbecoviruses; 0.5 for SARS-CoV-2 Omicron variants). Antibody neutralization was assessed with 12-point titration curves in technical quadruplicate with four-fold serial dilutions starting at 18 µg/ml (final concentration in the tissue culture plate).

For combination experiments, antibodies were mixed at a 1:1 ratio and anti-spike antibodies tested as single agents were combined with isotype control at the same concentrations to account for overall mass of anti-spike antibodies tested as combinations. For selected experiments (**Figures S1C, S3C-E, S4A and S5**), antibody neutralization was assessed with 8-point titration curves in technical duplicate with four-fold serial dilutions starting at 4.5 µg/ml (final concentration in the tissue culture plates). After 30-60 minutes of incubation at 37°C, 5% CO_2_, 20 µl of pseudovirus/antibody mixture was added to the pre-seeded tissue culture plates, achieving final antibody concentrations in a total volume of 40 µl. After 24 hours at 37°C, 5% CO_2_, an equal volume of luciferase substrate (BPS Bioscience Inc, 60690-3) was added directly to culture plates and luminescence was quantified on the EnVision plate reader (PerkinElmer, serial #411177671). Percentage neutralization was calculated with the following formula, where “signal_positiveControl_” is defined by the average luminescence signal of wells containing cells without pseudovirus, “signal_negativeContro_l” is defined as the average luminescence signal of wells containing cells with pseudovirus, “signal_well_” is defined as the average luminescence signal of wells containing cells with both antibody and pseudovirus:

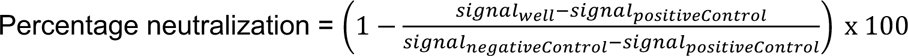

### Live virus neutralization

Live virus neutralization experiments were performed at Virology Research Services Ltd (Sittingbourne, United Kingdom). Vero cells were detached and counted, diluted in complete media and seeded at 8,000 cells per 100 µl. After seeding, plates were incubated at 37°C, 5% CO_2_ until the following day. Antibody neutralization was assessed with 12-point titration curves in technical triplicate with four-fold serial dilutions starting at 18 µg/ml (final concentration in the tissue culture plate). For combination conditions, antibodies were mixed at a 1:1 ratio and anti-spike antibodies tested as single agents were combined with isotype control at the same concentrations to account for overall mass of anti-spike antibodies tested as combinations. A predetermined MOI of each SARS-CoV-2 variant, required to obtain between 10% and 20% infection, was added to each well of the antibody dilutions and in the infected untreated control wells. Media only (without virus) was added to the uninfected untreated control wells. After 1 hour, media was removed from the cells and replaced with the virus-antibody dilution mixtures.

Plates were incubated for 6 hours at 37°C and 5% CO_2_, and the virus-antibodies mixture was left on cells for the duration of the experiment. After 6 hours, the infection plates were washed with PBS, fixed for 30 minutes with 4% formaldehyde, washed again with PBS, and stored until staining. Cells were immunostained by quenching residual formaldehyde with 50 mM ammonium chloride, after which cells were permeabilized (0.1% Triton X-100) and labelled with an antibody recognizing SARS-CoV-2 nucleocapsid (GeneTex, HL488). Primary antibody was detected with an Alexa-488 conjugate secondary antibody (Life Technologies, AA11034), and nuclei were stained with Hoechst. Images were acquired on a CellInsight CX5 high content platform (Thermo Scientific) using a 4X objective, and percentage infection calculated using CellInsight CX5 software (infected cells/total cells x 100). Normalized percentages of inhibition were calculated using the following formula, where “% infection sample” is the percentage infection of any given dilution of the test article, “% infection uninfected control” is the percentage infection measured in untreated wells that have not been infected (background signal), and “% infection infected control” is the percentage infection in untreated wells that have been infected with the same amount of virus as the test articles (maximum infection):

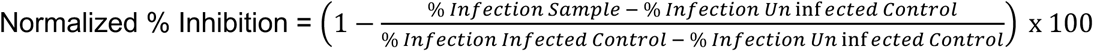

### Identification of viral escape *in vitro* through resistance passaging

Escape studies as well as EC50 and EC90 determinations were performed at Virology Research Services Ltd (Sittingbourne, United Kingdom). First, the EC50 and EC90 of each antibody or antibody combination therapy was determined. Vero E6 cells were seeded at 8,000 cells per well into a 96 well plate in complete media and infected with SARS-CoV-2 Omicron BA.1 (MOI 0.01) following pre-incubation with different antibody concentrations in a 12-point titration curve, 4-fold dilution series with the final top concentration at 18 µg/ml, in technical triplicate for 2 hours at 37°C, 5% CO_2_. Positive and negative control antibodies were included as assay controls. At 72 hours, the neutralization activity was determined using an MTT assay to measure antiviral activity via cytopathic effect. Briefly, 20 µl of MTT reagent (Sigma, M5655) was added to cells for 2 hours at 37°C, 5% CO_2_. Media was then removed and the precipitate solubilized with 50 µl of a 1:1 Isopropanol:DMSO mixture for 20 minutes. The supernatant was then transferred to a clean plate and the signal read at 570 nm. The EC50 and EC90 of each antibody or antibody combination was calculated. The EC50 values were extrapolated from curves representing the best fit (non-linear regression analysis, variable slope) of the logarithm of antibody concentration versus the normalized percentages of viability using GraphPad Prism 9.0 software. To assess the escape of SARS-CoV-2 Omicron BA.1 from GB-0669, PRO-37587 antibodies individually and in combination, Vero E6 cells were plated at 55,000 cells per well in 24 well plates. The EC50, EC90, and 3x the EC90 of the antibodies singly or in combination (1:1) were added to SARS-CoV-2 Omicron BA.1 at an MOI of 0.01 for 1 hour at 37°C, 5% CO_2_. The antibody-virus mixture was added to the cells for 2 hours at 37°C, 5% CO_2_. Following infection, fresh media was added keeping the final antibody concentration consistent. The plates were incubated for 72 hours at 37°C, 5% CO_2_ and cytopathic effect determined (passage 1). Each condition was completed in sextuplicate. Once cytopathic effect was visible virus supernatant was harvested and pooled from the replicate wells. To quantify infectious virus in pooled supernatants, TCID50 was determined in Vero E6 cells seeded at 8,000 cells per well in complete media. A 10-fold, 8-point virus dilution series was performed and used to infect cells. 72 hours later, TCID50 titer was determined. Virus harvested from the highest antibody concentration, with a sufficient titer to infected cells at an MOI of 0.01, was selected for the next passage. The lowest test antibody concentration was then replaced by the same concentration from which the virus was harvest from. The middle and highest antibody concentrations were adjusted to be three times higher than the lowest test antibody concentration, resulting in an overall increase in the antibody concentration. Passaging was continued until complete escape was observed after monoclonal or combination antibody treatment at a concentration of > 10 µg/ml or until 10 passages was reached. To determine if there was any change in sensitivity to the antibodies, the EC50 final resistance passage virus was assessed using a CPE-based microneutralization assay as described above.

### Viral RNA extraction and sequencing

Once viral escape was evident as determined by CPE, cell-cleared supernatant samples from resistance passaging with the monoclonal or combination antibody treatments were extracted for RNA. The isolated RNA was subjected to RT-qPCR to determine viral load using the Luna Universal One-Step RT-qPCR kit (New England Biolabs, E3005L) with primers to SARS-CoV-2 N1. The *spike* gene was amplified and sequenced by Cerba Research (formerly Viroclinics-DDL, Rijswijk, Netherlands) utilizing the NextSeq 1000 Illumina platform (Illumina, serial number: VL00351). Amplification of the full-length *spike* gene was performed using a selection of primers from the ARTIC SARS-CoV-2 whole genome primer sets covering the entirety of the spike gene, SARS-CoV-2 v4 scheme: https://community.artic.network/t/sars-cov-2-version-4-scheme-release/312, with modifications to improve the amplification of Omicron variant spike gene mutations (Spike ARTIC v4.2). The NGS analytics system available at Cerba Research was used to assess quality control as well as perform primer trimming and demultiplexing providing the sequences used for variant calling analysis.^75,76^

### Variant calling analysis

A computational workflow was developed to identify spike protein mutations associated with antibody evasion from each molecule or the combination. The sequenced samples corresponded to unique combinations of antibody, concentration, and passage number. First, the reads were mapped to the SARS-CoV-2 Omicron BA.1.18 reference genome (GISAID accession EPI_ISL_7605630) using BWA-MEM2 (version 2.2.1).^77^ Next, FreeBayes (version v1.3.7) was used to identify small polymorphisms in each sample relative to the reference.^78^ For each polymorphism and each sample, an alternate allele frequency was calculated as the proportion of reads harboring the mutant allele. The putative escape mutations were defined to be those that were not present in the starting stock and showed increasing allele frequency with drug concentration or passage number. Finally, each identified DNA polymorphism was translated to its corresponding amino acid substitution.

### Viral growth of resistance passaged virus

Multistep growth curves were performed in duplicate by inoculating 260,000 Vero E6 cells at an MOI of 0.005 in the absence of antibody treatment. Supernatants were harvested and clarified by centrifugation at 6 h, 16 h, 24 h, 32 h, 48 h, and 72 h. The amount of infectious virus in the cleared supernatant was quantified using a TCID50 assay. Briefly, eight 10-fold dilutions of the harvested viruses (starting from 10^-1^) were added to monolayers of Vero E6 cells seeded at 8,000 cells/well in a 96 well plate format. After 96 h, the cells were fixed with 4% formaldehyde in 1x PBS and stained with crystal violet. The titers (TCD50/ml) at each timepoint were determined using the Reed-Muench method.

### Identification of relative frequency of mutations in the GB-0669 and PRO-37587 target region sequences

The GISAID database was queried to determine the frequency of our putative escape mutations across deposited SARS-CoV-2 sequences. First, global sequence alignment was used to map coordinates relative to the reference spike sequence (EPI_ISL_7605630) to the coordinates used by the GISAID database (the original Wuhan sequence EPI_ISL_402124); the mutations of interest were then converted to GISAID reference-relative representations. Next, the GISAID data was accessed through CovSPECTRUM via the LAPIS API.^79,80^ For each mutation, we fetched the proportion of deposited sequences harboring the mutation in GISAID over two-time intervals: March 2023 – March 2024, and January 2020 – March 2024. All sequences harboring any mutations at both PRO-35787-associated positions were also fetched, confirming that the combination of the two putative escape alleles has never been observed.

### Hamster SARS-CoV-2 Omicron BA.2 challenge

Hamster challenge experiments were performed at BIOQUAL (Rockville, USA). A total of 60 male Golden Syrian Hamsters (10 groups, 6 animals per group) were intranasally challenged at day 0 with 2.8x10^3^ TCID50/Dose of SARS-CoV-2 Omicron BA.2 variant (BEI Resources, NR-56522). On day -1, animals were intraperitoneally administered isotype control (isotype, palivizumab [anti-RSV F mAb] variable regions expressed as human IgG1 with LS mutation in the Fc region), positive controls (bebtelovimab; sotrovimab VH/VL - huIgG1-LS, sotrovimab variable regions expressed as human IgG1 with LS mutation in the Fc region), GB-0669, or PRO-37587. All single agents were administered at 12 mg/kg, except for the isotype, which was administered at 60 mg/kg. GB-0669 and PRO-37587 in combination were administered in a 1:1 ratio at either 0.75, 1.5, 3, 6, or 12 mg/kg per antibody. Lung samples were collected on day 4, with the right lung sectioned for RT-qPCR assay (genomic and subgenomic RNA), TCID50 assay, and homogenization in QIAzol (Qiagen, 73906), and the left lung fixed in 10% neutral buffered formalin for histopathology.

### hACE2 AC70 mouse SARS-CoV-2 Omicron XBB.1.5 challenge

Mouse challenge experiments were performed at BIOQUAL (Rockville, USA). A total of 70 female hACE2 AC70 mice (7 groups, 10 animals per group) were intranasally challenged at day 0 with 1.96 x 10^4^ PFU/Dose of SARS-CoV-2 XBB.1.5 variant (BEI Resources, NR-59105). On day -1, animals were intraperitoneally administered isotype control (isotype, palivizumab [anti-RSV F mAb] variable regions expressed as human IgG1 with LS mutation in the Fc region), positive control (pemivibart), GB-0669, PRO-37587 or a combination of GB-0669 and PRO-37587. GB-0669 was administered at doses of 7.5, 15 and 30 mg/kg as a single agent. PRO-37857 and pemivibart were administered at a dose of 7.5 mg/kg as a single agent. The combination of GB-0669 and PRO-37857 was administered at a dose of 7.5 mg/kg for each antibody. Palivizumab was administered at a dose of 30 mg/kg. Lung samples from 5 animals of each group were collected on day 4, sectioned and snap-frozen for RT-qPCR (genomic and subgenomic RNA) and TCID50 assays. The remaining 5 animals from each group were allowed to proceed to study endpoint of day 14 unless a body weight loss of >20% occurred at which time the animal was euthanized. During the study period, daily body weights and clinical observations were monitored.

### Quantification and statistical analysis

Statistical analysis was performed with Prism 9.5.0 software. Half maximal effective concentration (EC50) values and 95% confidence intervals were derived for each Log-transformed antibody titration curve using log(inhibitor) vs. response -- Variable slope (four parameters) equation. 90% maximal effective concentration (EC90) values and 95% confidence intervals were derived for each Log-transformed antibody titration curve using log(agonist) vs. response -- Find ECanything equation by setting F parameter to 90. Maximal neutralization (Emax %) values and 95% confidence intervals were defined as percentage neutralization at top concentration (18,000 ng/ml). Area under the curve (AUC) values derived for each Log-transformed antibody titration curve setting baseline Y = 0 and ignoring any peaks below this baseline. When necessary, data were Log-transformed before statistical analysis to approximate normal distributions. Statistical differences between groups in datasets with one categorical variable and more than 2 groups were evaluated by one-way ANOVA corrected for multiple comparisons. Statistical differences between groups in datasets with two categorical variables were evaluated by two-way ANOVA corrected for multiple comparisons. ^*^ and ^**^ respectively indicate p ≤ 0.05 and 0.01.

## RESOURCE AVAILABILITY

### Lead Contact

Requests for further information, resources, and reagents should be directed to the lead contact, Francesco Borriello.

### Materials availability

Materials described in this manuscript may be made available to qualified, academic, noncommercial researchers through a materials transfer agreement upon request. Novel antibody sequences reported in this study are from the structures deposited in PDB.

### Data and code availability

The structures are available from the PDB: 9NFT (PRO-37587 bound to SARS-CoV-2 Omicron BA.1 spike), 9NHP (GB-0669 bound to S2 stem helix). Cryo-EM density maps are available from the EMDB: EMD-49461 (GB-0669 and PRO-37587 bound to SARS-CoV-2 Omicron BA.1 spike), EMD-49372 (PRO-37587 bound to SARS-CoV-2 Omicron BA.1 spike composite map), EMD-49566 (PRO-37587 bound to SARS-CoV-2 Omicron BA.1 spike consensus map), and EMD-49567 (PRO-37587 bound to SARS-CoV-2 Omicron BA.1 spike local refinement).

Illumina-based deep sequencing data from the resistance passaging experiments are available from the NCBI Sequence Read Archive (SRA), BioProject: PRJNA1234183, BioSample Accession Numbers: SAMN47134171-SAMN47134194.

## ACKNOWLEDGEMENTS

We thank former and current Generate Biomedicines employees Robin Green, Babu R.P. Saravanan, Benjamin Manning, Ramsey Fadlalla, Adi Ravipati and Alexandra Gurney for supporting high-throughput protein production workflows; Brian Patuto, Avery Farnham, Sandra Sipsey, Nathan Wong and Darren Ferguson for supporting large scale production workflows; Atreyi Bhattacharya, Lyle McPherson, Alex Ayoub, Patrick Brophy and Mark Benhaim for supporting developability assessment workflows; Danielle Curran for expression of samples used for Cryo-EM, Charleen Zhao for vitrification of Cryo-EM grids; Denise Murphy for project management support.

## AUTHOR CONTRIBUTIONS

Conceived research: G.C.M., K.P.S., M.I.Z., A.H.R., D.H. and F.B.; SPR experiments: G.C.M. and M.Z.M.; virology experiments: G.C.M, K.P.S., G.S., H.L., E.C., V.V. and H.T.; DELFIA experiments: G.C.M. and G.S.; Cryo-EM experiments and data analysis: M.I.Z., A.J., C.L.M.P., E.P. and J.D.G.; viral escape experiments, sequencing and analysis: G.C.M., K.P.S., B.H., P.R. and J.T.; yeast display experiments: R.S.F., A.L. and N.L.V.; antibody production and analytics: N.M.S. and N.J.; crystallography experiments and data analysis: D.M.L., M.I.Z. and E.P.; ML-guided protein optimization: A.H.R., Z.K., V.F., and G.G.; oversight and discussion of wet lab activities and data: K.H., A.P., G.C.K.W.K., H.V.E., A.R., D.H. and F.B.; oversight and discussion of dry lab activities and data: G.G.; wrote the manuscript: G.C.M., K.P.S., M.I.Z., D.H. and F.B; all the authors have reviewed and contributed to the manuscript.

## DECLARATION OF INTERESTS

At the time this work was performed, all authors were employees and shareholders of Generate Biomedicines.

## SUPPLEMENTAL INFORMATION

**Document S1. Figures S1-S8 and Tables S1-S6.**

**Video S1, Interpolations between Cryo-EM derived conformational states of PRO-37587 bound to SARS-CoV-2 spike protein.**

Molecular model of the SARS-CoV-2 spike protein (surface representation with chains colored blue, gray, and green) binding to PRO-37587 (cartoon representation with heavy and light chains colored light pink and purple) and subsequent conformational heterogeneity. The first frames depict the conformational change between the down (PDB: 6VXX) and open conformation (PDB: 9NFT). The second part of the video depicts the conformational changes of the RBD once bound to PRO-37587, which was generated by individually docking PRO-37587 and the SARS-CoV-2 RBD into multiple Cryo-EM density maps derived from a 3D classification in CryoSPARC.

## REFERENCES

1. Drosten, C., Günther, S., Preiser, W., Werf, S. van der, Brodt, H.-R., Becker, S., Rabenau, H., Panning, M., Kolesnikova, L., Fouchier, R.A.M., et al. (2003). Identification of a Novel Coronavirus in Patients with Severe Acute Respiratory Syndrome. N. Engl. J. Med. 348, 1967– 1976. 10.1056/nejmoa030747.

2. Zhong, N., Zheng, B., Li, Y., Poon, L., Xie, Z., Chan, K., Li, P., Tan, S., Chang, Q., Xie, J., et al. (2003). Epidemiology and cause of severe acute respiratory syndrome (SARS) in Guangdong, People’s Republic of China, in February, 2003. Lancet (Lond., Engl.) 362, 1353–1358. 10.1016/s0140-6736(03)14630-2.

3. Tu, C., Crameri, G., Kong, X., Chen, J., Sun, Y., Yu, M., Xiang, H., Xia, X., Liu, S., Ren, T., et al. (2004). Antibodies to SARS Coronavirus in Civets. Emerg. Infect. Dis. 10, 2244–2248. 10.3201/eid1012.040520.

4. Wang, L.F., and Eaton, B.T. (2007). Bats, civets and the emergence of SARS. Curr. Top. Microbiol. Immunol. 315, 325–344. 10.1007/978-3-540-70962-6_13.

5. Zhou, P., Yang, X.-L., Wang, X.-G., Hu, B., Zhang, L., Zhang, W., Si, H.-R., Zhu, Y., Li, B., Huang, C.-L., et al. (2020). A pneumonia outbreak associated with a new coronavirus of probable bat origin. Nature 579, 270–273. 10.1038/s41586-020-2012-7.

6. Wu, J.T., Leung, K., and Leung, G.M. (2020). Nowcasting and forecasting the potential domestic and international spread of the 2019-nCoV outbreak originating in Wuhan, China: a modelling study. Lancet 395, 689–697. 10.1016/s0140-6736(20)30260-9.

7. Viruses, C.S.G. of the I.C. on T. of, Gorbalenya, A.E., Baker, S.C., Baric, R.S., Groot, R.J. de, Drosten, C., Gulyaeva, A.A., Haagmans, B.L., Lauber, C., Leontovich, A.M., et al. (2020). The species Severe acute respiratory syndrome-related coronavirus: classifying 2019-nCoV and naming it SARS-CoV-2. Nat. Microbiol. 5, 536–544. 10.1038/s41564-020-0695-z.

8. Wilder-Smith, A., Chiew, C.J., and Lee, V.J. (2020). Can we contain the COVID-19 outbreak with the same measures as for SARS? Lancet Infect. Dis. 20, e102–e107. 10.1016/s1473-3099(20)30129-8.

9. Muralidar, S., Ambi, S.V., Sekaran, S., and Krishnan, U.M. (2020). The emergence of COVID-19 as a global pandemic: Understanding the epidemiology, immune response and potential therapeutic targets of SARS-CoV-2. Biochimie 179, 85–100. 10.1016/j.biochi.2020.09.018.

10. Focosi, D., Franchini, M., Casadevall, A., and Maggi, F. (2024). An update on the anti-spike monoclonal antibody pipeline for SARS-CoV-2. Clin. Microbiol. Infect. 30, 999–1006. 10.1016/j.cmi.2024.04.012.

11. Roemer, C., Sheward, D.J., Hisner, R., Gueli, F., Sakaguchi, H., Frohberg, N., Schoenmakers, J., Sato, K., O’Toole, Á., Rambaut, A., et al. (2023). SARS-CoV-2 evolution in the Omicron era. Nat. Microbiol. 8, 1952–1959. 10.1038/s41564-023-01504-w.

12. Carabelli, A.M., Peacock, T.P., Thorne, L.G., Harvey, W.T., Hughes, J., Silva, T.I. de, Peacock, S.J., Barclay, W.S., Silva, T.I. de, Towers, G.J., et al. (2023). SARS-CoV-2 variant biology: immune escape, transmission and fitness. Nat. Rev. Microbiol. 21, 162–177. 10.1038/s41579-022-00841-7.

13. Gruell, H., Vanshylla, K., Weber, T., Barnes, C.O., Kreer, C., and Klein, F. (2022). Antibody-mediated neutralization of SARS-CoV-2. Immunity 55, 925–944. 10.1016/j.immuni.2022.05.005.

14. Evans, T.S., Tan, C.W., Aung, O., Phyu, S., Lin, H., Coffey, L.L., Toe, A.T., Aung, P., Aung, T.H., Aung, N.T., et al. (2023). Exposure to diverse sarbecoviruses indicates frequent zoonotic spillover in human communities interacting with wildlife. Int. J. Infect. Dis. 131, 57–64. 10.1016/j.ijid.2023.02.015.

15. Sánchez, C.A., Li, H., Phelps, K.L., Zambrana-Torrelio, C., Wang, L.-F., Zhou, P., Shi, Z.-L., Olival, K.J., and Daszak, P. (2022). A strategy to assess spillover risk of bat SARS-related coronaviruses in Southeast Asia. Nat. Commun. 13, 4380. 10.1038/s41467-022-31860-w.

16. Menachery, V.D., Yount, B.L., Sims, A.C., Debbink, K., Agnihothram, S.S., Gralinski, L.E., Graham, R.L., Scobey, T., Plante, J.A., Royal, S.R., et al. (2016). SARS-like WIV1-CoV poised for human emergence. Proc. Natl. Acad. Sci. 113, 3048–3053. 10.1073/pnas.1517719113.

17. Corti, D., Purcell, L.A., Snell, G., and Veesler, D. (2021). Tackling COVID-19 with neutralizing monoclonal antibodies. Cell 184, 3086–3108. 10.1016/j.cell.2021.05.005.

18. Li, C.-J., and Chang, S.-C. (2023). SARS-CoV-2 spike S2-specific neutralizing antibodies. Emerg. Microbes Infect. 12, 2220582. 10.1080/22221751.2023.2220582.

19. Barnes, C.O., Jette, C.A., Abernathy, M.E., Dam, K.-M.A., Esswein, S.R., Gristick, H.B., Malyutin, A.G., Sharaf, N.G., Huey-Tubman, K.E., Lee, Y.E., et al. (2020). SARS-CoV-2 neutralizing antibody structures inform therapeutic strategies. Nature 588, 682–687. 10.1038/s41586-020-2852-1.

20. Romero, P.A., and Arnold, F.H. (2009). Exploring protein fitness landscapes by directed evolution. Nat. Rev. Mol. Cell Biol. 10, 866–876. 10.1038/nrm2805.

21. Rappazzo, C.G., Tse, L.V., Kaku, C.I., Wrapp, D., Sakharkar, M., Huang, D., Deveau, L.M., Yockachonis, T.J., Herbert, A.S., Battles, M.B., et al. (2021). Broad and potent activity against SARS-like viruses by an engineered human monoclonal antibody. Sci. (N. York, Ny) 371, 823– 829. 10.1126/science.abf4830.

22. Hie, B.L., Shanker, V.R., Xu, D., Bruun, T.U.J., Weidenbacher, P.A., Tang, S., Wu, W., Pak, J.E., and Kim, P.S. (2024). Efficient evolution of human antibodies from general protein language models. Nat. Biotechnol. 42, 275–283. 10.1038/s41587-023-01763-2.

23. Tortorici, M.A., Czudnochowski, N., Starr, T.N., Marzi, R., Walls, A.C., Zatta, F., Bowen, J.E., Jaconi, S., Iulio, J.D., Wang, Z., et al. (2021). Broad sarbecovirus neutralization by a human monoclonal antibody. Nature 597, 103–108. 10.1038/s41586-021-03817-4.

24. Jennewein, M.F., MacCamy, A.J., Akins, N.R., Feng, J., Homad, L.J., Hurlburt, N.K., Seydoux, E., Wan, Y.-H., Stuart, A.B., Edara, V.V., et al. (2021). Isolation and characterization of cross-neutralizing coronavirus antibodies from COVID-19+ subjects. Cell Rep. 36, 109353. 10.1016/j.celrep.2021.109353.

25. Hurlburt, N.K., Homad, L.J., Sinha, I., Jennewein, M.F., MacCamy, A.J., Wan, Y.-H., Boonyaratanakornkit, J., Sholukh, A.M., Jackson, A.M., Zhou, P., et al. (2022). Structural definition of a pan-sarbecovirus neutralizing epitope on the spike S2 subunit. Commun. Biol. 5, 342. 10.1038/s42003-022-03262-7.

26. Li, W., Chen, Y., Prévost, J., Ullah, I., Lu, M., Gong, S.Y., Tauzin, A., Gasser, R., Vézina, D., Anand, S.P., et al. (2022). Structural basis and mode of action for two broadly neutralizing antibodies against SARS-CoV-2 emerging variants of concern. Cell Rep. 38, 110210. 10.1016/j.celrep.2021.110210.

27. Wang, Q., Guo, Y., Iketani, S., Nair, M.S., Li, Z., Mohri, H., Wang, M., Yu, J., Bowen, A.D., Chang, J.Y., et al. (2022). Antibody evasion by SARS-CoV-2 Omicron subvariants BA.2.12.1, BA.4 and BA.5. Nature 608, 603–608. https://10.1038/s41586-022-05053-w.

28. Liu, L., Iketani, S., Guo, Y., Chan, J.F.-W., Wang, M., Liu, L., Luo, Y., Chu, H., Huang, Y., Nair, M.S., et al. (2022). Striking antibody evasion manifested by the Omicron variant of SARS-CoV-2. Nature 602, 676–681. 10.1038/s41586-021-04388-0.

29. Grunst, M.W., Qin, Z., Dodero-Rojas, E., Ding, S., Prévost, J., Chen, Y., Hu, Y., Pazgier, M., Wu, S., Xie, X., et al. (2024). Structure and inhibition of SARS-CoV-2 spike refolding in membranes. Science 385, 757–765. 10.1126/science.adn5658.

30. Zalevsky, J., Chamberlain, A.K., Horton, H.M., Karki, S., Leung, I.W.L., Sproule, T.J., Lazar, G.A., Roopenian, D.C., and Desjarlais, J.R. (2010). Enhanced antibody half-life improves in vivo activity. Nat. Biotechnol. 28, 157–159. 10.1038/nbt.1601.

31. Husic, B.E., and Pande, V.S. (2018). Markov State Models: From an Art to a Science. J. Am. Chem. Soc. 140, 2386–2396. 10.1021/jacs.7b12191.

32. Starr, T.N., Greaney, A.J., Hilton, S.K., Ellis, D., Crawford, K.H.D., Dingens, A.S., Navarro, M.J., Bowen, J.E., Tortorici, M.A., Walls, A.C., et al. (2020). Deep Mutational Scanning of SARS-CoV-2 Receptor Binding Domain Reveals Constraints on Folding and ACE2 Binding. Cell 182, 1295–1310.e20. 10.1016/j.cell.2020.08.012.

33. Zimmerman, M.I., Porter, J.R., Ward, M.D., Singh, S., Vithani, N., Meller, A., Mallimadugula, U.L., Kuhn, C.E., Borowsky, J.H., Wiewiora, R.P., et al. (2021). SARS-CoV-2 simulations go exascale to predict dramatic spike opening and cryptic pockets across the proteome. Nat. Chem. 13, 651–659. 10.1038/s41557-021-00707-0.

34. Zondlo, N.J. (2013). Aromatic–Proline Interactions: Electronically Tunable CH/π Interactions. Acc. Chem. Res. 46, 1039–1049. 10.1021/ar300087y.

35. Huang, K.-Y.A., Chen, X., Mohapatra, A., Nguyen, H.T.V., Schimanski, L., Tan, T.K., Rijal, P., Vester, S.K., Hills, R.A., Howarth, M., et al. (2023). Structural basis for a conserved neutralization epitope on the receptor-binding domain of SARS-CoV-2. Nat. Commun. 14, 311. 10.1038/s41467-023-35949-8.

36. Iketani, S., Liu, L., Guo, Y., Liu, L., Chan, J.F.-W., Huang, Y., Wang, M., Luo, Y., Yu, J., Chu, H., et al. (2022). Antibody evasion properties of SARS-CoV-2 Omicron sublineages. Nature 604, 553–556. 10.1038/s41586-022-04594-4.

37. Baum, A., Fulton, B.O., Wloga, E., Copin, R., Pascal, K.E., Russo, V., Giordano, S., Lanza, K., Negron, N., Ni, M., et al. (2020). Antibody cocktail to SARS-CoV-2 spike protein prevents rapid mutational escape seen with individual antibodies. Sci. (N. York, Ny) 369, eabd0831. 10.1126/science.abd0831.

38. Copin, R., Baum, A., Wloga, E., Pascal, K.E., Giordano, S., Fulton, B.O., Zhou, A., Negron, N., Lanza, K., Chan, N., et al. (2021). The monoclonal antibody combination REGEN-COV protects against SARS-CoV-2 mutational escape in preclinical and human studies. Cell 184, 3949–3961.e11. 10.1016/j.cell.2021.06.002.

39. Meulen, J. ter, Brink, E.N. van den, Poon, L.L.M., Marissen, W.E., Leung, C.S.W., Cox, F., Cheung, C.Y., Bakker, A.Q., Bogaards, J.A., Deventer, E. van, et al. (2006). Human Monoclonal Antibody Combination against SARS Coronavirus: Synergy and Coverage of Escape Mutants. PLoS Med. 3, e237. 10.1371/journal.pmed.0030237.

40. Wang, L., Shi, W., Chappell, J.D., Joyce, M.G., Zhang, Y., Kanekiyo, M., Becker, M.M., Doremalen, N. van, Fischer, R., Wang, N., et al. (2018). Importance of Neutralizing Monoclonal Antibodies Targeting Multiple Antigenic Sites on the Middle East Respiratory Syndrome Coronavirus Spike Glycoprotein To Avoid Neutralization Escape. J. Virol. 92. 10.1128/jvi.02002-17.

41. Zost, S.J., Gilchuk, P., Case, J.B., Binshtein, E., Chen, R.E., Nkolola, J.P., Schäfer, A., Reidy, J.X., Trivette, A., Nargi, R.S., et al. (2020). Potently neutralizing and protective human antibodies against SARS-CoV-2. Nature 584, 443–449. 10.1038/s41586-020-2548-6.

42. Low, J.S., Jerak, J., Tortorici, M.A., McCallum, M., Pinto, D., Cassotta, A., Foglierini, M., Mele, F., Abdelnabi, R., Weynand, B., et al. (2022). ACE2-binding exposes the SARS-CoV-2 fusion peptide to broadly neutralizing coronavirus antibodies. Science 377, eabq2679. 10.1126/science.abq2679.

43. Sun, X., Yi, C., Zhu, Y., Ding, L., Xia, S., Chen, X., Liu, M., Gu, C., Lu, X., Fu, Y., et al. (2022). Neutralization mechanism of a human antibody with pan-coronavirus reactivity including SARS-CoV-2. Nat. Microbiol. 7, 1063–1074. 10.1038/s41564-022-01155-3.

44. Mast, F.D., Fridy, P.C., Ketaren, N.E., Wang, J., Jacobs, E.Y., Olivier, J.P., Sanyal, T., Molloy, K.R., Schmidt, F., Rutkowska, M., et al. (2021). Highly synergistic combinations of nanobodies that target SARS-CoV-2 and are resistant to escape. eLife 10, e73027. 10.7554/elife.73027.

45. Piepenbrink, M.S., Park, J.-G., Deshpande, A., Loos, A., Ye, C., Basu, M., Sarkar, S., Khalil, A.M., Chauvin, D., Woo, J., et al. (2022). Potent universal beta-coronavirus therapeutic activity mediated by direct respiratory administration of a Spike S2 domain-specific human neutralizing monoclonal antibody. PLoS Pathog. 18, e1010691. 10.1371/journal.ppat.1010691.

46. Planchais, C., Fernández, I., Chalopin, B., Bruel, T., Rosenbaum, P., Beretta, M., Dimitrov, J.D., Conquet, L., Donati, F., Prot, M., et al. (2024). Broad sarbecovirus neutralization by combined memory B cell antibodies to ancestral SARS-CoV-2. iScience 27, 110354. 10.1016/j.isci.2024.110354.

47. Wang, Q., Iketani, S., Li, Z., Liu, L., Guo, Y., Huang, Y., Bowen, A.D., Liu, M., Wang, M., Yu, J., et al. (2023). Alarming antibody evasion properties of rising SARS-CoV-2 BQ and XBB subvariants. Cell 186, 279–286.e8. 10.1016/j.cell.2022.12.018.

48. Muñoz-Fontela, C., Widerspick, L., Albrecht, R.A., Beer, M., Carroll, M.W., Wit, E. de, Diamond, M.S., Dowling, W.E., Funnell, S.G.P., García-Sastre, A., et al. (2022). Advances and gaps in SARS-CoV-2 infection models. PLoS Pathog. 18, e1010161. 10.1371/journal.ppat.1010161.

49. Bao, L., Deng, W., Huang, B., Gao, H., Liu, J., Ren, L., Wei, Q., Yu, P., Xu, Y., Qi, F., et al. (2020). The pathogenicity of SARS-CoV-2 in hACE2 transgenic mice. Nature 583, 830–833. 10.1038/s41586-020-2312-y.

50. Clinicaltrials.gov (2024). Efficacy and Safety of the Anti-COVID-19 Antibody SA55 for Injection in Patients With Hematological Disorders Who Are Persistently Positive for COVID-19 (NCT05675943). https://clinicaltrials.gov/study/NCT05675943?term=SA55&rank=4.

51. Clinicaltrials.gov (2024). SA55 Injection Phase II Study in the Treatment of Mild/Moderate COVID-19 Patients (NCT06042764). https://clinicaltrials.gov/study/NCT06042764?term=SA55&rank=1.

52. Clinicaltrials.gov (2024). SA55 Injection: a Potential Therapy for the Prevention and Treatment of COVID-19 (NCT06050460). https://clinicaltrials.gov/study/NCT06050460?term=SA55&rank=2.

53. Clinicaltrials.gov (2024). SA55 Novel Coronavirus Broad-spectrum Neutralizing Antibody Nasal Spray in Health People (NCT06048393). https://clinicaltrials.gov/study/NCT06048393?term=SA55&rank=3.

54. Nutalai, R., Zhou, D., Tuekprakhon, A., Ginn, H.M., Supasa, P., Liu, C., Huo, J., Mentzer, A.J., Duyvesteyn, H.M.E., Dijokaite-Guraliuc, A., et al. (2022). Potent cross-reactive antibodies following Omicron breakthrough in vaccinees. Cell 185, 2116–2131.e18. 10.1016/j.cell.2022.05.014.

55. Cai, Y., Diallo, S., Rosenthal, K., Ren, K., Flores, D.J., Dippel, A., Oganesyan, V., Dyk, N. van, Chen, X., Cantu, E., et al. (2024). AZD3152 neutralizes SARS-CoV-2 historical and contemporary variants and is protective in hamsters and well tolerated in adults. Sci. Transl. Med. 16, eado2817. 10.1126/scitranslmed.ado2817.

56. Planas, D., Staropoli, I., Planchais, C., Yab, E., Jeyarajah, B., Rahou, Y., Prot, M., Guivel-Benhassine, F., Lemoine, F., Enouf, V., et al. (2024). Escape of SARS-CoV-2 variants KP1.1, LB.1 and KP3.3 from approved monoclonal antibodies. bioRxiv, 2024.08.20.608835. 10.1101/2024.08.20.608835.

57. Wang, Q., Guo, Y., Ho, J., and Ho, D.D. (2024). Pemivibart is less active against recent SARS-CoV-2 JN.1 sublineages. bioRxiv, 2024.08.12.607496. 10.1101/2024.08.12.607496.

58. Rosen, L.E., Tortorici, M.A., Marco, A.D., Pinto, D., Foreman, W.B., Taylor, A.L., Park, Y.-J., Bohan, D., Rietz, T., Errico, J.M., et al. (2024). A potent pan-sarbecovirus neutralizing antibody resilient to epitope diversification. Cell. 10.1016/j.cell.2024.09.026.

59. Tamura, T., Mizuma, K., Nasser, H., Deguchi, S., Padilla-Blanco, M., Oda, Y., Uriu, K., Tolentino, J.E.M., Tsujino, S., Suzuki, R., et al. (2024). Virological characteristics of the SARS-CoV-2 BA.2.86 variant. Cell Host Microbe 32, 170–180.e12. 10.1016/j.chom.2024.01.001.

60. Tsujino, S., Deguchi, S., Nomai, T., Padilla-Blanco, M., Plianchaisuk, A., Wang, L., Begum, M.M., Uriu, K., Mizuma, K., Nao, N., et al. (2024). Virological characteristics of the SARS-CoV-2 Omicron EG.5.1 variant. Microbiol. Immunol. 10.1111/1348-0421.13165.

61. Kaku, Y., Yo, M.S., Tolentino, J.E., Uriu, K., Okumura, K., Consortium, T.G. to P.J. (G2P-J., Ito, J., and Sato, K. (2024). Virological characteristics of the SARS-CoV-2 KP.3, LB.1, and KP.2.3 variants. Lancet Infect. Dis. 24, e482–e483. 10.1016/s1473-3099(24)00415-8.

62. Ingraham, J.B., Baranov, M., Costello, Z., Barber, K.W., Wang, W., Ismail, A., Frappier, V., Lord, D.M., Ng-Thow-Hing, C., Vlack, E.R.V., et al. (2023). Illuminating protein space with a programmable generative model. Nature 623, 1070–1078. 10.1038/s41586-023-06728-8.

63. Chao, G., Lau, W.L., Hackel, B.J., Sazinsky, S.L., Lippow, S.M., and Wittrup, K.D. (2006). Isolating and engineering human antibodies using yeast surface display. Nat. Protoc. 1, 755– 768. 10.1038/nprot.2006.94.

64. Punjani, A., Rubinstein, J.L., Fleet, D.J., and Brubaker, M.A. (2017). cryoSPARC: algorithms for rapid unsupervised cryo-EM structure determination. Nat. Methods 14, 290–296. 10.1038/nmeth.4169.

65. Goddard, T.D., Huang, C.C., Meng, E.C., Pettersen, E.F., Couch, G.S., Morris, J.H., and Ferrin, T.E. (2018). UCSF ChimeraX: Meeting modern challenges in visualization and analysis. Protein Sci. 27, 14–25. 10.1002/pro.3235.

66. Igaev, M., Kutzner, C., Bock, L.V., Vaiana, A.C., and Grubmüller, H. (2019). Automated cryo-EM structure refinement using correlation-driven molecular dynamics. eLife 8, e43542. 10.7554/elife.43542.

67. Abraham, M.J., Murtola, T., Schulz, R., Páll, S., Smith, J.C., Hess, B., and Lindahl, E. (2015). GROMACS: High performance molecular simulations through multi-level parallelism from laptops to supercomputers. SoftwareX 1, 19–25. 10.1016/j.softx.2015.06.001.

68. Lindorff-Larsen, K., Piana, S., Palmo, K., Maragakis, P., Klepeis, J.L., Dror, R.O., and Shaw, D.E. (2010). Improved side-chain torsion potentials for the Amber ff99SB protein force field. Proteins 78, 1950–1958. 10.1002/prot.22711.

69. Jorgensen, W.L., Chandrasekhar, J., Madura, J.D., Impey, R.W., and Klein, M.L. (1983). Comparison of simple potential functions for simulating liquid water. J. Chem. Phys. 79, 926– 935. 10.1063/1.445869.

70. Hess, B. (2007). P-LINCS: A Parallel Linear Constraint Solver for Molecular Simulation. J. Chem. Theory Comput. 4, 116–122. 10.1021/ct700200b.

71. Feenstra, K.A., Hess, B., and Berendsen, H.J.C. (1999). Improving efficiency of large time-scale molecular dynamics simulations of hydrogen-rich systems. J. Comput. Chem. 20, 786– 798. 10.1002/(sici)1096-987x(199906)20:8<786::aid-jcc5>3.0.co;2-b.

72. Bussi, G., Donadio, D., and Parrinello, M. (2007). Canonical sampling through velocity rescaling. J. Chem. Phys. 126, 014101. 10.1063/1.2408420.

73. Zimmerman, M.I., and Bowman, G.R. (2015). FAST Conformational Searches by Balancing Exploration/Exploitation Trade-Offs. J. Chem. Theory Comput. 11, 5747–5757. 10.1021/acs.jctc.5b00737.

74. Porter, J.R., Zimmerman, M.I., and Bowman, G.R. (2019). Enspara : Modeling molecular ensembles with scalable data structures and parallel computing. J. Chem. Phys. 150, 044108. 10.1063/1.5063794.

75. Subramanian, S., Schnell, G., Iulio, J. di, Gupta, A.K., Shapiro, A.E., Sarkis, E.H., Lopuski, A., Peppercorn, A., Aldinger, M., Hebner, C.M., et al. (2023). Resistance analysis following sotrovimab treatment in participants with COVID-19 during the phase III COMET-ICE study. Futur. Virol. 18, 975–990. 10.2217/fvl-2023-0146.

76. King, C.H.S., Keeney, J., Guimera, N., Das, S., Weber, M., Fochtman, B., Walderhaug, M.O., Talwar, S., Patel, J.A., Mazumder, R., et al. (2022). Communicating regulatory high-throughput sequencing data using BioCompute Objects. Drug Discov. Today 27, 1108–1114. 10.1016/j.drudis.2022.01.007.

77. Li, H. (2013). Aligning sequence reads, clone sequences and assembly contigs with BWA-MEM. arXiv. 10.48550/arxiv.1303.3997.

78. Garrison, E., and Marth, G. (2012). Haplotype-based variant detection from short-read sequencing. arXiv. 10.48550/arxiv.1207.3907.

79. Chen, C., Nadeau, S., Yared, M., Voinov, P., Xie, N., Roemer, C., and Stadler, T. (2021). CoV-Spectrum: analysis of globally shared SARS-CoV-2 data to identify and characterize new variants. Bioinformatics 38, 1735–1737. 10.1093/bioinformatics/btab856.

80. Chen, C., Taepper, A., Engelniederhammer, F., Kellerer, J., Roemer, C., and Stadler, T. (2023). LAPIS is a fast web API for massive open virus sequencing data. BMC Bioinform. 24, 232. 10.1186/s12859-023-05364-3.

