## Supplementary material for "Functional and structural characterization of a combination of pan-sarbecovirus antibodies with potent antiviral activity": Document S1

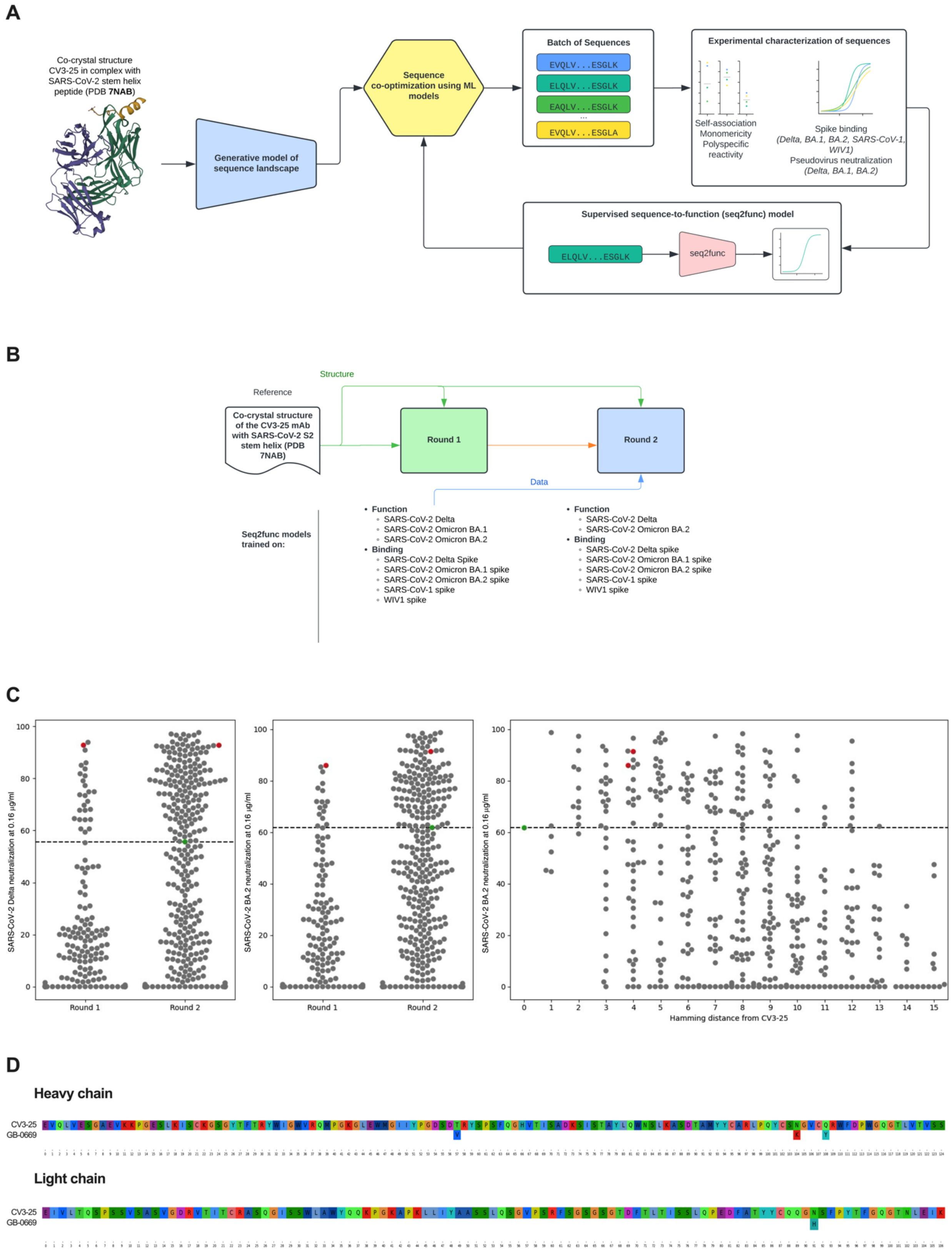


**Figure S1. Screening campaign and identification of anti-S2 stem helix antibodies with improved neutralizing activity.**
(A) Schematic representation of the optimization cycle for the anti-S2 stem helix campaign, outlining experimental characterization of novel antibody sequences and the two ML models used as part of this iterative process: generative model of sequence landscape that uses the co-crystal structure of the reference Fab CV3-25 with SARS-CoV-2 S2 stem helix peptide (PDB:

7NAB); sequence-to-function (seq2func) model built with data generated as part of the characterization of novel antibody sequences.

(B) Schematic representation of the screening campaign, including: structure leveraged for the generative model; the two rounds of sequence generation (round 1 and round 2); functional (pseudovirus neutralization) and binding (DELFIA with recombinant spike trimers) data generated for each round, with data from round 1 leveraged to build the seq2func model.

(C) Schematic representation of SARS-CoV-2 Delta and Omicron BA.2 pseudovirus neutralization data for round 1 and round 2. Graphs report percentage neutralization of the indicated pseudoviruses at the fixed mAb concentration of 0.16 μg/ml. Each dot represents an individual mAb, with the following color code: gray, designed mAbs; green, reference mAb CV3-25; red, lead candidate GB-0669. Hamming distance indicates number of positions at which each designed variant differs compared to the reference mAb CV3-25.

(D) Comparison of CV-35 and GB-0669 sequences representative of heavy and light chain variable regions.

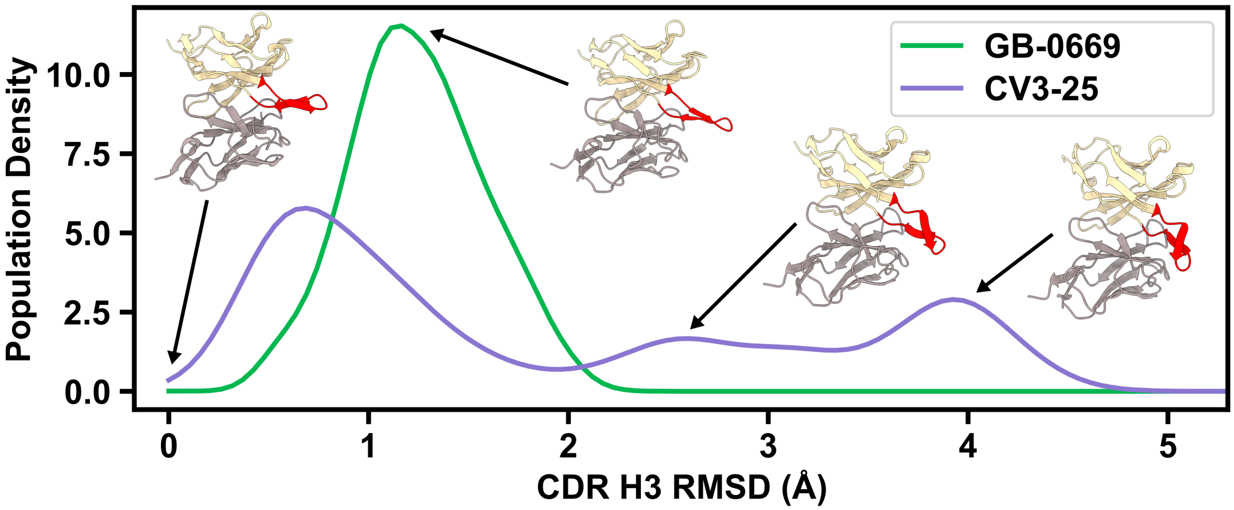


**Figure S2. Probability distributions from MD simulations of GB-0669 and CV3-25 show large differences in CDR H3 ensembles.**
Probability distributions for GB-0669 (green) and CV3-25 (purple) as a function of CDR H3 RMSD to the bound conformation. Representative structures are depicted as cartoon, with heavy chain colored tan, light chain colored gray, and CDR H3 colored red.

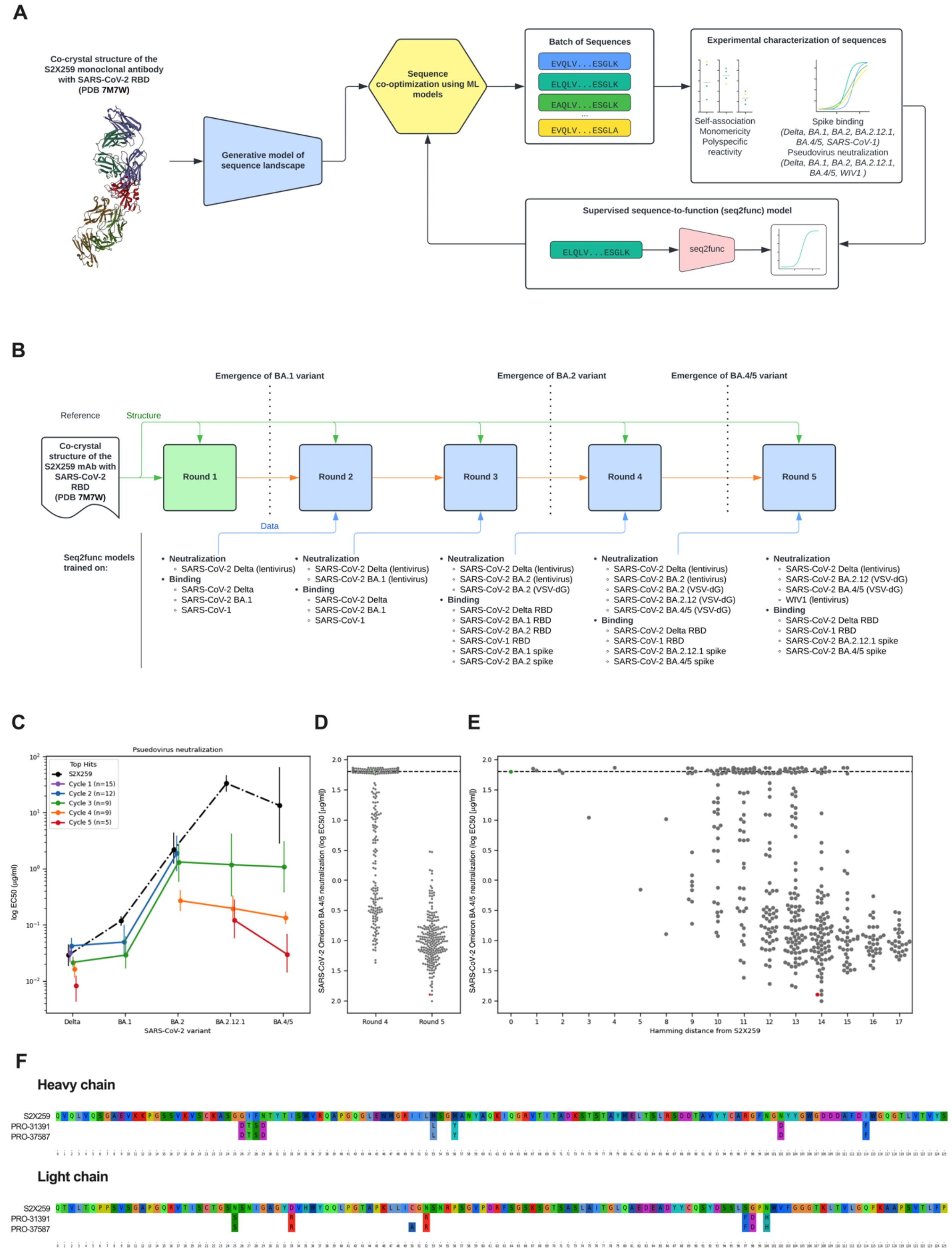


**Figure S3. Screening campaign and identification of class 4 anti-RBD antibodies with improved neutralizing activity.**
(A) Schematic representation of the optimization cycle for the class 4 anti-RBD campaign, outlining experimental characterization of novel antibody sequences and the two ML models used as part of this iterative process: generative model of sequence landscape that uses the co-crystal structure of the reference Fab S2X259 with SARS-CoV-2 RBD (PDB: 7M7W); sequence-

to-function (seq2func) model built with data generated as part of the characterization of novel antibody sequences.
(B) Schematic representation of the screening campaign, including: structure leveraged for the generative model; the five rounds of sequence generation (round 1 through 5); functional (pseudovirus neutralization) and binding (DELFIA with recombinant RBD proteins or spike trimers) data generated for each round, with data from round 1 through 4 leveraged to build the seq2func model.
(C) Representation of the neutralization potencies (reported as log EC50 [µg/ml]) for the indicated SARS-CoV-2 pseudoviruses of S2X259 and top hits across the 5 screening cycles.
(D) Schematic representation of SARS-CoV-2 Omicron BA.4/5 pseudovirus neutralization data (reported as log EC50 [µg/ml]) for round 4 and round 5. Each dot represents an individual mAb, with the following color code: gray, designed mAbs; green, reference mAb S2X259; red, candidate PRO-31391.
(E) Schematic representation of SARS-CoV-2 Omicron BA.4/5 pseudovirus neutralization data (reported as log EC50 [µg/ml]) as function of Hamming distance from S2X259 (number of positions at which each designed variant differs compared to the reference mAb S2X259). Each dot represents an individual mAb, with the following color code: gray, designed mAbs; green, reference mAb S2X259; red, candidate PRO-31391.
(F) Comparison of S2X259, PRO-31391 and PRO-37587 sequences representative of heavy and light chain variable regions.

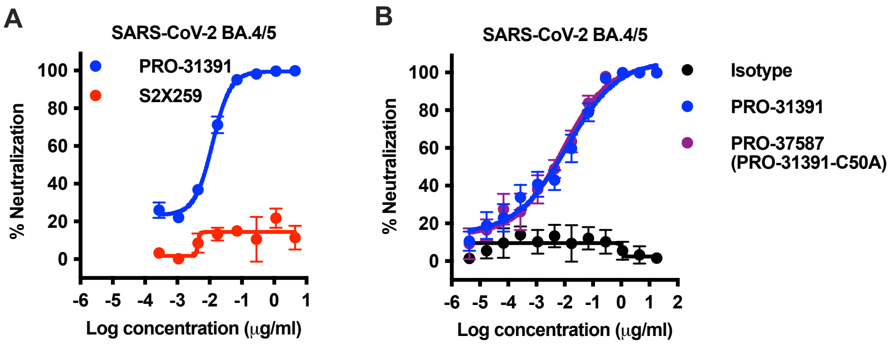


**Figure S4. Optimization of the class 4 anti-RBD hit PRO-31391 and identification of the lead molecule PRO-37587.**

(A) Neutralization profiles of PRO-31391 and the reference mAb against SARS-CoV-2 Omicron BA.4/5 pseudovirus. Results are reported as percentage (%) neutralization and shown as mean ± SD (representative of one experiment, two [PRO-31391] or four [S2X259] technical replicates).

(B) Neutralization profiles of PRO-31391, PRO-37587 (same sequence as PRO-31391 with C50A substitution [PRO-31391-C50A]) and isotype control mAb against SARS-CoV-2 Omicron BA.4/5 pseudovirus. Results are reported as percentage (%) neutralization and shown as mean ± SD (representative of two independent experiments, four technical replicates each).

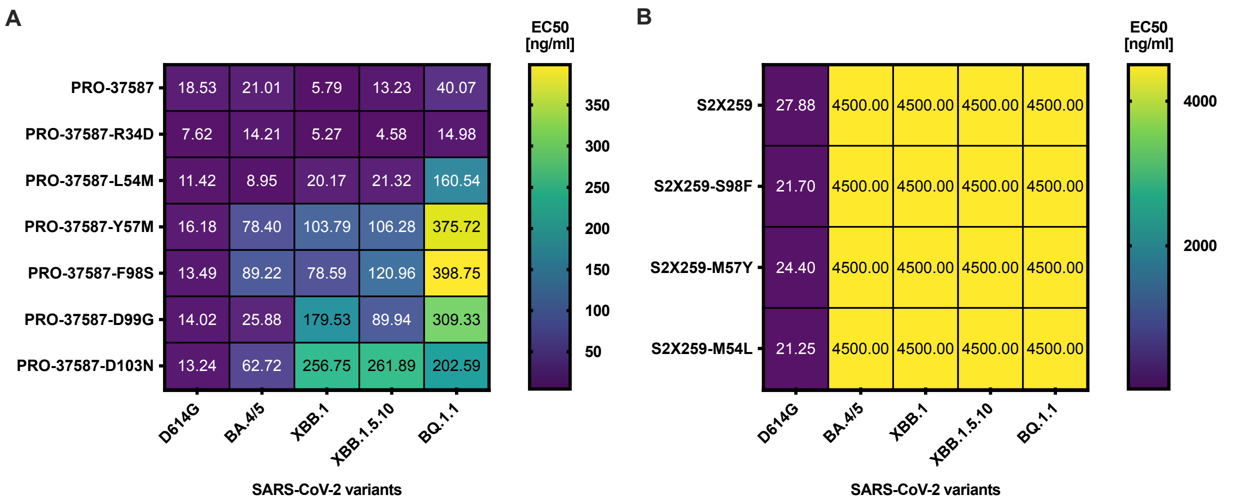


**Figure S5. Contribution of selected CDR mutations in PRO-37587 to breadth of SARS-CoV-2 pseudovirus neutralization.**
(A) Heatmap representing the neutralization profiles of PRO-37587 and variants with selected CDR amino acids reverted to those of the reference mAb S2X259, tested against the indicated SARS-CoV-2 variants.
(B) Heatmap representing the neutralization profiles of reference mAb S2X259 and variants with selected CDR amino acids mutated to those of the lead molecule PRO-37587, tested against the indicated SARS-CoV-2 variants.
Numbers in the cells represent the corresponding EC50 values (average of two technical replicates and expressed in ng/ml) and are color-coded according to the provided color scales. EC50 values greater than 4,500 ng/ml (top concentration tested in the neutralization curve) are reported as 4,500.

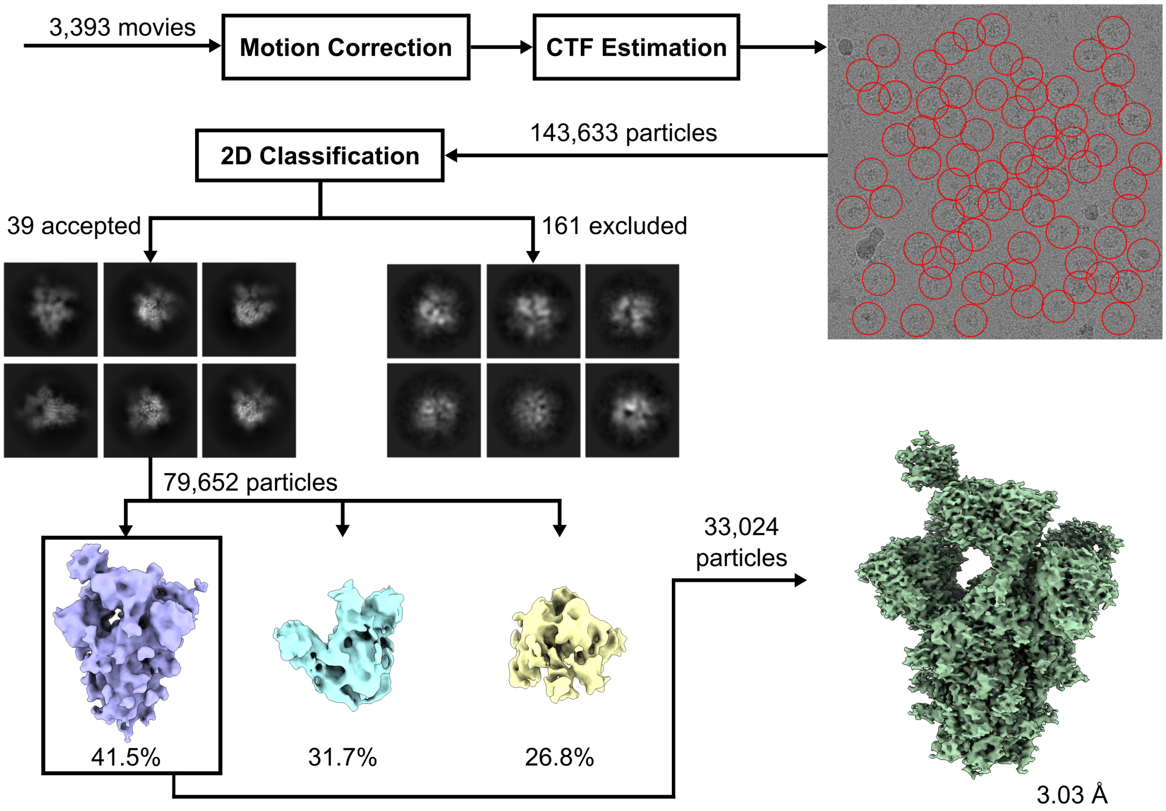


**Figure S6. Cryo-EM data processing workflow of SARS-CoV-2 Omicron BA.1 spike bound to PRO-37587 (part 1).**
Illustrated data stream for the first set of movies collected of the spike:antibody complex. Processing consisted of motion correction, CTF estimation, particle picking, 2D classification, *ab initio* reconstruction, and a non-uniform refinement. The final map had a good nominal resolution, however, was not sufficient for model building as judged by visual inspection. The final particles were pooled with the second data collection processing stream, as illustrated in Figure S7.

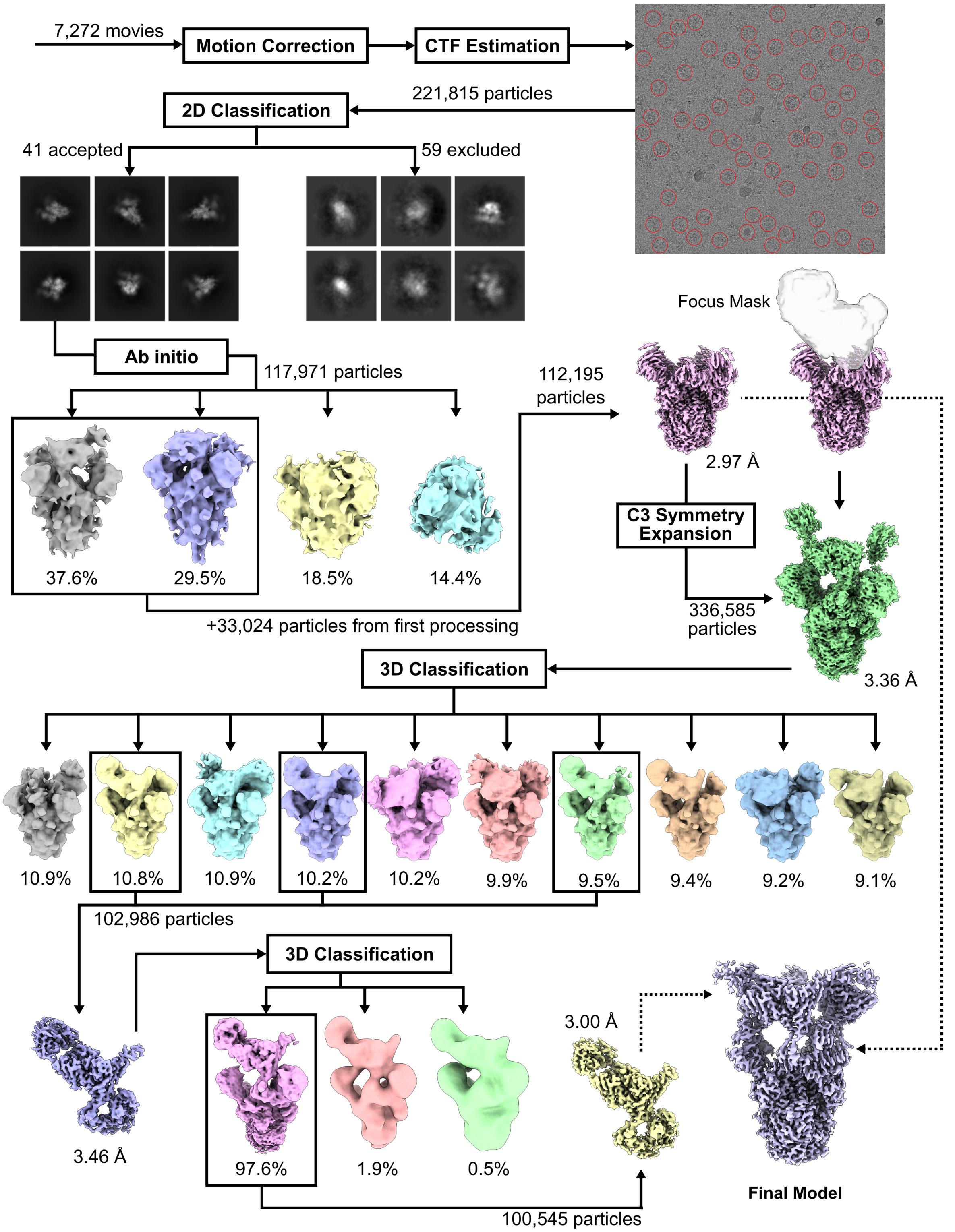


**Figure S7. Cryo-EM data processing workflow of SARS-CoV-2 Omicron BA.1 spike bound to PRO-37587 (part 2).**
Illustrated data stream for the second set of movies collected of the spike:antibody complex. Processing consisted of motion correction, CTF estimation, particle picking, 2D classification, ab initio reconstruction, and were then pooled with particles from Figure S6 for a non-uniform refinement. Symmetry expansion followed by a series of focused refinements and 3D classifications around the SARS-CoV-2 Omicron BA.1 RBD and PRO-37587 led to a 3.00 Å map for this region that was sufficient for an atomically detailed model. A final composite map was generated by merging the consensus map with the locally refined map around the RBD:PRO-37587 binding region.

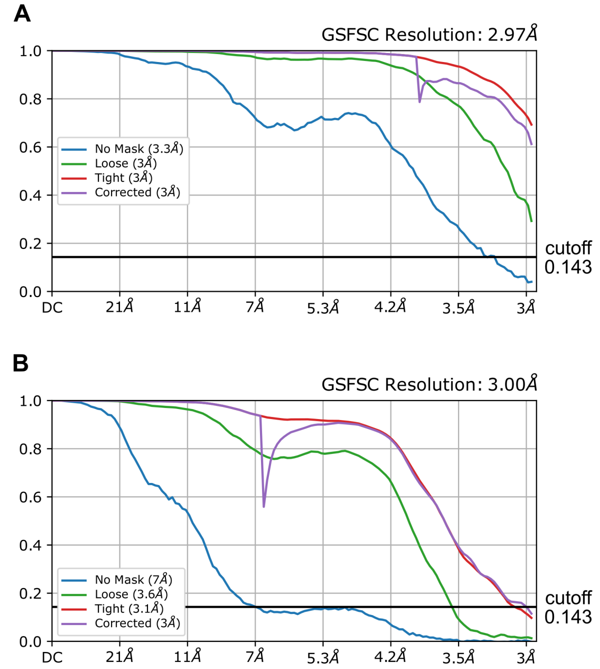


**Figure S8. Gold-Standard Fourier Shell Correlation (GSFSC) for consensus map and local refinement.**
Curves represent Fourier shell correlations between independently aligned half maps, where a cutoff of 0.143 was used to determine final resolutions. Shown are the GSFSC curves for (A) the consensus map (2.97 Å) and (B) the locally refined region around RBD and PRO-37587 (3.00 Å).

**Table S1. Neutralization profiles of GB-0669 and CV3-25 against pseudoviruses representative of SARS-CoV-2 variants and non-SARS-CoV-2 sarbecoviruses.**

| Strain | Neutralization Parameters[a] | GB-0669 [ng/ml] | CV3-25 [ng/ml] |
|---|---|---|---|
| **SARS-CoV-2 D614G** | EC50 (95% CI) | 19 (16 to 24) | 180 (110 to 380) |
| | EC90 (95% CI) | 600 (390 to 1,000) | 8,900 (2,400 to 73,000) |
| | Emax % (95% CI) | 94.36 (93.79 to 94.92) | 84.69 (83.42 to 85.96) |
| **SARS-CoV-2 Delta** | EC50 (95% CI) | 39 (30 to 52) | 800 (360 to 4,600) |
| | EC90 (95% CI) | 1,400 (730 to 3,100) | 130,000 (20,000 to 4,980,000) |
| | Emax % (95% CI) | 87.73 (86.31 to 89.16) | 73.30 (71.73 to 75.30) |
| **SARS-CoV-2 BA.2** | EC50 (95% CI) | 9.6 (6.9 to 13) | 160 (73 to 740) |
| | EC90 (95% CI) | 460 (220 to 1,100) | 82,000 (11,000 to 4,400,000) |
| | Emax % (95% CI) | 90.52 (89.16 to 91.89) | 84.72 (82.60 to 86.83) |
| **SARS-CoV-2 BA.4/5** | EC50 (95% CI) | 4.8 (2.4 to 8.4) | NA[b] |
| | EC90 (95% CI) | 580 (190 to 3,200) | NA[b] |
| | Emax % (95% CI) | 89.75 (88.37 to 91.12) | 84.67 (82.51 to 86.83) |
| **SARS-CoV-2 BQ.1.1** | EC50 (95% CI) | 5.1 (1.9 to 10) | NA[b] |
| | EC90 (95% CI) | 1,000 (240 to 14,000) | NA[b] |
| | Emax % (95% CI) | 93.54 (92.15 to 94.93) | 87.48 (85.19 to 89.78) |
| **SARS-CoV-2 XBB.1.5** | EC50 (95% CI) | 8.8 (6.5 to 12) | 93 (33 to 3,000) |
| | EC90 (95% CI) | 540 (280 to 1200) | NA[b] |
| | Emax % (95% CI) | 92.88 (91.87 to 93.89) | 88.11 (86.67 to 89.54) |
| **SARS-CoV-1** | EC50 (95% CI) | 12 (9.4 to 15) | 220 (130 to 540) |
| | EC90 (95% CI) | 240 (150 to 410) | 31,000 (7,800 to 270,000) |
| | Emax % (95% CI) | 95.81 (94.85 to 96.77) | 86.11 (84.05 to 88.17) |
| **WIV1-CoV** | EC50 (95% CI) | 63 (53 to 75) | 1,100 (640 to 2,700) |
| | EC90 (95% CI) | 1,500 (970 to 2,300) | 89,000 (26,000 to 550,000) |
| | Emax % (95% CI) | 92.90 (91.59 to 94.20) | 81.97 (79.69 to 84.24) |

[a]EC50 and EC90 expressed in ng/ml; Emax % represents percentage neutralization at the highest antibody concentration (18,000 ng/ml).
[b]Poor or incomplete best fit value, unable to determine confidence interval.

**Table S2. Neutralization profiles of PRO-37587 and S2X259 against pseudoviruses representative of SARS-CoV-2 variants and non-SARS-CoV-2 sarbecoviruses.**

| Strain | Neutralization Parameters[a] | PRO-37587 [ng/ml] | S2X259 [ng/ml] |
|---|---|---|---|
| **SARS-CoV-2 D614G** | EC50 (95% CI) | 22 (18 to 25) | 21 (17 to 26) |
| | EC90 (95% CI) | 220 (150 to 320) | 430 (280 to 740) |
| **SARS-CoV-2 Delta** | EC50 (95% CI) | 12 (9 to 16) | 25 (19 to 35) |
| | EC90 (95% CI) | 210 (120 to 420) | 630 (320 to 1500) |
| **SARS-CoV-2 BA.2** | EC50 (95% CI) | 27 (22 to 32) | NA[b] |
| | EC90 (95% CI) | 310 (210 to 500) | NA[b] |
| **SARS-CoV-2 BA.4/5** | EC50 (95% CI) | 15 (11 to 22) | NA[b] |
| | EC90 (95% CI) | 150 (76 to 370) | NA[b] |
| **SARS-CoV-2 BQ.1.1** | EC50 (95% CI) | 38 (31 to 48) | NA[b] |
| | EC90 (95% CI) | 320 (200 to 550) | NA[b] |
| **SARS-CoV-2 XBB.1.5** | EC50 (95% CI) | 12 (9 to16) | NA[b] |
| | EC90 (95% CI) | 140 (81 to 250) | NA[b] |
| **SARS-CoV-1** | EC50 (95% CI) | 0.77 (0.52 to 1.1) | 2.2 (1.4 to 3.3) |
| | EC90 (95% CI) | 7.4 (3.7 to 16) | 50 (23 to 120) |
| **WIV1-CoV** | EC50 (95% CI) | 0.38 (0.19 to 0.65) | 1.7 (0.95 to 2.7) |
| | EC90 (95% CI) | 14 (6.5 to 33) | 140 (63 to 370) |

[a]EC50 and EC90 expressed in ng/ml.
[b]Poor or incomplete best fit value, unable to determine confidence interval.

**Table S3. Amino acid sequences of RBDs representative of SARS-CoV-2-related clade 1b, SARS-CoV-related clade 1a, Asia non-ACE2 bat sarbecovirus (sarbecov.) clade 2, and Africa+Europe bat sarbecovirus clade 3 expressed on yeast.**

| Receptor Binding Domains | Sequence |
|---|---|
| P0DTC2_Omicron\|316-538 Domain (Ancestral/SARS-CoV-2) | RVQPTESIVRFPNITNLCPFDEVFNATRFASVYAWNRKRISNCVADYSVLYNLAP<br>FFTFKCYGVSPTKLNDLCFTNVYADSFVIRGDEVRQIAPGQTGNIADYNYKLPDD<br>FTGCVIAWNSNKLDSKVSGNYNYLYRLFRKSNLKPFERDISTEIYQAGNKPCNG<br>VAGFNCYFPLRSYSFRPTYGVGHQPYRVVVLSFELLHAPATVCGPKKSTNLVKN<br>KCVNF |
| SARS-CoV-2 B.1.617.2 Delta 319-541 (L452R, T478K) (Delta) | RVQPTESIVRFPNITNLCPFGEVFNATRFASVYAWNRKRISNCVADYSVLYNSAS<br>FSTFKCYGVSPTKLNDLCFTNVYADSFVIRGDEVRQIAPGQTGKIADYNYKLPDD<br>FTGCVIAWNSNNLDSKVGGNYNYRYRLFRKSNLKPFERDISTEIYQAGSKPCNG<br>VEGFNCYFPLQSYGFQPTNGVGYQPYRVVVLSFELLHAPATVCGPKKSTNLVK<br>NKCVNF |
| SARS-CoV-1 306-527 (SARS-CoV-1) | RVVPSGDVVRFPNITNLCPFGEVFNATKFPSVYAWERKKISNCVADYSVLYNST<br>FFSTFKCYGVSATKLNDLCFSNVYADSFVVKGDDVRQIAPGQTGVIADYNYKLP<br>DDFMGCVLAWNTRNIDATSTGNYNYKYRYLRHGKLRPFERDISNVPFSPDGKP<br>CTPPALNCYWPLNDYGFYTTTGIGYQPYRVVVLSFELLNAPATVCGPKLSTDLIK<br>NQCVNF |
| SARS-CoV-2 omicron BA.1 319-541 \|EPI_ISL_9976794 (BA.1) | RVQPTESIVRFPNITNLCPFDEVFNATRFASVYAWNRKRISNCVADYSVLYNLAP<br>FFTFKCYGVSPTKLNDLCFTNVYADSFVIRGDEVRQIAPGQTGKIADYNYKLPDD<br>FTGCVIAWNSNNLDSKVGGNYNYLYRLFRKSNLKPFERDISTEIYQAGNKPCNG<br>VAGFNCYFPLRSYSFRPTYGVGHQPYRVVVLSFELLHAPATVCGPKKSTNLVKN<br>KCVNF |
| SARS-CoV-2 omicron BA.1.1 319-541 \|EPI_ISL_9936303 (BA.1.1) | RVQPTESIVRFPNITNLCPFDEVFNATKFASVYAWNRKRISNCVADYSVLYNLAP<br>FFTFKCYGVSPTKLNDLCFTNVYADSFVIRGDEVRQIAPGQTGKIADYNYKLPDD<br>FTGCVIAWNSNNLDSKVGGNYNYLYRLFRKSNLKPFERDISTEIYQAGNKPCNG<br>VAGFNCYFPLRSYSFRPTYGVGHQPYRVVVLSFELLHAPATVCGPKKSTNLVKN<br>KCVNF |
| SARS-CoV-2 omicron BA.2 319-541 \|EPI_ISL_9984188 (BA.2) | RVQPTESIVRFPNITNLCPFDEVFNATRFASVYAWNRKRISNCVADYSVLYNFAP<br>FFAFKCYGVSPTKLNDLCFTNVYADSFVIRGNEVSQIAPGQTGNIADYNYKLPDD<br>FTGCVIAWNSNKLDSKVGGNYNYLYRLFRKSNLKPFERDISTEIYQAGNKPCNG<br>VAGFNCYFPLRSYGFRPTYGVGHQPYRVVVLSFELLHAPATVCGPKKSTNLVK<br>NKCVNF |
| GD-Pangolin (GD-Pangolin) | NITNLCPFGEVFNATTFASVYAWNRKRISNCVADYSVLYNSTSFSTFKCYGVSPT<br>KLNDLCFTNVYADSFVVRGDEVRQIAPGQTGRIADYNYKLPDDFTGCVIAWNSN<br>NLDSKVGGNYNYLYRLFRKSNLKPFERDISTEIYQAGSTPCNGVEGFNCYFPLQ<br>SYGFHPTNGVGYQPYRVVVLSFELLNAPATVCGPKQST |
| RaTG13 (RaTG13) | NITNLCPFGEVFNATTFASVYAWNRKRISNCVADYSVLYNSTSFSTFKCYGVSPT<br>KLNDLCFTNVYADSFVITGDEVRQIAPGQTGKIADYNYKLPDDFTGCVIAWNSKH<br>IDAKEGGNFNYLYRLFRKANLKPFERDISTEIYQAGSKPCNGQTGLNCYYPLYRY<br>GFYPTDGVGHQPYRVVVLSFELLNAPATVCGPKKST |
| GX-Pangolin (GX-Pangolin) | NITNLCPFGEVFNASKFASVYAWNRKRISNCVADYSVLYNSTSFSTFKCYGVSP<br>TKLNDLCFTNVYADSFVVKGDEVRQIAPGQTGVIADYNYKLPDDFTGCVIAWNS<br>VKQDALTGGNYGYLYRLFRKSKLKPFERDISTEIYQAGSTPCNGQVGLNCYYPL<br>ERYGFHPTTGVNYQPFRVVVLSFELLNGPATVCGPKLST |
| SARS-CoV-1_HGZ8L1-A_HP03E (HGA8L1-A) | NITNLCPFGEVFNATKFPSVYAWERKRISNCVADYSVLYNSTFFSTFKCYGVSAT<br>KLNDLCFSNVYADSFVVKGDDVRQIAPGQTGVIADYNYKLPDDFMGCVLAWNT<br>RNIDATSTGNYNYKYRYLRHGKLRPFERDISNVPFSPDGKPCTPPALNCYWPLN<br>DYGFYTTTGIGYQPYRVVVLSFELLNAPATVCGPKLST |
| SARS-CoV-1_GZ-C_HP03L (GZC) | NITNLCPFGEVFNATKFPSVYAWERKKISNCVADYSVLYNSTFFSTFKCYGVSAT<br>KLNDLCFSNVYADSFVVKGDDVRQIAPGQTGVIADYNYKLPDDFMGCVLAWNT<br>RNIDATSTGNHNYKYRYLRHGKLRPFERDISNVPFSPDGKPCTPPALNCYWPLN<br>DYGFYTTTGIGYQPYRVVVLSFELLNAPATVCGPKLST |
| SARS-CoV-1_Sin852_HP03L (Sin852) | NITNLCPFGEVFNATKFPSVYAWERKKISNCVADYFVLYNSTFFSTFKCYGVSAT<br>KLNDLCFSNVYADSFVVKGDDVRQIAPGQTGVIADYNYKLPDDFMGCVLAWNT<br>RNIDATSTGNYNYKYRYLRHGKLRPFERDISNVPFSPDGKPCTPPALNCYWPLN<br>DYGFYTTTGIGYQPYRVVVLSFELLNAPATVCGPKLST |

| | |
|---|---|
| SARS-CoV-1_GD03T0013_HP04<br>(GD03 T0013) | NITNLCPFGEVFNATKFPSVYAWERKRISNCVADYSVLYNSTSFSTFKCYGVSATKLNDLCFSNVYADSFVVKGDDVRQIAPGQTGVIADYNYKLPDDFMGCVLAWNTRNIDATSTGNYNYKYRYLRHGKLRPFERDISNVPFSPDGKPCTPPAPNCYWPLNGYGFYTTSGIGYQPYRVVVLSFELLNAPATVCGPKLST |
| SARS-CoV-1_GZ0402_HP04<br>(GZ0402) | NITNLCPFGEVFNATKFPSVYAWERKRISNCVADYSVLYNSTSFSTFKCYGVSATKLNDLCFSNVYADSFVVKGDDVRQIAPGQTGVIADYNYKLPDDFMGCVLAWNIRNIDATSTGNYNYKYRYLRHGKLRPFERDISNVPFSPDGKPCTPPAPNCYWPLNGYGFYTTSGIGYQPYRVVVLSFELLNAPATVCGPKLST |
| SARS-CoV-1_PC4-127_PC04<br>(PC4-127) | NITNLCPFGEVFNATKFPSVYAWERKRISNCVADYSVLYNSTSFSTFKCYGVSATKLNDLCFSNVYADSFVVKGDDVRQIAPGQTGVIADYNYKLPDDFMGCVLAWNTRNIDATSTGNYNYKYRYLRHGKLRPFERDISNVPFSPDGKPCTPPAPNCYWPLRGYGFYTTSGIGYQPYRVVVLSFELLNAPATVCGPKLST |
| SARS-CoV-1_PC4-137_PC04<br>(PC4-137) | NITNLCPFGEVFNATKFPSVYAWERKRISNCVADYSVLYNSTSFSTFKCYGVSATKLNDLCFSNVYADSFVVKGDDVRQIAPGQTGVIADYNYKLPDDFMGCVLAWNTRNIDATSTGNYNYKYRYLRHGKLRPFERDISNVPFSPDGKPCTPPAPNCYWPLKGYGFYTTSGIGYQPYRVVVLSFELLNAPATVCGPKLST |
| SARS-CoV-1_PC4-13_PC04<br>(PC4-13) | NITNLCPFGEVFNATKFPPVYAWERKRISNCVADYSVLYNSTSFSTFKCYGVSATKLNDLCFSNVYADSFVVKGDDVRQIAPGQTGVIADYNYKLPDDFMGCVLAWNTRNIDATSTGNYNYKYRYLRHGKLRPFERDISNVPFSSDGKPCTPPAPNCYWPLRGYGFYTTSGIGYQPYRVVVLSFELLNAPATVCGPKLST |
| WIV1<br>(WIV1) | NITNLCPFGEVFNATTFPSVYAWERKRISNCVADYSVLYNSTSFSTFKCYGVSATKLNDLCFSNVYADSFVVKGDDVRQIAPGQTGVIADYNYKLPDDFTGCVLAWNTRNIDATQTGNYNYKYRSLRHGKLRPFERDISNVPFSPDGKPCTPPAFNCYWPLNDYGFYITNGIGYQPYRVVVLSFELLNAPATVCGPKLST |
| Rs7327<br>(Rs7327) | NITNLCPFGEVFNATTFPSVYAWERKRISNCVADYSVLYNSTSFSTFKCYGVSATKLNDLCFSNVYADSFVVKGDDVRQIAPGQTGVIADYNYKLPDDFMGCVLAWNTRNIDATSTGNYNYKYRSLRHGKLRPFERDISNVPFSPDGKPCTPPAFNCYWPLNDYGFFTTNGIGYQPYRVVVLSFELLNAPATVCGPKLST |
| Rs4231<br>(Rs4231) | NITNLCPFGEVFNATTFPSVYAWERKRISNCVADYSVLYNSTSFSTFKCYGVSATKLNDLCFSNVYADSFVVKGDDVRQIAPGQTGVIADYNYKLPDDFLGCVLAWNTNSKDSSTSGNYNYLYRWVRRSKLNPYERDLSNDIYSPGGQSCSAIGPNCYNPLRPYGFFTTAGVGHQPYRVVVLSFELLNAPATVCGPKLST |
| Rs4084<br>(Rs4084) | NITNLCPFGEVFNATTFPSVYAWERKRISNCVADYSILYNSTSFSTFKCYGVSATKLNDLCFSNVYADSFVVKGDDVRQIAPGQTGVIADYNYKLPDDFLGCVLAWNTNSKDSSTSGNYNYLYRWVRRSKLNPYERDLSNDIYSPGGQSCSAVGPNCYNPLRPYGFFTTAGVGHQPYRVVVLSFELLNAPATVCGPKLST |
| BM48-31<br>(BM48-31) | NITQLCPFNEVFNITSFPSVYAWERMRITNCVADYSVLYNSSASFSTFQCYGVSPTKLNDLCFSSVYADYFVVKGDDVRQIAPAQTGVIADYNYKLPDDFTGCVIAWNTNSLDSSNEFFYRRFRHGKIKPYGRDLSNVLFNPSGGTCSAEGLNCYKPLASYGFTQSSGIGFQPYRVVVLSFELLNAPATVCGPKQST |
| BtKY72<br>(BtKY72) | NITNLCPFGQVFNASNFPSVYAWERLRISDCVADYAVLYNSSSSFSTFKCYGVSPTKLNDLCFSSVYADYFVVKGDDVRQIAPAQTGVIADYNYKLPDDFTGCVLAWNTNSVDSKSGNNFYYRLFRHGKIKPYERDISNVLYNSAGGTCSSISQLGCYEPLKSYGFTPTVGVGYQPYRVVVLSFELLNAPATVCGPKKST |
| ZXC21<br>(ZXC21) | NITNVCPFHKVFNATRFPSVYAWERTKISDCIADYTVFYNSTSFSTFKCYGVSPSKLIDLCFTSVYADTFLIRFSEVRQVAPGQTGVIADYNYKLPDDFTGCVIAWNTAKQDTGHYFYRSHRSTKLKPFERDLSSDENGVRTLSTYDFNPNVPLEYQATRVVVLSFELLNAPATVCGPKLST |
| ZC45<br>(ZC45) | NITNVCPFHKVFNATRFPSVYAWERTKISDCIADYTVFYNSTSFSTFKCYGVSPSKLIDLCFTSVYADTFLIRFSEVRQVAPGQTGVIADYNYKLPDDFTGCVIAWNTAKQDVGNYFYRSHRSTKLKPFERDLSSDENGVRTLSTYDFNPNVPLEYQATRVVVLSFELLNAPATVCGPKLST |
| JL2012<br>(JL2012) | NITNVCPFDKVFNATRFPSVYAWERTKISDCVADYTVFYNSTSFSTFNCYGVSPSKLIDLCFTSVYADTFLIRFSEVRQVAPGQTGVIADYNYKLPDDFIGCVIAWNTAKQDVGSYFYRSHRSSKLKPFERDLSSEENGVLTLSTYDFNQNVPLEYQATRVVVLSFELLNAPATVCGPKLST |
| Rf1<br>(Rf1) | NITNLCPFDKVFNATRFPSVYAWERTKISDCVADYTVFYNSTSFSTFNCYGVSPSKLIDLCFTSVYADTFLIRFSEVRQVAPGQTGVIADYNYKLPDDFTGCVIAWNTAKQDVGSYFYRSHRSSKLKPFERDLSSEENGVRTLSTYDFNQNVPLEYQATRVVVLSFELLNAPATVCGPKLST |

| | |
|---|---|
| HeB2013<br>(HeB2013) | NITNLCPFDKVFNATRFPSVYAWERTKISDCVADYTVFYNSTSFSTFNCYGVSPS<br>KLIDLCFTSVYADTFLIRFSEVRQVAPGQTGVIADYNYKLPDDFTGCVIAWNTAK<br>QDVGSYFYRSHRSSKLKPFERDLSSEENGVRTLSTYDFNQYVPLEYQATRVVV<br>LSFELLNAPATVCGPKLST |
| 273-2005<br>(273-2005) | NITNLCPFDKVFNATRFPSVYAWERTKISDCVADYTVFYNSTSFSTFNCYGVSPS<br>KLIDLCFTSVYADTFLIRFSEVRQVAPGQTGVIADYNYKLPDDFTGCVIAWNTAK<br>QDVGSYFYRSHRSSKLKPFERDLSSVEENGRTLSTYDFNQNVPLEYQATRVVV<br>LSFELLNAPATVCGPKLST |
| YN2013<br>(YN2013) | NITNRCPFDSIFNASRFPSVYAWERTKISDCVADYTVLYNSTLFSTFKCYGVSPS<br>KLIDLCFTSVYADTFLIRFSEVRQVAPGETGVIADYNYRLPDDFTGCVIAWNTAN<br>QDVGSYFYRSHRSTKLKPFERDLSSDENGVRTLSTYDFNPNVPLDYQATRVVV<br>LSFELLNAPATVCGPKLST |
| RmYN02<br>(RmYN02) | NITNFCPFDKVFNATRFPNVYAWQRTKISDCIADYTVLYNSTSFSTFKCYGVSPS<br>KLIDLCFTSVYADTFLIRFSEVRQIAPGETGVIADYNYKLPDDFTGCVLAWNTAQ<br>QDIGSYFYRSHRAVKLKPFERDLSSDENGVRTLSTYDFNPNVPLDYQATRVVVL<br>SFELLNAPATVCGPKLST |
| As6526<br>(As6526) | NITNRCPFDKVFNATRFPSVYAWERTKISDCVADYTVLYNSTSFSTFKCYGVSP<br>SKLIDLCFTSVYADTFLIRSSEVRQVAPGETGVIADYNYKLPDDFTGCVIAWNTA<br>QQDKGQYYYRSSRKTKLKPFERDLSSDENGVRTLSTYDFYPTVPIEYQATRVVV<br>LSFELLNAPATVCGPKLST |
| Rs4237<br>(Rs4237) | NITNRCPFDKVFNASRFPNVYAWERTKISDCVADYTVLYNSTSFSTFKCYGVSP<br>SKLIDLCFTSVYADTFLIRSSEVRQVAPGETGVIADYNYKLPDDFTGCVIAWNTAK<br>QDQGQYYYRSSRKTKLKPFERDLSSDENGVRTLSTYDFYPTVPIEYQATRVVVL<br>SFELLNAPATVCGPKLST |
| Rs4081<br>(Rs4081) | NITNRCPFDKVFNASRFPNVYAWERTKISDCVADYTVLYNSTSFSTFKCYGVSP<br>SKLIDLCFTSVYADTFLIRSSEVRQVAPGETGVIADYNYKLPDDFTGCVIAWNTAK<br>QDQGQYYYRSSRKTKLKPFERDLTSDENGVRTLSTYDFYPNVPIEYQATRVVVL<br>SFELLNAPATVCGPKLST |
| Rp3<br>(Rp3) | NITNRCPFDKVFNATRFPNVYAWERTKISDCVADYTVLYNSTSFSTFKCYGVSP<br>SKLIDLCFTSVYADTFLIRSSEVRQVAPGETGVIADYNYKLPDDFTGCVIAWNTAK<br>QDQGQYYYRSHRKTKLKPFERDLSSDENGVRTLSTYDFYPSVPVAYQATRVVV<br>LSFELLNAPATVCGPKLST |
| 279-2005<br>(279-2005) | NITNRCPFDKVFNASRFPNVYAWERTKISDCVADYTVLYNSTSFSTFKCYGVSP<br>SKLIDLCFTSVYADTFLIRSSEVRQVAPGETGVIADYNYKLPDDFTGCVIAWNTA<br>QQDQGQYYYRSYRKEKLKPFERDLSSDENGVYTLSTYDFYPSIPVEYQATRVV<br>VLSFELLNAPATVCGPKLST |
| Shaanxi2011<br>(Shaanxi2011) | NITNRCPFDKVFNATRFPSVYAWERTKISDCVADYTVLYNSTSFSTFKCYGVSP<br>SKLIDLCFTSVYADTFLIRSSEVRQVAPGETGVIADYNYKLPDDFTGCVIAWNTA<br>NQDQGQYYYRSSRKEKLKPFERDLSSDENGVYTLSTYDFYPSVPLDYQATRVV<br>VLSFELLNAPATVCGPKLST |
| Yunnan2011<br>(Yunnan2011) | NITNRCPFDRVFNASRFPSVYAWERTKISDCVADYTVLYNSTSFSTFKCYGVSP<br>SKLIDLCFTSVYADTFLIRFSEVRQIAPGETGVIADYNYKLPDEFTGCVIAWNTAN<br>QDRGQYYYRSSRKTKLKPFERDLSSDENGVRTLSTYDFYPSVPLEYQATRVVV<br>LSFELLNAPATVCGPKLST |
| Rs4247<br>(Rs4247) | NITNRCPFDKVFNASRFPNVYAWERTKISDCVADYTVLYNSTSFSTFKCYGVSP<br>SKLIDLCFTSVYADTFLIRSSEVRQVAPGETGVIADYNYKLPDDFTGCVIAWNTAK<br>QDTGHYYYRSHRKTKLKPFERDLSSDDGNGVYTLSTYDFNPNVPVAYQATRVV<br>VLSFELLNAPATVCGPKLST |
| HKU3-1<br>(HKU3-1) | NITNRCPFDKVFNATRFPNVYAWERTKISDCVADYTVLYNSTSFSTFKCYGVSP<br>SKLIDLCFTSVYADTFLIRSSEVRQVAPGETGVIADYNYKLPDDFTGCVIAWNTAK<br>HDTGNYYYRSHRKTKLKPFERDLSSDDGNGVYTLSTYDFNPNVPVAYQATRVV<br>VLSFELLNAPATVCGPKLST |
| GX2013<br>(GX2013) | NITNRCPFDKVFNATRFPNVYAWERTKISDCVADYTVLYNSTSFSTFKCYGVSP<br>SKLIDLCFTSVYADTFLIRSSEVRQVAPGETGVIADYNYKLPDDFTGCVIAWNTAK<br>QDTGNYYYRSHRKTKLKPFERDLSSDDGNGVYTLSTYDFNPNVPVAYQATRVV<br>VLSFELLNAPATVCGPKLST |
| Longquan-140<br>(Longquan-140) | NITNRCPFDKVFNVTRFPNVYAWERTKISDCVADYTVLYNSTSFSTFKCYGVSP<br>SKLIDLCFTSVYADTFLIRSSEVRQVAPGETGVIADYNYKLPDDFTGCVIAWNTAK<br>QDIGNYYYRSHRKTKLKPFERDLSSDDGNGVYTLSTYDFNPNVPVAYQATRVV<br>VLSFELLNAPATVCGPKLST |

| | |
|---|---|
| HKU3-8<br>(HKU3-8) | NITNRCPFDRVFNASRFPSVYAWERTKISECVADYTVLYNSTSFSTFKCYGVSP<br>SKLIDLCFTSVYADTFLIRSSEVRQVAPGETGVIADYNYKLPDDFTGCVIAWNTAK<br>QDTGNYYYRSHRKTKLKPFERDLSSDDGNGVYTLSTYDFNPNVPVAYQATRVV<br>VLSFELLNAPATVCGPKLST |
| HuB2013<br>(HuB2013) | NITNRCPFDRVFNASRFPSVYAWERTKISDCVADYTVLYNSTSFSTFKCYGVSP<br>SKLIDLCFTSVYADTFLIRSSEVRQVAPGETGVIADYNYKLPDDFTGCVIAWNTAK<br>QDTGYYYYRSHRKTKLKPFERDLSSDDGNGVYTLSTYDFNPNVPVAYQATRVV<br>VLSFELLNAPATVCGPKLST |
| LC556375.1<br>(LC556375.1) | NITNLCPFSEVFNATTFASVYAWNRKRISNCVADYSVLYNSTSFSTFQCYGVSST<br>KLNDLCFTNVYADSFVVRGDEVRQIAPGQTGVIADYNYKLPDDFTGCVLAWNSR<br>NQDASTSGNFNYYYRIWRSEKLRPFERDIAHYDYQVGTQFKSSLKNYGFYSSA<br>GDSHQPYRVVVLSFELLNAPATVCGPKQST |
| hCoV-<br>19/bat/Yunnan/RsY<br>N04/2020\|EPI_ISL_<br>1699444\|2020-04-18<br>(RsYN04) | NITNLCPFSQVFNATRFPSVYAWTRERISNCIADYSVLYNSTSFSTFRCYGVSPT<br>KLNDLCFSNVYADSMVVRGDEVRQIAPSQTGVIADYNYKLPDDFTGCVIAWNSK<br>AKDENGQYFYRLFRKSKLLPFQRDVSNVTYGSGKNDGCNPSEADCYWPLLKY<br>GFTSSVSQDYQPYRVVVLSFELLNAPATVCGPKRST |
| hCoV-<br>19/bat/Cambodia/R<br>ShSTT182/2010\|EPI<br>_ISL_852604\|2010-<br>12-06<br>(RShSTT182) | NITNLCPFGEVFNATTFASVYAWNRRRISNCVADYSVLYNTTSFSTFKCYGVSPT<br>KLNDLCFTNVYADSFVVRGDEVRQIAPGQTGKIADYNYKLPDDFMGCVIAWNSI<br>SLDAGGSYYYRLFRKSVLKPFERDISTQLYQAGDKPCSVEGPDCYYPLQSYYFQ<br>STNGVGYQPYRVVVLSFELLNAPATVCGPKKST |
| SARS CoV-2<br>Omicron_BA.5_BA.2<br>(BA.5) | RVQPTESIVRFPNITNLCPFDEVFNATRFASVYAWNRKRISNCVADYSVLYNFAP<br>FFAFKCYGVSPTKLNDLCFTNVYADSFVIRGNEVSQIAPGQTGNIADYNYKLPDD<br>FTGCVIAWNSNKLDSKVGGNYNYRYRLFRKSNLKPFERDISTEIYQAGNKPCNG<br>VAGVNCYFPLQSYGFRPTYGVGHQPYRVVVLSFELLHAPATVCGPKKSTNLVK<br>NKCVNF |
| SARS CoV-2<br>Omicron_BA.2.12.1<br>__BA.2<br>(BA.2.12.1) | RVQPTESIVRFPNITNLCPFDEVFNATRFASVYAWNRKRISNCVADYSVLYNFAP<br>FFAFKCYGVSPTKLNDLCFTNVYADSFVIRGNEVSQIAPGQTGNIADYNYKLPDD<br>FTGCVIAWNSNKLDSKVGGNYNYQYRLFRKSNLKPFERDISTEIYQAGNKPCNG<br>VAGFNCYFPLRSYGFRPTYGVGHQPYRVVVLSFELLHAPATVCGPKKSTNLVK<br>NKCVNF |
| SARS CoV-2<br>Kappa_B.1.617.1<br>(Kappa) | RVQPTESIVRFPNITNLCPFGEVFNATRFASVYAWNRKRISNCVADYSVLYNSAS<br>FSTFKCYGVSPTKLNDLCFTNVYADSFVIRGDEVRQIAPGQTGKIADYNYKLPDD<br>FTGCVIAWNSNNLDSKVGGNYNYRYRLFRKSNLKPFERDISTEIYQAGSTPCNG<br>VQGFNCYFPLQSYGFQPTNGVGYQPYRVVVLSFELLHAPATVCGPKKSTNLVK<br>NKCVNF |
| SARS CoV-2<br>Mu_B.1.621<br>(Mu) | RVQPTESIVRFPNITNLCPFGEVFNATKFASVYAWNRKRISNCVADYSVLYNSAS<br>FSTFKCYGVSPTKLNDLCFTNVYADSFVIRGDEVRQIAPGQTGKIADYNYKLPDD<br>FTGCVIAWNSNNLDSKVGGNYNYLYRLFRKSNLKPFERDISTEIYQAGSTPCNG<br>VKGFNCYFPLQSYGFQPTYGVGYQPYRVVVLSFELLHAPATVCGPKKSTNLVK<br>NKCVNF |
| SARS CoV-2<br>Lambda_C.37<br>(Lambda) | RVQPTESIVRFPNITNLCPFGEVFNATRFASVYAWNRKRISNCVADYSVLYNSAS<br>FSTFKCYGVSPTKLNDLCFTNVYADSFVIRGDEVRQIAPGQTGKIADYNYKLPDD<br>FTGCVIAWNSNNLDSKVGGNYNYQYRLFRKSNLKPFERDISTEIYQAGSTPCNG<br>VEGFNCYSPLQSYGFQPTNGVGYQPYRVVVLSFELLHAPATVCGPKKSTNLVK<br>NKCVNF |
| SARS CoV-2<br>Gamma_P.1<br>(Gamma) | RVQPTESIVRFPNITNLCPFGEVFNATRFASVYAWNRKRISNCVADYSVLYNSAS<br>FSTFKCYGVSPTKLNDLCFTNVYADSFVIRGDEVRQIAPGQTGTIADYNYKLPDD<br>FTGCVIAWNSNNLDSKVGGNYNYLYRLFRKSNLKPFERDISTEIYQAGSTPCNG<br>VKGFNCYFPLQSYGFQPTYGVGYQPYRVVVLSFELLHAPATVCGPKKSTNLVK<br>NKCVNF |
| SARS CoV-2<br>Beta_B.1.351<br>(Beta) | RVQPTESIVRFPNITNLCPFGEVFNATRFASVYAWNRKRISNCVADYSVLYNSAS<br>FSTFKCYGVSPTKLNDLCFTNVYADSFVIRGDEVRQIAPGQTGNIADYNYKLPDD<br>FTGCVIAWNSNNLDSKVGGNYNYLYRLFRKSNLKPFERDISTEIYQAGSTPCNG<br>VKGFNCYFPLQSYGFQPTYGVGYQPYRVVVLSFELLHAPATVCGPKKSTNLVK<br>NKCVNF |
| SARS CoV-2<br>Alpha_B.1.1.7<br>(Alpha) | RVQPTESIVRFPNITNLCPFGEVFNATRFASVYAWNRKRISNCVADYSVLYNSAS<br>FSTFKCYGVSPTKLNDLCFTNVYADSFVIRGDEVRQIAPGQTGKIADYNYKLPDD<br>FTGCVIAWNSNNLDSKVGGNYNYLYRLFRKSNLKPFERDISTEIYQAGSTPCNG |

| | |
|---|---|
| | VEGFNCYFPLQSYGFQPTYGVGYQPYRVVVLSFELLHAPATVCGPKKSTNLVK<br>NKCVNF |
| SARS CoV-2<br>BA.2.75<br>(BA.2.75) | RVQPTESIVRFPNITNLCPFHEVFNATRFASVYAWNRKRISNCVADYSVLYNFAP<br>FFAFKCYGVSPTKLNDLCFTNVYADSFVIRGNEVSQIAPGQTGNIADYNYKLPDD<br>FTGCVIAWNSNKLDSKVGGNYNYLYRLFRKSNLKPFERDISTEIYQAGNKPCNG<br>VAGFNCYFPLQSYGFRPTYGVGHQPYRVVVLSFELLHAPATVCGPKKSTNLVK<br>NKCVNF |
| SARS CoV-2<br>BA.4.6/BA.5.2.6<br>(BA.4.6) | RVQPTESIVRFPNITNLCPFDEVFNATTFASVYAWNRKRISNCVADYSVLYNFAP<br>FFAFKCYGVSPTKLNDLCFTNVYADSFVIRGNEVSQIAPGQTGNIADYNYKLPDD<br>FTGCVIAWNSNKLDSKVGGNYNYRYRLFRKSNLKPFERDISTEIYQAGNKPCNG<br>VAGVNCYFPLQSYGFRPTYGVGHQPYRVVVLSFELLHAPATVCGPKKSTNLVK<br>NKCVNF |
| SARS CoV-2<br>BQ.1.1/XBB_L452R<br>_R493Q_R346T_F4<br>86V_N658S_N460K<br>_K444T_F490S<br>(BQ.1.1) | RVQPTESIVRFPNITNLCPFDEVFNATTFASVYAWNRKRISNCVADYSVLYNFAP<br>FFAFKCYGVSPTKLNDLCFTNVYADSFVIRGNEVSQIAPGQTGNIADYNYKLPDD<br>FTGCVIAWNSNKLDSTVGGNYNYRYRLFRKSKLKPFERDISTEIYQAGNKPCNG<br>VAGVNCYFPLQSYGFRPTYGVGHQPYRVVVLSFELLHAPATVCGPKKSTNLVK<br>NKCVNF |
| SARS CoV-2<br>BA.2.3.20<br>(BA.2.3.20) | RVQPTESIVRFPNITNLCPFDEVFNATRFASVYAWNRKRISNCVADYSVLYNFAP<br>FFAFKCYGVSPTKLNDLCFTNVYADSFVIRGNEVSQIAPGQTGNIADYNYKLPDD<br>FTGCVIAWNSNKLDSRVGGNYDYMYRLFRKSKLKPFERDISTEIYQAGNKPCNG<br>VRGFNCYFPLQSYGFRPTYGVGHQPYRVVVLSFELLHAPATVCGPKKSTNLVK<br>NKCVNF |
| SARS CoV-2 XBB<br>(XBB) | RVQPTESIVRFPNITNLCPFHEVFNATTFASVYAWNRKRISNCVADYSVIYNFAP<br>FFAFKCYGVSPTKLNDLCFTNVYADSFVIRGNEVSQIAPGQTGNIADYNYKLPDD<br>FTGCVIAWNSNKLDSKPSGNYNYLYRLFRKSKLKPFERDISTEIYQAGNKPCNG<br>VAGSNCYSPLQSYGFRPTYGVGHQPYRVVVLSFELLHAPATVCGPKKSTNLVK<br>NKCVNF |
| SARS CoV-2 BQ.1<br>(BQ.1) | RVQPTESIVRFPNITNLCPFDEVFNATRFASVYAWNRKRISNCVADYSVLYNFAP<br>FFAFKCYGVSPTKLNDLCFTNVYADSFVIRGNEVSQIAPGQTGNIADYNYKLPDD<br>FTGCVIAWNSNKLDSTVGGNYNYRYRLFRKSKLKPFERDISTEIYQAGNKPCNG<br>VAGVNCYFPLQSYGFRPTYGVGHQPYRVVVLSFELLHAPATVCGPKKSTNLVK<br>NKCVNF |
| SARS CoV-2 BN.1<br>(BN.1) | RVQPTESIVRFPNITNLCPFHEVFNATTFASVYAWNRTRISNCVADYSVLYNFAP<br>FFAFKCYGVSPTKLNDLCFTNVYADSFVIRGNEVSQIAPGQTGNIADYNYKLPDD<br>FTGCVIAWNSNKLDSKVSGNYNYLYRLFRKSKLKPFERDISTEIYQAGNKPCNG<br>VAGFNCYSPLQSYGFRPTYGVGHQPYRVVVLSFELLHAPATVCGPKKSTNLVK<br>NKCVNF |
| SARS CoV-2 CH.1.1<br>(CH.1.1) | RVQPTESIVRFPNITNLCPFHEVFNATTFASVYAWNRKRISNCVADYSVLYNFAP<br>FFAFKCYGVSPTKLNDLCFTNVYADSFVIRGNEVSQIAPGQTGNIADYNYKLPDD<br>FTGCVIAWNSNKLDSTVSGNYNYRYRLFRKSKLKPFERDISTEIYQAGNKPCNG<br>VAGSNCYFPLQSYGFRPTYGVGHQPYRVVVLSFELLHAPATVCGPKKSTNLVK<br>NKCVNF |
| SARS CoV-2<br>XBB.1.5<br>(XBB.1.5) | RVQPTESIVRFPNITNLCPFHEVFNATTFASVYAWNRKRISNCVADYSVIYNFAP<br>FFAFKCYGVSPTKLNDLCFTNVYADSFVIRGNEVSQIAPGQTGNIADYNYKLPDD<br>FTGCVIAWNSNKLDSKPSGNYNYLYRLFRKSKLKPFERDISTEIYQAGNKPCNG<br>VAGPNCYSPLQSYGFRPTYGVGHQPYRVVVLSFELLHAPATVCGPKKSTNLVK<br>NKCVNF |

**Table S4: Cryo-EM model building and refinement statistics (model shown in Figure 4).**

| Model | | SARS-CoV-2 : PRO-37587 |
|---|---|---|
| # of chains | | 9 |
| # of residues | | 4,332 |
| # of carbohydrates | | 39 |
| MolProbity score | | 1.63 |
| All-atom clash score | | 4.63 |
| Rotamer outliers (%) | | 1.76 |
| $C_\beta$ outliers | | 0.00 |
| Ramachandran Plot Values | Favored (%) | 96.69 |
| | Allowed (%) | 3.31 |
| | Outlier (%) | 0.00 |
| R.M.S Deviations | Bond lengths (Å) | 0.002 |
| | Bond angles (°) | 0.456 |
| Correlation Coefficient | | 0.82 |

**Table S5: List of contact residues on PRO-37587 to SARS-CoV-2 Omicron BA.1 RBD based on a 0.45 nm distance cutoff between heavy atoms. Parentheses indicate number of observed contacts.**

| Contacting molecule | PRO-37587 (light chain) contact residues | PRO-37587 (heavy chain) contact residues |
|---|---|---|
| RBD | Ala31 (2)<br>Gly32 (1)<br>Tyr33 (1)<br>Arg34 (1)<br>Tyr93 (2)<br>Ser95 (4)<br>Ser96 (1)<br>Leu97 (4)<br>Phe98 (1)<br>Asp99 (3)<br>Pro100 (1) | Ser29 (1)<br>Tyr32 (1)<br>Arg50 (1)<br>Ile52 (2)<br>Leu54 (1)<br>Ser55 (1)<br>Tyr57 (3)<br>Asn59 (1)<br>Asp103 (2)<br>Tyr104 (1)<br>Tyr105 (1)<br>Gly106 (7)<br>Trp107 (3)<br>Gly108 (5)<br>Asp109 (3)<br>Asp110 (4) |

**Table S6: List of contact residues on SARS-CoV-2 Omicron BA.1 RBD to PRO-37587 based on a 0.45 nm distance cutoff between heavy atoms. Parentheses indicate number of observed contacts.**

| Contacting molecule | RBD contact residues |
|---|---|
| PRO-37587 (light chain) | Gly404 (4)<br>Asp405 (3)<br>Arg408 (1)<br>Thr500 (1)<br>Tyr501 (3)<br>Gly502 (1)<br>Val503 (1)<br>Gly504 (1)<br>Gln506 (6) |
| PRO-37587 (heavy chain) | Tyr369 (1)<br>Asn370 (2)<br>Ala372 (2)<br>Pro373 (4)<br>Phe374 (1)<br>Phe375 (7)<br>Thr376 (3)<br>Phe377 (1)<br>Lys378 (3)<br>Cys379 (2)<br>Tyr380 (3)<br>Val382 (2)<br>Ser383 (3)<br>Pro384 (2)<br>Thr385 (1) |
